# Identifying Evolutionary Relatedness Effects on Diversification from Phylogenies using Neural Networks

**DOI:** 10.64898/2026.01.12.699140

**Authors:** Tianjian Qin, Koen van Benthem, Luis Valente, Rampal Etienne

**Author notes:** Corresponding author. *Email addresses:* (Tianjian Qin), (Koen van Benthem), (Luis Valente), (Rampal Etienne). These authors contributed equally as joint senior authors (Luis Valente, Rampal Etienne).

## Abstract

Reconstructing the forces that shaped macroevolutionary histories from extant phylogenies is fundamentally challenging: richly parameterized diversification models are often only weakly identifiable; different evolutionary mechanisms can yield nearly indistinguishable tree shapes. Here we use a model with evolutionary relatedness dependence to evaluate how much information about such forces can be recovered from simulated trees. We train graph neural networks and long short-term memory classifiers to distinguish three scenarios of feedback of diversity on diversification: effect of phylogenetic diversity (total branch length), evolutionary distinctiveness (average phylogenetic distance of a species to all other species in a clade), and nearest-neighbor distance (phylogenetic distance to the mostly closely related species). We also train a suite of regression networks to estimate the underlying diversification parameters. We then analyze classification performance, calibration of predicted class probabilities, regression errors, and their dependence on tree size and on the strength and sign of richness and relatedness effects. Across network architectures and complexity levels, scenario classification is only moderately accurate and strongly asymmetric as revealed by the confusion matrix. Trees generated under an effect of nearest-neighbor distance on diversification tend to be correctly classified, whereas those with an effect of evolutionary distinctiveness are frequently misclassified. Regression networks systematically shrink predictions toward the empirical mean, especially for complex models, suggesting broad regions of parameter space with low identifiability. Strong global dependence of diversification rates on diversity further erodes recoverability by driving large variations in tree size that mask the subtler signatures of relatedness effects. In contrast, sufficiently strong speciation-relatedness effects can carve out narrow regions of parameter space in which scenarios and parameters become practically recoverable. Together, our results provide a map of when neural networks can and cannot infer diversification mechanisms from extant trees under our evolutionary relatedness dependence model, and they underscore the need for additional data or constraints when using flexible diversification models for macroevolutionary inference.

## 1. Introduction

Time-calibrated phylogenies are now central tools for studying macroevolutionary dynamics. From such trees, we seek to reconstruct how speciation and extinction rates varied through time and across lineages, and how ecological limits and biotic interactions have shaped present-day diversity (Nee et al., 1992; Quental and Marshall, 2010). Early work typically assumed constant diversification rates, which implies exponential lineage accumulation (Nee et al., 1994a; Kubo and Iwasa, 1995). However, reconstructed phylogenies for many clades show pronounced slowdowns in lineages-through-time curves, inconsistent with simple constant-rate models and motivating a broad family of more flexible diversification models, including time-dependent, diversity-dependent, and protracted speciation models (Phillimore and Price, 2008; Moen and Morlon, 2014; Etienne et al., 2012; Etienne and Rosindell, 2012; Rosindell et al., 2010). A common mechanistic theme in these models is that speciation rates decline as clades fill ecological or niche space, with species richness acting as a proxy for ecological limits (Srivastava et al., 2012; Kondratyeva et al., 2019).

At the same time, there is growing recognition that extant phylogenies carry only limited information about past diversification histories. Nee et al. (1994b) already showed that a constant-rate birth-death model is equivalent to a pure birth model with temporally declining speciation rate. This property allowed them to compute the probability of a phylogeny given the constant-rate birth-death model, but also implies that there is inherent indistinguishability of these models based on phylogenetic information alone. This result was generalized by Louca and Pennell (2020), who showed that, under general birth-death processes, an infinite number of distinct speciation-extinction trajectories can produce exactly the same distribution of extant time trees, forming large “congruence classes” of observationally equivalent models. Even for more constrained model families, branching patterns alone often cannot discriminate between alternative mechanisms. For instance, Pannetier et al. (2021) demonstrated that diversity-dependent and purely time-dependent diversification models that share the same expected diversity-through-time curve are essentially indistinguishable from extant trees using standard likelihood methods. These results highlight that increasing model flexibility does not automatically yield more informative inferences.

The recently-developed `eve` model of Qin et al. (2025b) augments diversity-dependent diversification by allowing speciation and extinction rates to depend linearly on both species richness and a measure of evolutionary relatedness (ER) for each lineage. The rationale for this is that ER may have an effect in diversification rates that is independent from species diversity, for instance, if closely related species are more likely to compete for niches. ER can be defined at different phylogenetic scales, for example using clade-wide phylogenetic diversity (PD, total branch length in the tree) or more localized, lineage-specific metrics such as evolutionary distinctiveness (ED, average phylogenetic distance to all other species in the clade) or nearest-neighbor distance (NND, phylogenetic distance to the most closely related species). Depending on the signs and magnitudes of the richness and ER effects, `eve` can generate a wide range of branching patterns and tree shapes, and previous work has suggested that the PD, ED and NND variants can produce near identical tree-shape signatures for moderate effect sizes, in terms of classic phylogenetic summary statistics.

The complexity and state dependence of `eve` make likelihood-based inference of parameters of the model analytically intractable, encouraging a shift toward simulation-based approaches. Neural networks and other deep-learning methods have recently been proposed to estimate diversification parameters and classify models from phylogenetic trees (Lajaaiti et al., 2023; Voznica et al., 2022; Lambert et al., 2023; Qin et al., 2025a). These methods can learn from graph representations of trees, branching times and summary statistics, and for relatively simple birth-death or diversity-dependent models they can achieve performance comparable to, or potentially exceeding, maximum likelihood estimation. However, when applied to models already known to suffer from non-identifiability, such as protracted speciation (Etienne et al., 2014), neural networks tend to converge to conservative predictions that reflect the empirical mean of the training distribution rather than the true parameters. In such cases, the limiting factor appears to be the information content of the trees, even after extensive sweeps over neural network architectures, depths, and hyperparameter settings, rather than the expressive power of the estimator (Qin et al., 2025a). This suggests that models as flexible as diversity-dependent diversification or protracted speciation may already lie close to the practical limits of what can be identified from extant trees, regardless of whether one uses maximum likelihood or neural networks, and thus there is little reason to expect neural networks to reliably recover all parameters of an even more complex model such as eve.

In this study we use neural networks as means to explore the putative limits of information content of phylogenies regarding diversification processes in complex models. We simulate large collections of extant trees under the three ER scenarios (PD, ED, NND), three levels of model complexity (2, 4 or 6 free parameters), and a broad range of effect sizes for the effects of species richness and evolutionary relatedness. We then train graph neural networks (GNNs) and long short-term memory (LSTM) networks as classifiers to distinguish PD, ED and NND, and as regressors (in an ensemble GNN+LSTM architecture as developed by Qin et al. (2025a)) to recover the underlying parameters from tree representations and branching times. Performance is evaluated as a function of tree size, true parameter values, model complexity and regressor/classifier misspecification, using simulated tree datasets.

By asking when these neural network learners succeed or fail, we obtain a map of the practical recoverability of `eve` under a range of scenarios. High classification accuracy or precise parameter recovery marks regions of parameter space where the three ER scenarios generate distinct, possibly information-rich tree patterns. Systematic misclassification and regression predictions collapsing toward the conditional mean correlate to non-recoverability, suggesting that different parameter combinations or ER mechanisms give rise to overlapping tree shapes that cannot be distinguished. Small trees provide fewer branching events and shorter histories, and thus are expected to fall into low-information, more non-identifiable settings, analogous to sample size in time-series hidden Markov models (Cole, 2019).

Our goals are therefore twofold. First, we quantify how tree size (number of nodes of the trees), effect size (magnitude of the values of parameters controlling the effects of species richness and evolutionary relatedness on diversification) and model complexity (number of effective parameters) interact to determine the recoverability of `eve` parameters and the discriminability of PD, ED and NND from extant trees. Second, we interpret the characteristic failure modes of our networks, e.g., conservative regression, confusion between ER scenarios, and differing sensitivity to global (PD) versus local (ED/NND) forces, to gain insight into the expected behaviors for parameter estimation using deep learning on complex diversification models.

## 2. Materials and Methods

### 2.1. Software and Hardware

We used `PyTorch 1.12.1` (Imambi et al., 2021), `PyTorch Geometric 2.3.1` (Fey and Lenssen, 2019), `Python 3.7.1` and `CUDA 12.2.2` (Luebke, 2008) for the neural networks and `R 4.2.1` (R Core Team, 2025) for data processing, simulation and visualization. All the computationally heavy tasks were performed on the Hábrók high-performance computing cluster of the University of Groningen. Our neural networks were trained, optimized and evaluated on the NVIDIA A100 and V100 tensor core GPUs of the Hábrók cluster.

### 2.2. Simulation Approaches

We used the `evesim` R package (Qin et al., 2025b) — an efficient C++ implementation of the `eve` model — to simulate phylogenetic trees. We kept only extant lineages for each of the trees. The parameter settings used for simulation were chosen to limit the maximum number of extant lineages to maintain a manageable memory and computational demand for both simulation and neural network training. We filtered out simulated trees containing more than 1500 lineages (terminal tips), due to limited available hardware resources. Large trees were rare in our settings, typically less than 10 trees were removed per complete run. The sizes of the simulated trees may vary per each complete run. We recorded how many trees were removed in this way.

In the `eve` diversification model, there are three evolutionary scenarios, each of which represents a unique relatedness metric affecting macroevolutionary trajectories: PD (Phylogenetic Diversity), ED (Evolutionary Distinctiveness) and NND (Nearest Neighbor Distance). Each of the evolutionary scenarios are characterized by six parameters that shape the phylogenies: intrinsic speciation rate rate *λ*_0_; intrinsic extinction rate rate *µ*_0_; effect size of species richness on speciation *β_N_*; effect size of evolutionary relatedness on speciation *β_Φ_*; effect size of species richness on extinction *γ_N_*; effect size of evolutionary relatedness on extinction *γ_Φ_*.

Let *N_t_* be species richness at time *t* and *Φ_i,t_* the evolutionary relatedness score for lineage *i* at time *t* (we write *Φ_t_* when it does not depend on lineage). Then we have

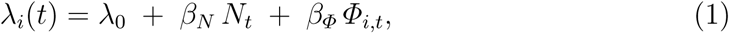

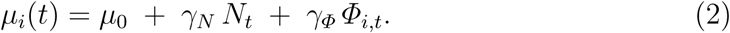

Under the PD scenario, *Φ_i,t_*≡ *Φ_t_* for all lineages *i*. Consequently, *λ_i_*(*t*) = *λ_j_*(*t*) and *µ_i_*(*t*) = *µ_j_*(*t*) for all *i, j*. Under ED and NND scenarios, *Φ_i,t_* depends on *i*, so *λ_i_*(*t*) and *µ_i_*(*t*) may differ across lineages at the same *t*. See Qin et al. (2025b) for more details.

By changing the number of parameters involved in simulation, we generated datasets of three different levels of model complexity for each of the three evolutionary scenarios. The low-complexity datasets were the results of simulation using *λ*_0_ and *µ*_0_; the medium-complexity datasets were the results of simulation using two more parameters *β_N_* and *β_Φ_*; the high-complexity datasets were the results of simulation using two additional parameters *γ_N_*and *γ_Φ_*. This staged parameterization helps disentangle the contributions of time-varying speciation and time-varying extinction by progressively introducing separate sets of parameters that modulate each process and comparing their incremental effects on the simulated trees.

Per complexity level and per scenario, we randomly sampled the parameters from uniform distributions. The upper bound for the extinction rates were proportionally dependent on the drawn speciation rate to avoid cases where extinction rates could be larger than speciation rates, because in such cases the whole tree likely goes extinct. We simulated 100,000 trees (before size-filtering) per scenario per complexity and split later for neural network training (90%) and validation (10%). For each scenario-complexity combination, we additionally simulated 10,000 trees for the purpose of performance test. The number of simulated trees is bounded by time limits and available hardware resources of the computing cluster. See Table 1 for detailed parameter settings and Figure E.25 for the distribution of parameters of successfully generated and retained trees.

**Table 1:**
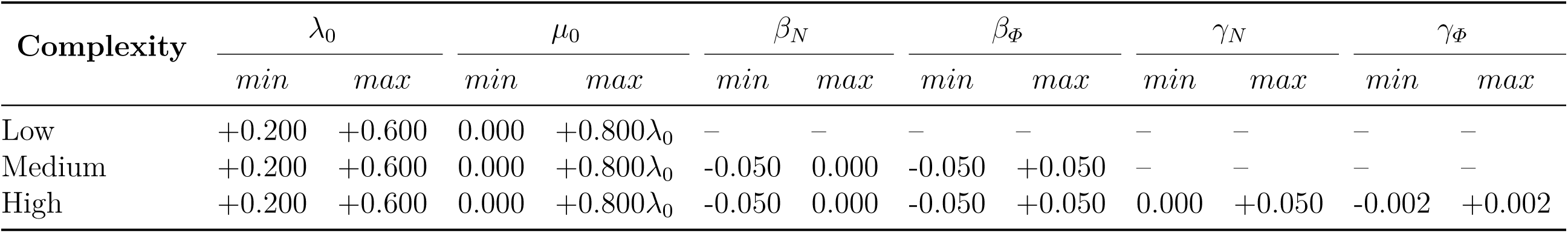
Parameter settings for the simulated tree datasets using different model settings. The “Complexity” column specifies which level of model complexity the datasets belong to, with regard to the number of parameters. Each complexity level is crossed with all three evolutionary scenarios as described in the methods. All simulated trees have identical crown age of 10 time units. For each complexity level and evolutionary scenario combination, 100,000 trees were simulated. The sub-columns under each of the parameters specify the lower (*min*) and the upper (*max*) bounds of the parameter space; all parameters are sampled from *U* (*min, max*), where *U* denotes uniform distribution. *λ*_0_: intrinsic speciation rate rate; *µ*_0_: intrinsic extinction rate rate; *β_N_* : effect size of species richness on speciation; *β_Φ_*: effect size of evolutionary relatedness on speciation; *γ_N_* : effect size of species richness on extinction; *γ_Φ_*: effect size of evolutionary relatedness on extinction. For each parameter, the symbol “–” indicates that this parameter is always 0, thus disabling its impact on the simulation.

We assumed an identical crown age of 10 time units (*t* = 10) for all phylogenies. The choice of crown age is arbitrary because we can rescale the crown age arbitrarily, as long as we rescale the generating parameters accordingly. See also Figure 1 for an illustration of simulation settings.

**Figure 1:**
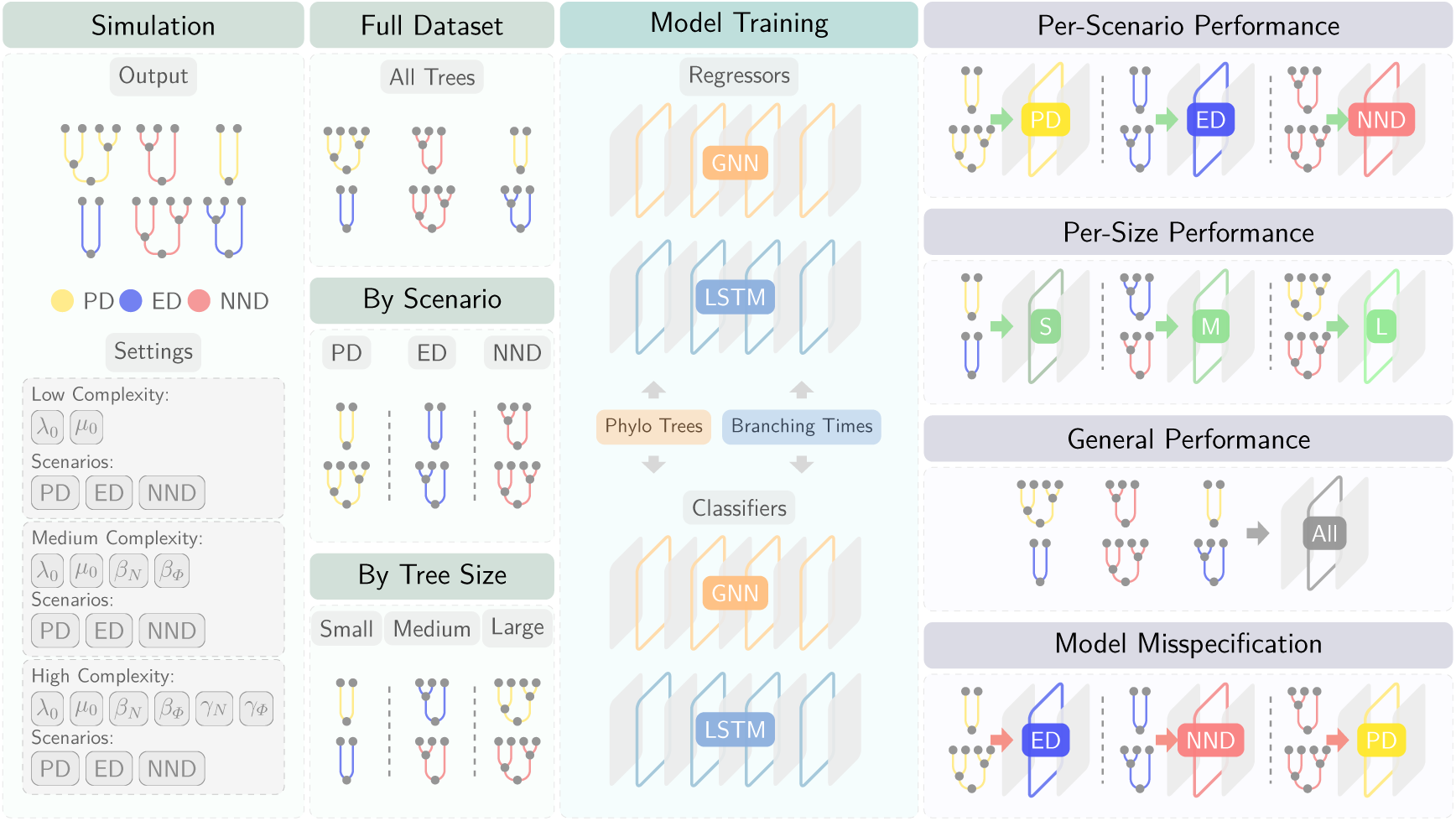
General overview of the workflow: phylogenetic trees are simulated under PD (phylogenetic diversity), ED (evolutionary distinctiveness), and NND (nearest-neighbor distance) scenarios across low/medium/high complexity parameterizations, then assembled into the full dataset, or stratified into per-scenario and per-tree-size datasets. These datasets were used to train neural network classifiers/regressors using information such as tree representations and branching times. Neural networks were evaluated per scenario, per size, overall, and under model misspecification.

### 2.3. Classification

#### 2.3.1. Network Architecture and Training

The `eve` simulations involve three evolutionary scenarios. To investigate whether neural networks can identify these scenarios, we trained graph neural network (GNN) and long short-term memory (LSTM) classifiers independently on the simulated datasets. In short, in our settings, GNNs learn representations by iterative neighborhood aggregation over the graph topology (Kipf and Welling, 2016), whereas LSTMs are gated recurrent networks that summarize sequential inputs via a memory cell (Sak et al., 2014). For the GNN, the full phylogeny was represented as a graph, with nodes corresponding to taxa and edges representing evolutionary relationships. For the LSTM, we transformed the phylogeny into a sequence of branching times, treating these as time-series data. Input data were prepared using `PyTorch` utilities and our custom functions in the codebase `eveGNN` (Qin, 2023).

Detailed base architectures and hyperparameter settings of these neural networks are described in Qin et al. (2025a). This design has shown good regression performance on phylogenetic trees (Qin et al., 2025a). Here, instead of outputting parameter estimates, for each input phylogeny a classifier produces a three-class prediction, which we write as a probability vector

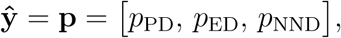

with *p*_PD_ + *p*_ED_ + *p*_NND_ = 1. Each component represents the predicted probability that the input tree was generated under the corresponding scenario.

During training, we optimized the classifiers using cross-entropy loss, which directly compares the predicted probabilities **ŷ** to the true class labels **y**. Because the classification task requires phylogenies from all three evolutionary scenarios, the classifiers were trained on the full dataset that pools trees generated under PD, ED, and NND.

For the GNN classifiers, two GNN-specific loss terms designed to facilitate graph-structured learning were added to the cross-entropy loss term. The full specification of the classification loss is given in Appendix A. For optimization, we employed the AdamW (adaptive moment estimation with decoupled weight decay) algorithm (Loshchilov and Hutter, 2017) to iteratively minimize the total loss and update network weights.

To balance computational efficiency and GPU memory usage, training was conducted using mini-batches of size 64, where each mini-batch contained data from 64 simulated phylogenies. During training, model checkpoints were saved at multiple epochs. We compared training and validation losses across epochs and selected the checkpoint that achieved stable generalization performance, balancing underfitting and overfitting.

#### 2.3.2. Performance Analysis

We evaluated the trained classifiers using confusion matrices, standard classification metrics, distributional comparisons of generating parameters, empirical accuracy surfaces, and probability calibration diagnostics. Here we illustrate the main ideas, precise definitions and formulas for all classification performance measures are given in Appendix H.

To summarize scenario-level performance, we computed confusion matrices for each classifier and for different groupings of the test data. For every tree, the predicted class was taken to be the scenario with the largest posterior probability among {PD, ED, NND}, ties (equal probabilities) were broken at random but were rare in practice. The resulting 3 × 3 confusion matrices have rows corresponding to the true scenario and columns to the predicted scenario, so that each entry *C_rc_* counts how many trees with true class *r* were predicted as class *c*. For visualization, we row-normalized each confusion matrix so that the entries in each row *r* sum to one, i.e. *C̃_rc_* = *C_rc_*/ Σ_*c′*_ *C_rc′_*. The diagonal elements then correspond directly to the class-wise recall (true positive rate) for each scenario, whereas the off-diagonal elements give the conditional misclassification rates. We constructed two sets of contrasts based on these matrices: (i) a comparison between GNN and LSTM architectures, and (ii) a comparison between medium- and high-complexity versions of the `eve` model (four versus six free parameters), fixing the architecture to GNN. For each contrast and each cell (*r, c*) we computed the change in row-normalized recall between methods *A* and *B*,

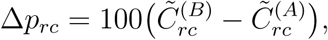

In addition to confusion matrices, we computed overall and per-scenario accuracy, precision, recall (sensitivity), and F1-score for each classifier. The exact formulas and notation used are provided in Appendix H. To study how classification performance varies across parameter space, we also sliced the results into consecutive ranges based on the true phylogenetic parameters and the simulated tree sizes, as detailed in Appendix J. These ranges allowed us to examine how accuracy and other metrics depend on effect sizes and tree size.

To further investigate how classification success depends on the true diversification parameters, we treated the correctness of the prediction for each tree as a binary response (`correct` ∈ {0, 1}) and compared the distributions of the generating parameters between correctly and incorrectly classified trees. First, we summarized the marginal distributions of each parameter {*λ*_0_*, µ*_0_*, β_N_, β_Φ_, γ_N_, γ_Φ_*} and of tree size (|𝒯|) using ridge–line density plots (i.e., vertically stacked and slightly overlapping density curves on a shared horizontal axis), stratified by architecture (GNN vs. LSTM) and true scenario (PD, ED, NND). For each scenario-architecture-parameter combination we carried out a two-sample Kolmogorov-Smirnov (KS) test comparing the parameter values for correctly versus incorrectly classified trees, and adjusted the resulting *p*-values across tests by the Benjamini-Hochberg procedure. The KS statistic provides a nonparametric measure of the maximum discrepancy between empirical cumulative distributions and is sensitive to differences of distributions in both location and shape.

Second, we quantified a global measure of dependence between correctness and the full parameter vector (*λ*_0_*, µ*_0_*, β_N_, β_Φ_, γ_N_, γ_Φ_,* |𝒯|*, λ*_0_ − *µ*_0_) while controlling for scenario and architecture. We used partial distance correlation, with the true class and model encoded as conditioning variables. This test evaluates the null hypothesis that, given scenario and architecture, the parameter vector is independent of the correctness indicator. The resulting *p*-value is reported as an overall measure of residual parameter dependence.

To evaluate how well the class probabilities output by our classifiers reflect true predictive uncertainty, we performed a calibration analysis based on reliability diagrams and expected calibration error (ECE). For each phylogeny in the test sets we recorded the predicted scenario (*ŷ* ∈ {PD, ED, NND}), the corresponding maximum predicted probability *p̂* = max{*p̂*_PD_*, p̂*_ED_*, p̂*_NND_}, and an indicator of cor-rectness 𝕀(*y* = *y*), where *y* denotes the true scenario. Following standard practice, we partitioned the confidence interval [0, 1] into *M* = 10 equally wide bins *B_m_* = [(*m* − 1)*/M, m/M*) and, for every analysis stratum, computed for each bin the average confidence

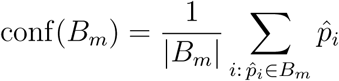

and the empirical accuracy

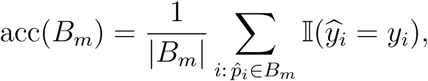

where |*B_m_*| is the number of test trees whose confidence falls in bin *B_m_*. To summarize miscalibration with a single scalar, we used the expected calibration error

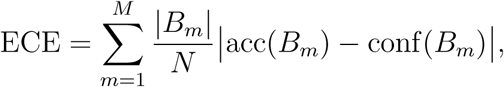

where *N* is the total number of test trees in the stratum; lower ECE indicates better agreement between predicted confidences and observed accuracies. We computed reliability curves and ECE in several complementary strata (overall, per scenario, per tree size group, and across model complexities) and also after partitioning trees into ranges of *β_N_* and *β_Φ_*.

### 2.4. Regression

#### 2.4.1. Network Architecture and Training

Like classification, we also used GNN and LSTM for the regression tasks. Additionally, we combined GNN and LSTM using a sequential “boosting” method to leverage all available information and reduce prediction error. In this approach, the GNN first provides an initial parameter estimate; the LSTM then predicts the residual errors of the GNN, and the final estimate is obtained by adding this correction. Details of the neural network regressors and the “boosting” method are described and discussed in Qin et al. (2025a). For each level of complexity, the vector of predicted variables 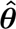 comprises all nonzero parameters used in simulation in the model (two, four, or six for the low-, medium-, and high-complexity settings, respectively, as described above).

For the regression analyses, the simulated tree datasets were reorganized into subsets tailored to different questions. Across all complexity levels, we partitioned the data according to (i) evolutionary scenario in the `eve` model, yielding three subsets that contain only phylogenies generated under PD, ED, or NND, respectively, for studying per-scenario and regressor-misspecification performance, and (ii) tree size (i.e., the total number of nodes), yielding three consecutive ranges: small trees (*n* ∈ [0, 200]), medium trees (*n* ∈ [201, 500]), and large trees (*n >* 500), for studying size-specific performance. In addition to these subsets, we retained the full dataset that contains all simulated phylogenies for assessing overall performance (see also Figure 1).

For the LSTM regressors, we used the Huber loss (Huber, 1992) to quantify prediction error during training. For the GNN regressors and the boosting method, two GNN-specific loss terms designed to facilitate graph-structured learning were added to the Huber loss term. The full specification of the regression loss is given in Appendix A. For optimization and training, similar approaches were used as in the classification tasks.

For each complexity level, the above subsetting allowed us to train regressors separately on (i) the complete training dataset, (ii) scenario-specific training datasets, and (iii) size-specific training datasets. For each neural network architecture, this yields a total of seven regression models per complexity level. The structure of the regression analyses and data partitions is summarized in Figure 1.

#### 2.4.2. Performance Analysis

Regression performance was evaluated using residual diagnostics and correlation-based measures. Here we provide a brief overview; formal definitions and equations are given in Appendix I. For each trained regressor, we computed residuals as the differences between the true parameters and the predicted parameters, 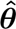 − ***θ***, and summarized these residuals as a function of (i) the true net diversity-independent part of the diversification rate *λ*_0_ − *µ*_0_, (ii) tree size (the total number of nodes, including root, internal, and tip nodes), and (iii) the four evolutionary relatedness effect sizes *β_N_*, *β_Φ_*, *γ_N_*, and *γ_Φ_*. As a reference for expected tree sizes under a simple birth-death process, we also computed the expectation and an approximate confidence interval for tree size (Appendix K). To quantify how closely the predictions follow the conditional mean of the training data, we calculated four additional metrics: the coefficient of determination, a slope difference measure, the distance correlation, and Spearman’s rank correlation. These metrics characterize, respectively, linear fit, systematic shrinkage toward the mean, overall dependence (including nonlinear structure), and monotonic association between predictions and true parameters.

The regression analyses focused on four aspects of performance: overall performance across all trees, per-size performance, per-scenario performance, and performance under regressor misspecification. Performance under regressor misspecification was assessed by applying scenario-specific regressors (trained on trees from only one scenario) to the complete testing datasets that combine trees from all three scenarios. Because the testing sets contain trees generated under all scenarios, this cross-application reveals how strongly regressor performance degrades when the assumed scenario does not match the true generative process. Our pipeline first classifies trees into scenarios and then applies a scenario-specific regressor to estimate parameters. Quantifying how sensitive the regressors are to upstream classification errors therefore provides insight into how the full pipeline may perform in practice. Figure 1 provides an overview of the analysis steps.

## 3. Results

### 3.1. Classification

Overall, the classifiers recover only a moderate amount of information about the underlying diversification scenario. Across architectures and model complexities, F1 scores and scenario-specific recalls rarely approach one and are often closer to 0.5, especially for ED trees, indicating that misclassification is common. In the following sections, we examine when misclassification becomes less common, treating this primarily as a diagnostic of practical scenario recoverability from extant trees.

#### 3.1.1. Confusion matrices

The row-normalized confusion matrices in Figure 2 show that all classifiers recover substantial information about the underlying diversification scenario. The errors are strongly structured and non-symmetric across scenarios. For both architectures, when trained on all complexities simultaneously (see the two panels in the top row of Figure 2), NND trees are easiest to classify and ED trees are consistently the most difficult. For example, under the GNN trained on all complexities (top-left panel), around 72% of NND trees are correctly identified, whereas only about 56% of PD trees and 21% of ED trees fall on the diagonal. Large proportions of misclassified ED trees are predicted as NND, potentially indicating that the ED and NND scenarios generate overlapping tree patterns. PD trees are also more frequently confused with NND, whereas the reverse error (NND mislabeled as PD) is comparatively rare.

**Figure 2:**
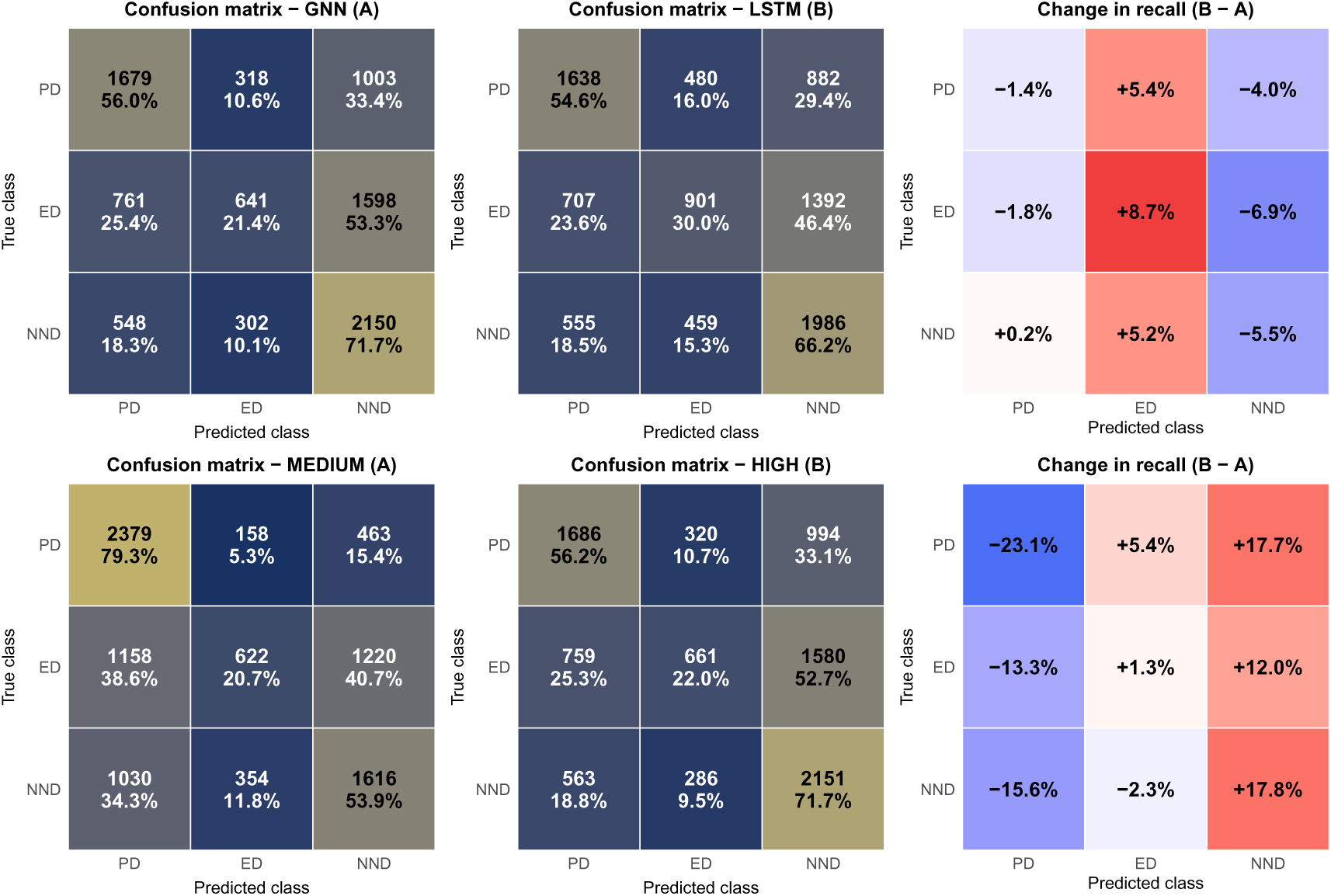
Row-normalized confusion matrices and changes in scenario-specific recall. **Top row**: comparison between classifier architectures on trees of *all complexities*. The left and middle panels show 3 × 3 confusion matrices for the GNN (A) and LSTM (B) classifiers, respectively, evaluated on the same test trees and pooled across all complexity levels (low, medium, high). Rows correspond to the true evolutionary scenario (PD, ED, NND) and columns to the predicted scenario. The y-axis order is such that the diagonal from top left to bottom right corresponds to matching true and predicted classes (PD, ED, NND). Each tile shows the absolute number of trees and the row-wise percentage of trees in that cell; colors encode the row-normalized value *C̃_rc_*, so the diagonal entries represent the per-scenario recall (true positive rate) and off-diagonal entries represent conditional misclassification rates. The right-hand panel displays the change in recall (in percentage points) between methods B and A for each cell, 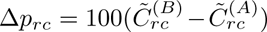. Red tiles indicate that method B predicts that combination of true and predicted class more frequently than method A, and blue tiles indicate the opposite. **Bottom row**: analogous comparison *between complexity levels* of the `eve` model. Both confusion matrices are obtained with the GNN classifier, but the left panel (A) uses only trees simulated under the medium-complexity (four-parameter) settings, whereas the middle panel (B) uses only trees from the high-complexity (six-parameter) settings. The right-hand panel again shows changes in recall 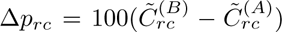 in the same format as above. Together, these panels reveal which scenarios are systematically confused with one another, how this pattern depends on architecture, and how increasing the number of free parameters in the `eve` model shifts errors towards particular predicted scenarios.

Comparing architectures on the same test set (top row in Figure 2), the LSTM trades some PD and NND recall for better separation of the ED scenario. The change-in-recall panel (top right in Figure 2) shows that, across all true scenarios, the LSTM systematically shifts probability mass away from predicting NND and toward predicting ED (positive changes in the ED column and negative changes in the NND column). In other words, the LSTM architecture partially recovers information that distinguishes ED from NND, but at the cost of slightly more confusion between PD and ED and a modest reduction in NND accuracy. This trade-off suggests that even with a different representation of the trees, the ED scenario remains only weakly recoverable from PD and NND.

The bottom row of Figure 2 examines how model complexity affects scenario-level recoverability when using the GNN classifier. For trees generated under the medium-complexity (four-parameter) `eve` model (bottom-left panel), PD recall is high (about 79%), NND recall is moderate (about 54%), and ED recall remains low (about 21%). When complexity is increased to the full six-parameter model (bottom-middle panel), the change panel (bottom right) reveals that, across all true scenarios, the high-complexity setting induces a shift toward predicting NND.

Taken together, these results indicate that scenario information in `eve` trees is unevenly distributed and, unsurprisingly, degrades as the number of free parameters increases. At the scenario level, increased model flexibility in `eve` potentially leads to practical non-recoverability between evolutionary relatedness mechanisms.

#### 3.1.2. Accuracy metrics

The sliced accuracy curves in Figure 3 and Figure 4 show that, once we condition on scenario and complexity, differences between the GNN and LSTM architectures are modest compared with the effects of the underlying diversification parameters. Across most panels the two architectures track each other closely, and all three accuracy measures (F1, precision, recall) exhibit qualitatively similar trends. The main determinants of classification performance are the strength and direction of the diversification effects and, to a lesser extent, tree size and model complexity.

**Figure 3:**
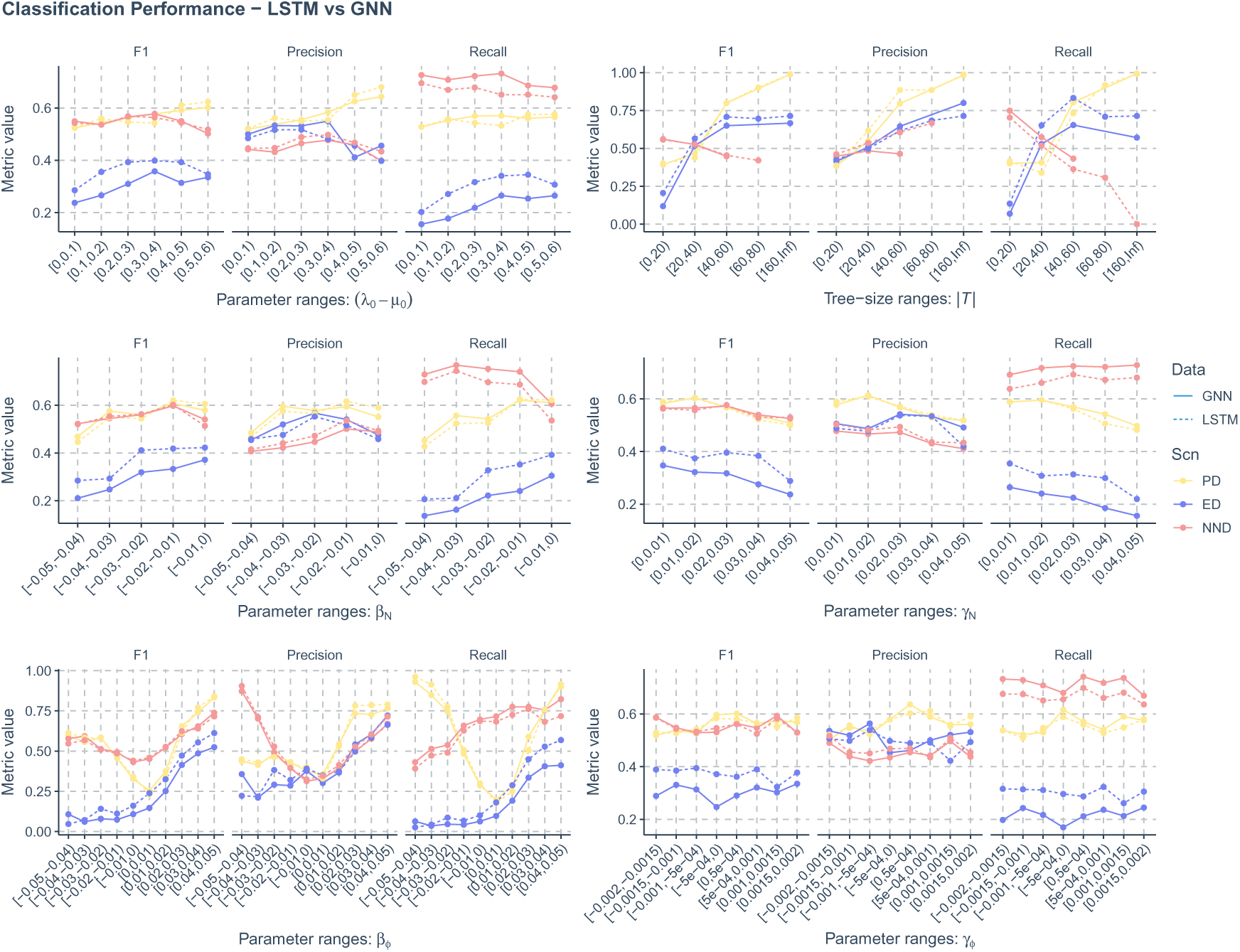
Comparison of trends of neural network classification accuracies (changing along true value slices of the parameters) between GNN (graph neural networks) and LSTM (long short-term memory recurrent networks) performances, indicated by solid and dashed lines, and between three evolutionary scenarios (true scenario under which the trees were generated). Light yellow lines represent performances of trees generated under the phylogenetic diversity (PD) scenario, dark blue lines stand for performances of trees under the evolutionary distinctiveness (ED) scenario, and red lines stand for performances of trees under the nearest neighbor distance (NND) scenario. X-axis: true value slices of the parameters. Y-axis: classification performance metrics, as shown by the column facet strips. See Appendix H for detailed explanation of the three classification performance metrics.

**Figure 4:**
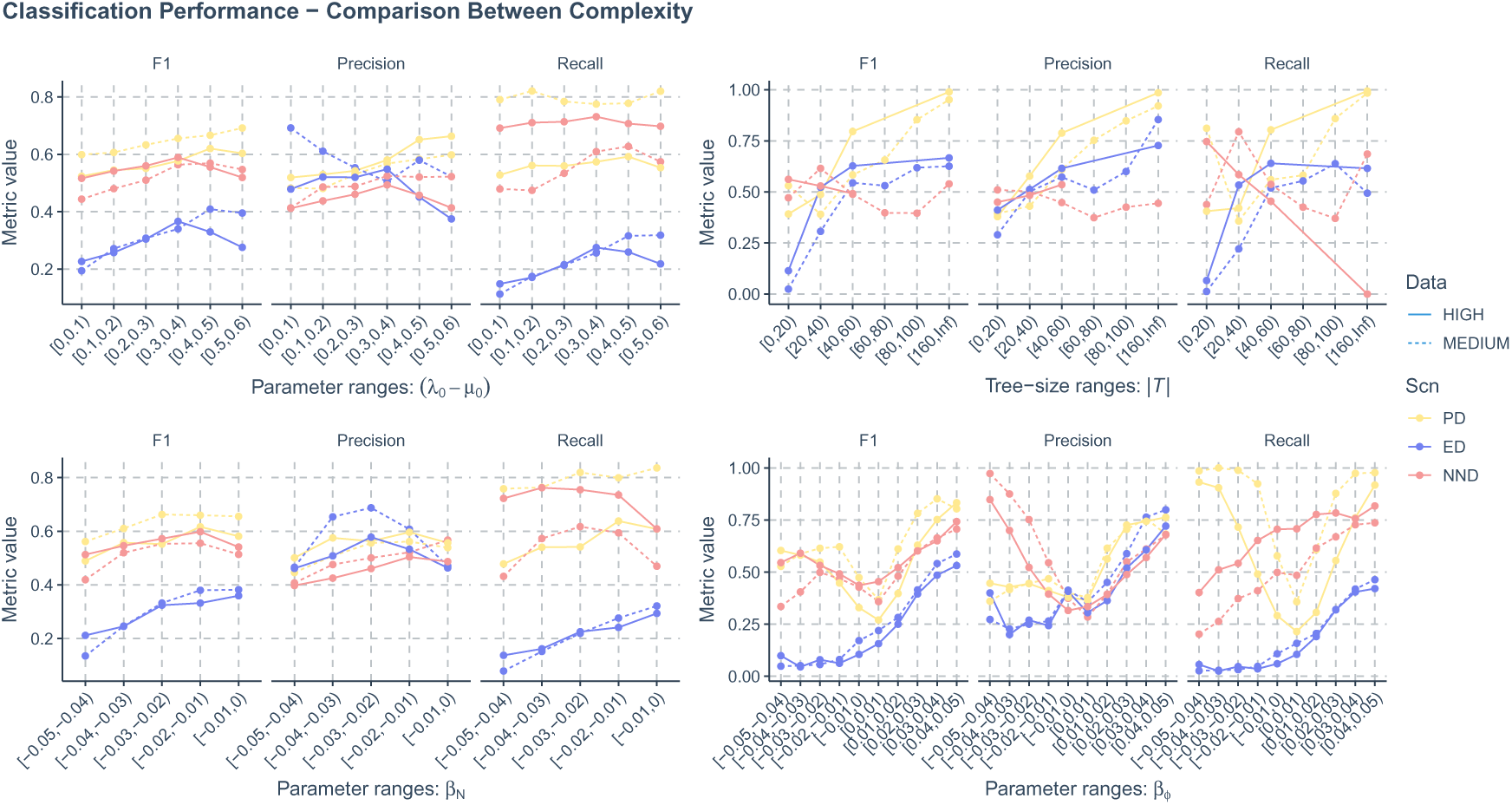
Comparison of trends of neural network classification accuracies (changing along true value slices of the parameters) between medium and high complexity levels (which indicate the number of parameters used to generate the trees) as shown by solid and dashed lines. The low complexity scenario was not considered because parameters other than speciation and extinction rates are not in effect in that scenario. The comparisons were also made between three evolutionary scenarios (true scenario under which the trees were generated). Light yellow lines stand for performances of trees generated under the phylogenetic diversity (PD) scenario, dark blue lines stand for performances of trees under the evolutionary distinctiveness (ED) scenario, and red lines stand for performances of trees under the nearest neighbor distance (NND) scenario. X-axis: true value slices of the parameters. Y-axis: classification performance metrics, as shown by the column facet strips. See Appendix H for detailed explanation of the three classification performance metrics.

Effect sizes, as proxied by the true parameter values or composite quantities, play a central role. An increase in the true net diversification rate (*λ*_0_ − *µ*_0_) is generally associated with higher F1 scores, for PD trees (Figure 3, top row, left most panel). However, this trend flattens or even reverses for ED and NND trees once *λ*_0_ − *µ*_0_ exceeds roughly 0.4. Larger absolute richness effects on speciation and extinction (|*β_N_* | and |*γ_N_* |) consistently depress F1 scores and recall across all scenarios, indicating that strong global diversity dependence tends to obscure differences between PD, ED and NND (Figure 3, middle row). By contrast, increasing the absolute magnitude of the evolutionary relatedness effect on speciation (|*β_Φ_*|) improves classification performance for PD and NND trees, with the highest scores observed for large positive *β_Φ_*. For ED trees, performance improves as *β_Φ_* moves from negative to positive values. No robust, monotonic effect on any of the three metrics is observed for the extinction-relatedness parameter *γ_Φ_*; its influence on scenario recoverability appears weak compared to the speciation components (*β_N_*and *β_Φ_*).

Tree size exerts an additional, mostly positive influence on performance. For PD and ED trees, F1, precision and recall generally increase with the number of nodes, reflecting the fact that larger trees contain more branching events and thus more informative. The pattern for NND trees is less straightforward: accuracy curves are flatter and in some slices even decline in the largest size bins. This is likely due to a combination of smaller typical sizes for NND trees and reduced sample sizes in the upper quantiles (NND trees are generally much smaller in size), so the apparent downturn should be interpreted cautiously. Overall, the consistent improvement for PD and ED with increasing size supports the view that small phylogenies fall into a largely non-recoverable part of parameter space.

In order to assess the impact of model complexity, we only compare medium-and high-complexity versions of the `eve` model because parameters other than speciation and extinction rates (*λ*_0_ and *µ*_0_) are set to zero (and thus not in effect) under the low complexity setting. Similarly, we do not study species richness and evolutionary relatedness effects on extinction rate (*γ_N_* and *γ_Φ_*) because these parameters are only in effect under the high complexity setting. We find that diversification model complexity has scenario-specific effects (Figure 4). For PD trees, moving from the four-parameter to the six-parameter setting generally leads to marginally higher precision but substantially lower recall, resulting in a net reduction in F1 (but these patterns are not observed in the tree size effect panel). Thus, under the high-complexity model the classifier is potentially more conservative: PD predictions are more often correct when they are made, but many true PD trees are reclassified as ED or NND. For ED and NND trees, there are no clear general trends.

#### 3.1.3. Parameter space associated with correct vs. incorrect classifications

To assess which parts of parameter space are associated with successful scenario classification, we compared the generating parameters of correctly and incorrectly classified trees (Figure 5). For each combination of true scenario (PD, ED, NND) and classifier architecture (GNN vs. LSTM), ridge–line densities show how the marginal distributions of *λ*_0_, *µ*_0_, *β_N_*, *β_Φ_*, *γ_N_*and *γ_Φ_* differ between correct and incorrect predictions. We observe that correctly classified trees tend to occupy more extreme regions of parameter space for *β_Φ_* and less negative (smaller absolute value) regions of *β_N_*, although NND trees are an exception. This observation further verifies the confounding effect of *β_N_*and the important role of the absolute effect sizes of *β_Φ_*.

**Figure 5:**
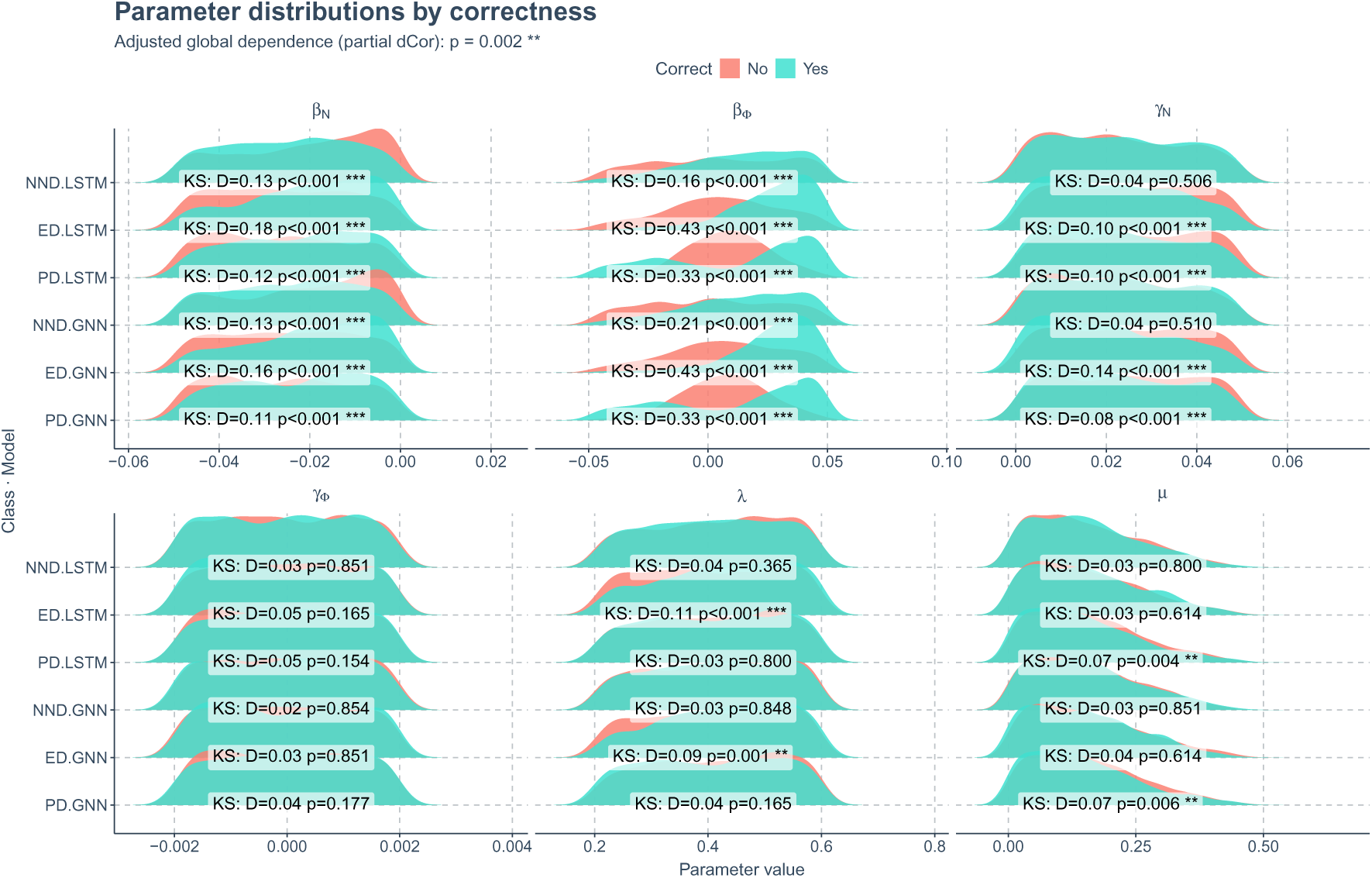
Parameter distributions for correct vs. incorrect classifications. Each panel shows ridge–line kernel density estimates of a generating parameter (*λ*_0_*, µ*_0_*, β_N_, β_Φ_, γ_N_, γ_Φ_*; columns) for a given combination of true scenario (rows within each panel: PD, ED, NND) and classifier architecture (GNN vs. LSTM). Within each ridge–line, turquoise densities correspond to trees that were correctly classified and salmon densities to misclassified trees. On top of the each ridge–lines, we report the two-sample Kolmogorov–Smirnov statistic and Benjamini–Hochberg adjusted *p*-value comparing the parameter distributions between correct and incorrect trees for that scenario-architecture-parameter combination. The adjusted global dependence between the eight-dimensional parameter vector (*λ*_0_*, µ*_0_*, β_N_, β_Φ_, γ_N_, γ_Φ_,* |𝒯|*, λ*_0_ − *µ*_0_) and the correctness indicator after conditioning on scenario and architecture is significant (*p* = 0.002). Together, these summaries highlight which parameters and scenarios exhibit systematic differences between correctly and incorrectly classified trees, and where classification errors appear compatible with a lack of parameter signal.

The one-dimensional Kolmogorov–Smirnov tests reveal that the strongest and most systematic separations between correct and incorrect classifications occur for the richness and relatedness effects on speciation, *β_N_* and *β_Φ_*. For all scenario-architecture combinations, the KS statistics for these parameters are moderate to large and the Benjamini–Hochberg adjusted *p*-values are mostly below 0.01, indicating that the distributions of *β_N_* and *β_Φ_* differ markedly between correctly and incorrectly classified trees. For the other parameters, the patterns show only weak or sporadic differences. For our classifiers, the speciation components (*β_N_* and *β_Φ_*) — particularly the interaction between species richness and evolutionary relatedness — carry more discriminative information than the extinction components (*γ_N_* and *γ_Φ_*) or the baseline rates (*λ*_0_ and *µ*_0_).

Partial distance correlation analysis further indicates a significant global dependence between the eight-dimensional parameter vector (*λ*_0_*, µ*_0_*, β_N_, β_Φ_, γ_N_, γ_Φ_,* |𝒯|*, λ*_0_− *µ*_0_) and the correctness indicator even after conditioning on scenario and architecture (Figure 5, subtitle). Together, these findings imply that classification errors are not purely random but are concentrated in regions of parameter space where effect sizes and tree sizes jointly yield weak or ambiguous signals. Strong richness and relatedness effects carve out parameter space in which the PD, ED and NND scenarios become practically non-recoverable.

#### 3.1.4. Calibration of scenario probabilities

Reliability diagrams for the GNN classifier (Figure 6) show that, when predictions are pooled across all trees, the network is systematically overconfident. In the overall panel, the reliability curve lies below the diagonal for most confidence bins. Predictions with nominal confidence between 0.5 and 0.8 attain observed accuracies closer to 0.4-0.6, and even the highest-confidence bin (*p̂* ≈ 1) reaches an accuracy of only about 0.8. The corresponding expected calibration error (ECE) is therefore non-negligible.

**Figure 6:**
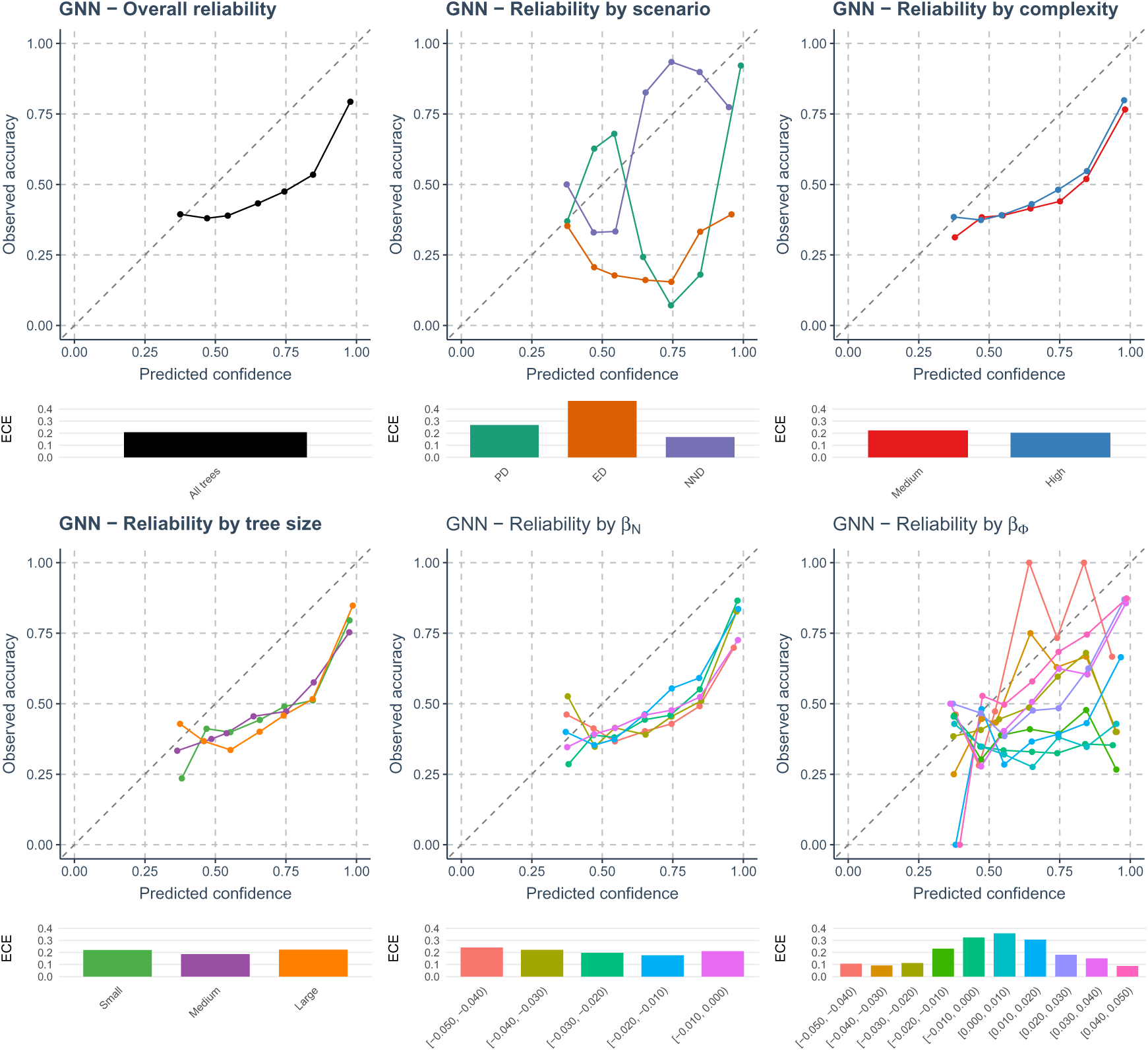
Calibration of the GNN classifier. Each main panel shows a reliability diagram in which the observed accuracy (y-axis) is plotted against the predicted confidence (x-axis; maximum class probability) in ten confidence bins; the dashed diagonal indicates perfect calibration. Top row: overall calibration across all test trees (left); calibration stratified by true diversification scenario (PD, ED, NND; middle); and calibration for GNNs trained on medium- and high-complexity `eve` models (right). Bottom row: calibration stratified by tree size (small, medium, large; left), by ranges of the richness effect on speciation *β_N_* (middle), and by ranges of the relatedness effect on speciation *β_Φ_* (right). Below each reliability panel, a horizontal bar plot reports the expected calibration error (ECE) for each curve, using matching colors; smaller ECE values indicate better calibration.

Stratifying by true diversification scenario (Figure 6, first row, middle panel) reveals that ED is strongly overconfident across all confidence bins, with observed accuracies that are far below the reported confidences and an ECE that is much larger than for PD or NND. This aligns with the confusion matrices (Figure 2), where ED is the hardest scenario to identify. The patterns for PD and NND are asymmetric and non-monotonic.

Calibration differences between medium- and high-complexity `eve` models are relatively minor (see Figure 6, top row, third column). The two complexity-specific curves almost coincide and are both below the diagonal. The increase of free parameters potentially only changes which class is predicted rather than how well probabilities match empirical frequencies. Calibration on all size groups show overconfidence. ECE exhibit no clear trend or difference for different complexities and tree sizes.

Calibration stratified by parameter space indicates that miscalibration depends more strongly on the relatedness effect *β_Φ_* than on the richness effect *β_N_*. Across *β_N_* ranges the reliability curves are fairly similar and moderately overconfident, with modest increase in ECE as |*β_N_* | increases. The *β_Φ_* panel shows larger between-bin variation. ECE generally decrease as |*β_Φ_*| decreases. These results suggest that parameter settings in which *β_Φ_* induces weak or ambiguous changes in tree shape are precisely where both classification performance and probability calibration deteriorate, reflecting a lack of practical recoverability for the `eve` scenarios in those regions of parameter space. This is also true when increasingly negative *β_N_* strengthens richness-driven confounding effect by shrinking the differences in tree summary statistics among PD, ED, and NND, thereby making the scenarios less distinguishable and degrading classification performance, as observed in Qin et al. (2025b).

### 3.2. Regression

Prediction accuracy for the net diversification rate (*λ*_0_ − *µ*_0_) improved with in-creasing true net diversification rate (see top facet rows of all panels in Figure 7), only at the low complexity level. Similar improvement was not observed at medium and high complexity levels, not only for the net diversification rate, but for all the parameters examined. See also the figures in Appendix B.

**Figure 7:**
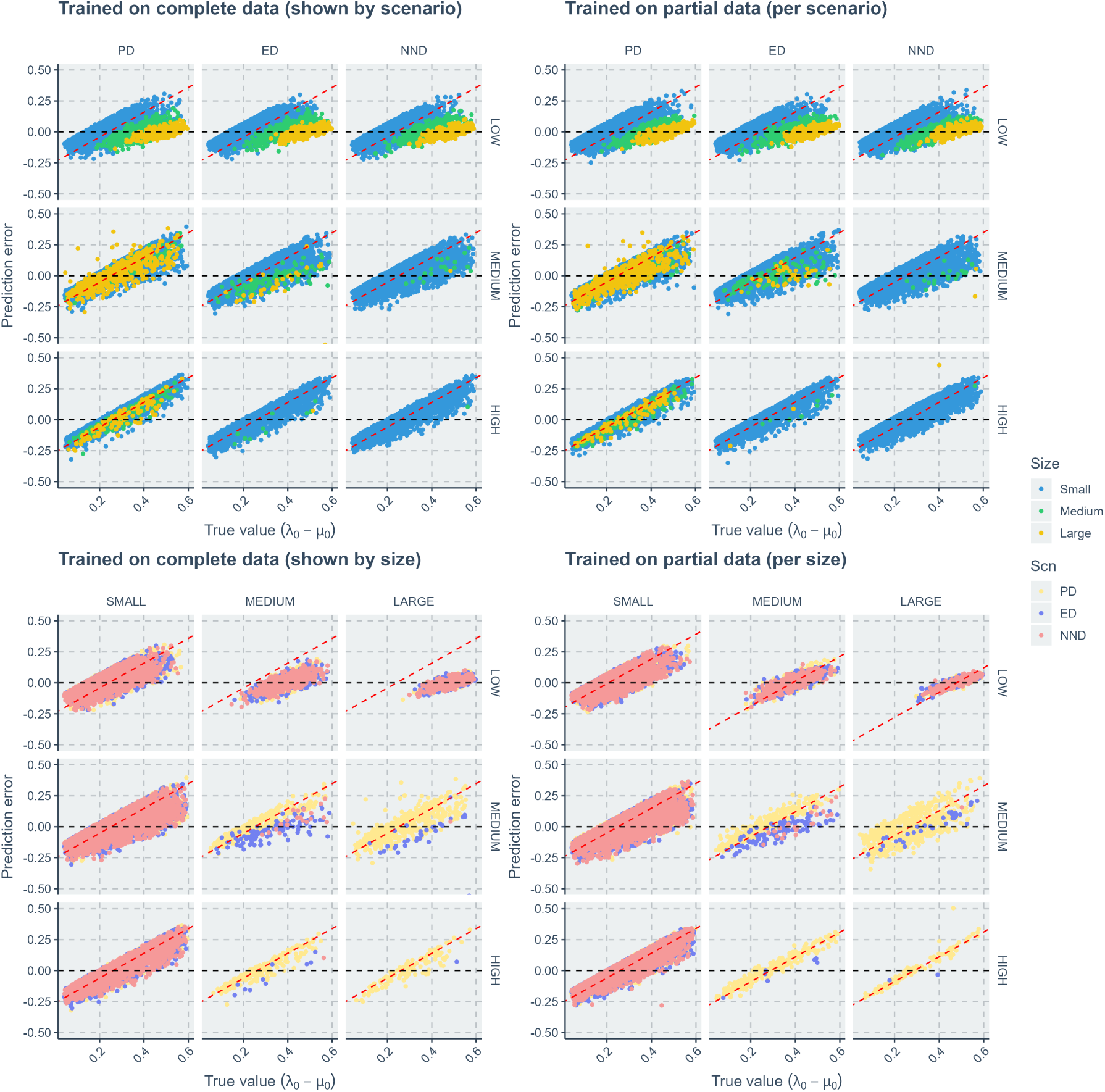
Comparison of neural network regression errors along true values of the net diversification rate (*λ*_0_ − *µ*_0_). The left two panels show the results when the neural networks were trained on complete datasets. The right two panels show the results when the neural networks were trained on partial datasets (training datasets were sliced either by evolutionary scenarios or by tree size groups while in visualization we slice by both). Within each panel, each facet column indicates results of either an evolutionary scenario or a tree size group. Each facet row indicates results of a diversification model complexity. When columns represent scenario groups, blue points correspond to large trees, green points to medium trees, and yellow points to large trees. When columns represent tree size groups, light yellow points correspond to performances of trees generated under the phylogenetic diversity scenario, dark blue points to performances of trees under the evolutionary distinctiveness scenario, and red points to performances of trees under the nearest neighbor distance scenario. X-axis: true value of the net diversification rate (*λ*_0_ − *µ*_0_). Y-axis: prediction error (absolute difference between true value and predicted value).

Training neural network regressors on partial datasets—either sliced by scenario or by tree size—did not yield obvious performance gains compared to training on complete datasets (see the performance differences between the left two panels and the right two panels of Figure 7. See also Figure 8). Further quantitative analyses even showed that training the regressors on partial datasets impaired the neural networks’ generalization ability, with predictions aligning closer to the midpoint of the generative parameter space of the training datasets (see Figure 8 and compare between metrics in the same color group). When trained on complete datasets, the neural networks generally predict better on larger trees for *λ*_0_ − *µ*_0_, *β_N_*, *γ_N_* and *γ_Φ_*. Increasing model complexity degrades performance in estimating the net diversification rate. In more complex scenarios, the expected correlation between larger tree sizes and higher accuracy diminishes, with predictions converging toward the sample mean of the parameters (points close to the red dashed line in Figure 7 and Figure 8; results for other parameters are in Appendix B and Appendix D).

**Figure 8:**
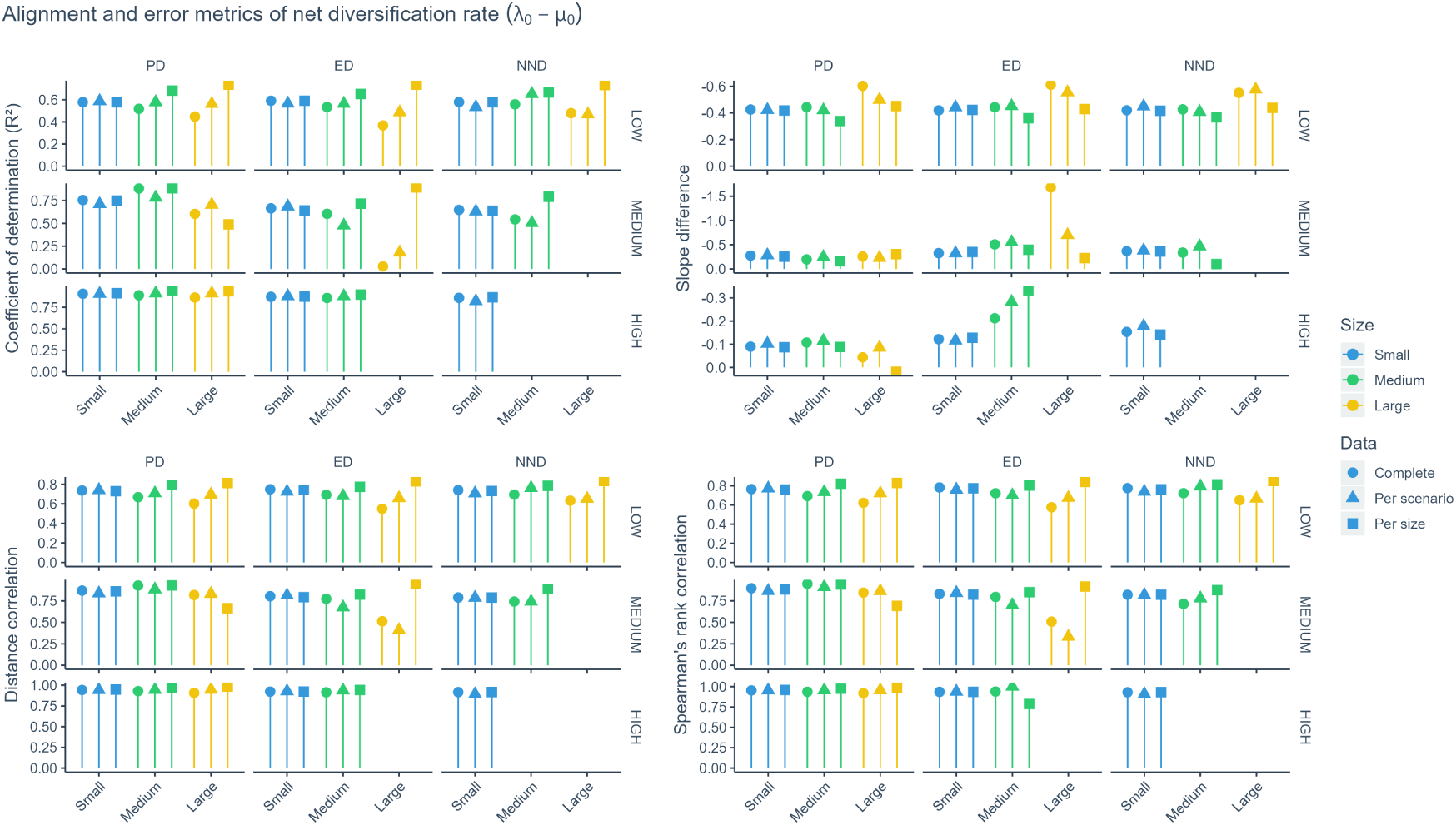
Comparison of the degree to which neural network regression estimates align with the conditional mean. There are four metrics, each describing a facet of the alignment, see Appendix I for details and interpretations. Within each panel, each facet column indicates results of a evolutionary scenario (PD for phylogenetic diversity, ED for evolutionary distinctiveness, NND for nearest neighbor distance); each facet row indicates results of a diversification model differing in complexity (as indicated by the number of parameters used to generate the trees). Yellow bars stand for metrics of small trees, green bars stand for medium trees and blue bars stand for large trees. At the top of the bars, shapes represent how the datasets were sliced to train the neural networks. Circles indicate complete datasets, triangles indicate slices by evolutionary scenario and squares indicate slices by tree size group. X-axis: size group of the trees. Y-axis: value of alignment metrics. As model complexity increases, the simulations rarely yield medium or large trees, so those size groups may be missing in the corresponding panels.

In the low-complexity scenario — a simple birth-death process — net diversification rate estimates (*λ*_0_ − *µ*_0_) appear more accurate when the actual tree sizes were near the mean of the generated sample distribution rather than the theoretical expectation conditioned on the true net diversification rate and crown age (see the left panel of Figure 9 as well as those in Figure F.26, Figure F.27, Figure F.28 and Figure F.29 in Appendix F). In medium- and high-complexity scenarios, although theoretical expectations are unavailable, we observed that net diversification rate estimates, as well as those for other parameters (*β_N_*, *β_Φ_*, *γ_N_*, and *γ_Φ_*), were likewise more accurate when tree sizes approximated the sample’s arithmetic mean. Moreover, tree-size distributions shifted systematically with model complexity: higher-complexity simulations generally produced smaller trees, with the reduction in tree size becoming increasingly pronounced from PD to ED to NND.

**Figure 9:**
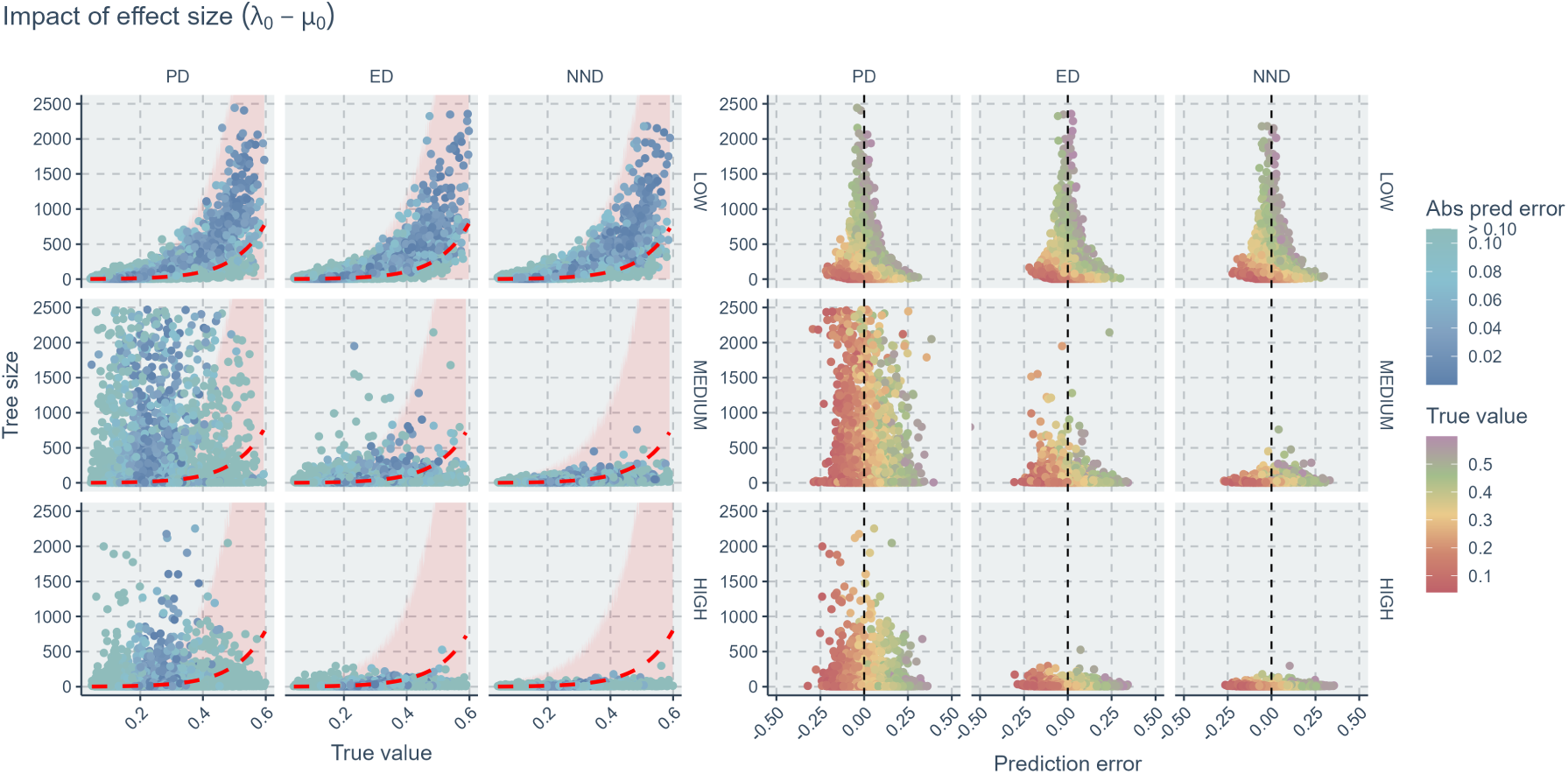
Relationships of tree size to true parameter values and prediction errors of the net diversification rate (*λ*_0_ − *µ*_0_). Within each panel, each column indicates results of an evolutionary scenario (PD for phylogenetic diversity, ED for evolutionary distinctiveness, NND for nearest neighbor distance); each row indicates results of a diversification model complexity (which indicates the number of parameters used to generate the trees). In the left panels, colors indicate the values of absolute prediction errors; as the blue points become darker, the errors become smaller thus the predictions more accurate. The red dashed lines represent expected tree size with respect to true net diversification rate and a fixed crown age 10. The pink ribbons represent possible variations of tree sizes due to stochasticity. In the right panels, the colors indicate the values of true net diversification rate. Points falling close to the vertical black dashed lines are accurate predictions (zero error). X-axis (left panel): true value of the net diversification rate (*λ*_0_ − *µ*_0_). X-axis (right panel): prediction error (absolute difference between true value and predicted value). Y-axis: tree size.

Additionally, larger tree sizes corresponded to more accurate estimates of both the net diversification rate (*λ*_0_ − *µ*_0_, see Figure 7 and Figure 9) and the effect of species richness on extinction (*γ_N_*, see Figure B.14 in Appendix B and Figure F.28 in Appendix F), whereas this trend was less evident for all the other parameters (*β_N_*, *β_Φ_*, and *γ_Φ_*, see the other figures in Appendix B and Appendix F). When a parameter exhibited a weak correlation with tree size, increasing the tree size did not lead to substantially improved estimation accuracy. This is particularly true for the parameters of higher complexity levels (see the performances between panels within the same rows in Figure 9 and Appendix F). In all cases, extremely small trees (with total number of nodes less than 50) were associated with very poor estimates.

In the regressor misspecification test, all `eve` model scenarios are identical under low complexity setting because there is no parameters other than the baseline rates (*λ*_0_ and *µ*_0_) in effect. Misspecification mainly affects the estimates of all parameters when trees are of medium or large size. The neural networks are always conservative by making predictions that are close to the sample mean of the training data on small trees. Regardless of which evolutionary scenario the neural networks were trained on, they produce much worse estimates when recovering parameters from trees generated under the PD scenario (where phylogenetic diversity controls the processes globally). As the real underlying scenarios of the trees to be tested shifting from global to local evolutionary forces, the estimates by the neural networks trained on all scenarios become more stable. See Figure 10 and Figure G.30, Figure G.31, Figure G.32 and Figure G.33 in Appendix G for details.

**Figure 10:**
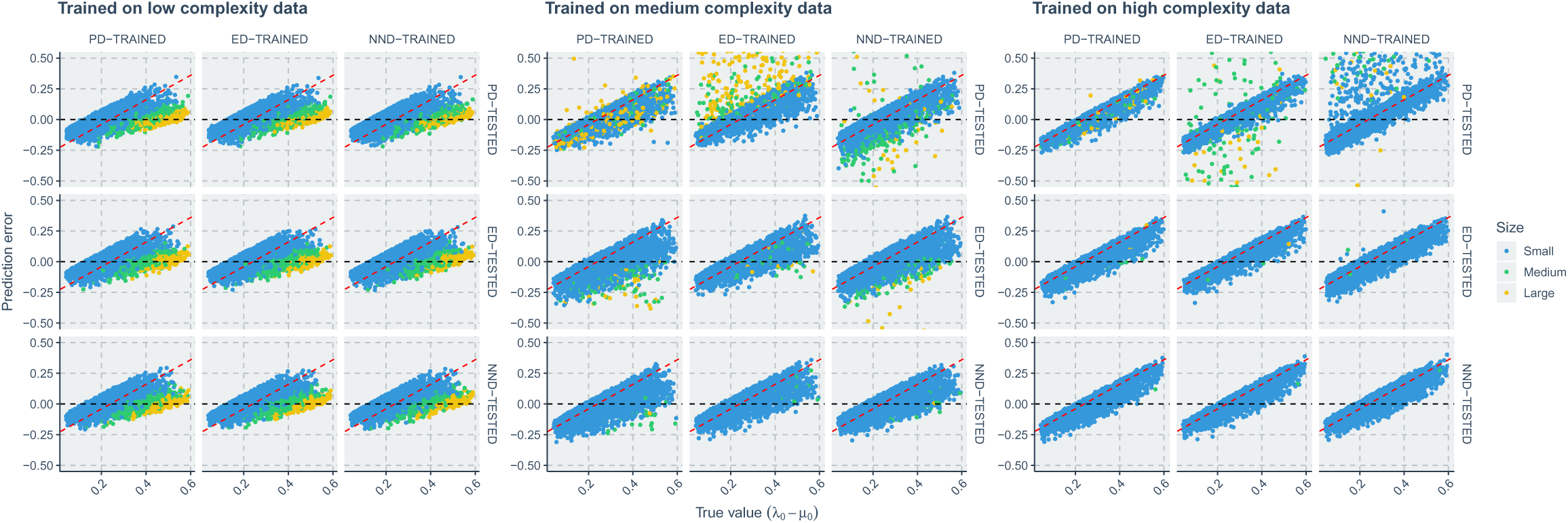
Comparison of neural network regression errors under misspecification. By misspecification, neural networks trained on trees under different evolutionary scenarios are used to estimate parameters on all trees, including those having different (misspecified) evolutionary scenarios. The three panels show the prediction error of three model complexity levels (which indicate the number of parameters used to generate the trees). Within each panel, each column indicates results of using neural networks trained on 3 evolutionary scenarios (see column facet strips, e.g. PD-TRAINED represents neural networks trained on trees generated under the phylogenetic diversity scenario) to estimate parameters on tree dataset generated under a particular scenario (see row facet strips, e.g. PD-TESTED indicate that neural network predictions were made on PD trees). Yellow points stand for results of small trees, green points stand for medium trees and blue points stand for large trees. X-axis: true value of the net diversification rate (*λ*_0_ − *µ*_0_). Y-axis: prediction error (absolute difference between true value and predicted value).

## 4. Discussion

### 4.1. Conservative predictions as indicators of limited information

Our regression experiments show that, for most `eve` parameters and especially in the medium- and high-complexity settings, neural networks often return predictions close to the empirical mean of the training distribution (e.g., Figure 7, Figure 8, Appendix B, Appendix D). This pattern is strongest for the ER parameters (*β_N_*, *β_Φ_*, *γ_N_*, *γ_Φ_*), where slopes of the error-versus-truth relationships and alignment metrics indicate that predictions behave almost like constant conditional means. Under standard loss functions such as squared error or Huber loss, this behavior is expected whenever the inputs carry little information about the target: in low-signal situations, the risk-minimizing solution is to predict the mean of the conditional distribution rather than to chase noisy fluctuations (similar situation was found in Qin et al. (2025a)). In our setting, the conservative predictions therefore point to parts of `eve` parameter space where extant trees potentially do not provide enough information for the networks to recover individual parameter values. Practically, this means that point estimates in those parts of parameter space should be interpreted as “typical values compatible with the data” rather than as precise predictions of the true parameters. Inference workflows that rely on such predictors should therefore report broad uncertainty, treat estimates as lower-dimensional summaries (e.g. of net diversification) or explicitly acknowledge that the available data support many alternative diversification histories, consistent with analytical results on non-identifiability of birth-death models from extant trees as presented by (Louca and Pennell, 2020; Pannetier et al., 2021).

### 4.2. Tree size, effect sizes and limits of parameter recovery

Across both classification and regression analyses, tree size and the strength of diversification effects emerge as the main determinants of how much information the trees potentially contain about the underlying process. For classification, F1 scores, precision and recall generally increase with tree size for PD and ED trees (Figure 3 and Figure 4). Very small trees are generally associated with very low scenario accuracy regardless of parameter values, which aligns with classical findings that tests of phylogenetic signal and model-based comparative methods have little power on small trees (Blomberg et al., 2003; Townsend et al., 2012), analogous to non-identifiability caused by small sample size in time-series hidden Markov models Cole (2019). The pattern for NND trees is less clear because these trees are typically small and occupy a narrow range of sizes, so even the largest NND trees in our simulations contain fewer branching events than PD and ED trees of comparable parameter values. For regression, larger trees help most clearly for estimates of net diversification rate and *γ_N_*, with errors decreasing with size when model complexity is low and, to a lesser extent, at medium complexity (Figure 7, Figure 9, Figure B.14). However, once the full six-parameter `eve` model is used, size-related improvements become modest or minimal and predictions for several parameters collapse back towards the empirical mean. Importantly, the trees that yield the most accurate estimates are not those with extreme sizes but those whose sizes are close to the typical sizes produced by the simulation models (Figure 9 and Figure F.29), suggesting that the networks effectively learn a mapping from size and a few coarse topological summaries to parameter averages, with limited ability to exploit more subtle shape information. After all, note that under our low-complexity setting (effectively a constant-rate birth-death process), the unlabeled tree topology is known to be independent of the speciation and extinction rates and therefore carries no information about the generative parameter (Steel and McKenzie, 2001; Gernhard, 2008). It is not surprising that our GNN-based approaches bring no substantial gain (see also (Qin et al., 2025a)). Together, these results indicate that working with larger clades of organisms helps only up to a point: larger trees improve recovery of a small number of parameters that tightly control overall diversification pace, but they do little for parameters that mainly modulate subtle ER patterns once diversity dependence and stochasticity are included.

Effect sizes play an equally important role. Across scenarios and architectures, strong species richness dependence (|*β_N_* | and |*γ_N_* |) consistently reduces classification accuracy and degrades regression performance. When richness effects are large, tree size reacts sensitively (we would expect smaller trees), and the resulting variation in size can overwhelm the weaker signatures of ER effects, making it harder for the networks to distinguish PD, ED and NND or to disentangle the contributions of *β_Φ_* and *γ_Φ_*. In contrast, strong relatedness effects on speciation (|*β_Φ_*|) improve scenario separability: PD and NND trees with large positive *β_Φ_* are classified more accurately, and correct trees in these scenarios preferentially occupy extreme *β_Φ_* values while avoiding strongly negative *β_N_* (Figure 5). For ED trees, shifting *β_Φ_* from negative to positive values gradually improves performance, which aligns with a shift from antagonistic to reinforcing interactions between richness and relatedness on speciation (see (Qin et al., 2025b) for how ED trees are generated). The combination of parameter-sliced accuracy curves, ridge-line distributions and statistical tests therefore provides an empirical map of the parts of parameter space where `eve` parameters and scenarios potentially leave strong enough imprints on tree shape to be recoverable, and the parts where their effects are washed out by stochasticity and richness dependence.

### 4.3. Scenario overlap and redundancy in complex eve models

Our scenario-level analyses show that, even when neural networks perform better than chance, PD, ED and NND are far from being cleanly separated in tree space. Confusion matrices reveal that NND trees are generally easiest to classify, while ED trees are consistently the most difficult (Figure 2). Many ED trees are mislabeled as NND, and PD trees are more often confused with NND than vice versa, indicating that the local ER mechanisms of ED and the neighborhood-based mechanism of NND frequently generates tree shapes that are compatible with the PD scenario. Increasing `eve` complexity exacerbates this problem: when the six-parameter model is used, misclassifications shift towards NND for all true scenarios, and scenario-specific recalls decline even though the networks are trained and tested on data generated exactly from the fitted model. The ridge-line summaries potentially also supports the idea, as correctly and incorrectly classified trees often arise from overlapping ranges of net diversification and ER effect sizes, with clear separations only in parameter-size combinations where *β_Φ_* is strong and tree size is moderate to large (see Figure 5). It can also be seen that regression and misspecification experiments tell a similar story. Even when the networks are trained on all scenarios simultaneously, estimates for ER parameters remain conservative and close to empirical means for most of the parameter space (Figure 8, Figure 10, Appendix G). We propose to visualize scenario redundancy as overlapping “clouds” of trees in representation space: the networks can detect broad differences between scenarios in parts of parameter space where ER effects are strong and diversity dependence is moderate, but they cannot reliably assign trees to a unique scenario in the more typical ranges of parameters explored by our simulations.

We further illustrate how overlapping clusters of trees in a hypothetical representation space can obscure scenario-specific signals and drive conservative, mean-regressing predictions by our classifiers and regressors as we observed, in Figure 11. This schematic is generated from toy simulations.

**Figure 11:**
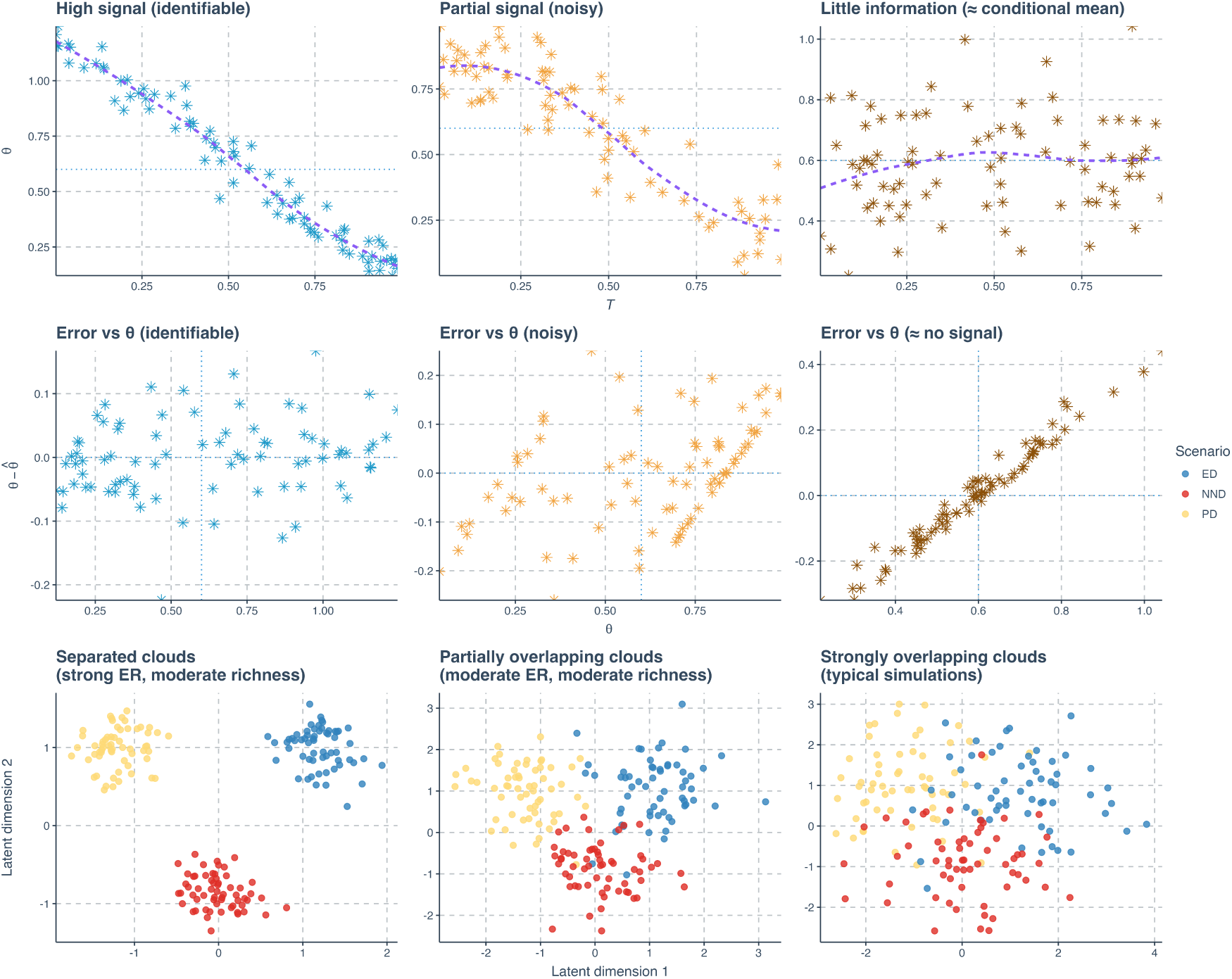
Illustration of how increasing model complexity (flexibility) reduces the information that tree summaries carry about diversification parameters and scenarios. **Top row:** for three hypothetical settings, points show simulated pairs of a tree-derived summary 𝒯 and a parameter value *θ*, the violet dashed curve is a LOESS (locally estimated scatterplot smoothing) estimate of the conditional mean 𝔼[*θ* | 𝒯], and the blue horizontal line marks the unconditional mean 𝔼[*θ*]. From left to right, the relationship between 𝒯 and *θ* weakens: a clear trend (high signal) is followed by a noisy trend (partial signal) and finally by an almost flat curve concentrated around the mean (little information, with many distinct parameter values producing similar summaries). **Middle row:** corresponding errors in the hypothetical estimation of parameters, where each point shows *θ* – 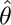 against the true *θ*; as signal decreases, errors become more structured and eventually align with a straight line, indicating that predictions collapse toward an almost constant conditional mean. **Bottom row:** schematic two-dimensional “representation spaces” under two hypothetical latent dimensions for three diversification scenarios (PD, ED, NND) under analogous conditions. When ER effects are strong and richness dependence is moderate (left), the three scenarios occupy well-separated clouds; with weaker or more noisy signal (middle), the clouds partially overlap; in parameter ranges similar to our typical simulations (right), the clouds strongly overlap, illustrating that the networks can still detect broad differences among scenarios in some parts of parameter space, but cannot reliably assign individual trees to a unique scenario when their representations become nearly indistinguishable.

### 4.4. Practical lessons and recommendations

Although our goal was not to build production-ready predictors, the results provide several practical lessons for using neural networks with complex diversification models. First, both classification and regression outputs are most trustworthy in specific combinations of parameter values and tree sizes. Scenario labels from our classifiers are most informative for large trees (roughly more than 200 nodes) with strong positive relatedness effects on speciation and moderate richness dependence; in those cases F1 scores approach 0.8-0.9 for some scenarios, confusion matrices are dominated by the diagonal (but not for ED), and parameter distributions of correct and incorrect trees differ. In contrast, for small trees, for weak or strongly negative *β_Φ_*, or when |*β_N_* | and |*γ_N_* | are large, our results indicate that extant trees alone do not contain enough information to support confident scenario assignments. In applied settings with complex diversification models, we therefore recommend treating our classifier outputs (or other potential neural classifiers alike) as qualitative “scenario hints” outside of the high-information parts of parameter space, and avoiding strong biological interpretations of small differences in scenario probabilities when F1 scores are low.

Second, for parameter estimation, our regressors are primarily useful for tracking broad trends in net diversification and a few extinction-related effects, not for precise recovery of the full `eve` parameter vector. Estimates for *λ*_0_ − *µ*_0_ and *γ_N_*improve with tree size and, at low complexity, can capture coarse gradients across the parameter space; however, predictions for *β_N_*, *β_Φ_* and *γ_Φ_* are often shrunk towards empirical means and show limited correlation with the truth. In practice, this means that neural networks trained on simulated `eve` trees could be used only as fast approximators for low-dimensional summaries, such as net diversification rate or composite ER indices, but should not be relied on for full parameter recovery unless extensive validation shows that the empirical trees lie in a high-information part of the simulated space.

Third, our calibration analysis highlights that miscalibration is most severe for the hardest scenario in terms of recoverability (ED) and for parameter ranges where relatedness effects are weak or ambiguous. Across most strata the GNN is overconfident, with predicted probabilities overstating actual accuracies by 10-30 percentage points. For downstream applications that require probabilistic statements, this suggests incorporating explicit calibration steps (e.g. temperature scaling or isotonic regression; (Zadrozny and Elkan, 2002; Niculescu-Mizil and Caruana, 2005; Guo et al., 2017)) trained on held-out simulated data, or moving to approaches that out-put predictive intervals or sets—for example, conformal prediction (Shafer and Vovk, 2008; Lei et al., 2018; Angelopoulos and Bates, 2022) or Bayesian neural architec-tures (Gal and Ghahramani, 2016; Blundell et al., 2015; Lakshminarayanan et al., 2017)—so that lack of information is expressed as wide uncertainty rather than over-confident point predictions. Such approaches cannot create information that is not present in the trees, but they can reduce the risk of over-interpretation.

Finally, our results underscore that attempts to fit highly flexible diversification models to single empirical trees should be paired with careful simulation-based diagnostics. Before estimating parameters or comparing complex scenarios on real data, one can run our workflow (or similar simulation pipelines) under plausible parameter ranges and tree sizes and check whether the empirical tree falls into a region where classification and regression errors are acceptably low. If not, simpler models tailored to specific hypotheses (e.g., omitting certain scenarios, constraining parameters to a lower-dimension, or focusing a subset of parameters) may be more appropriate. Complementary data sources (e.g., fossils, traits, spatial information or replicated trees across clades) are likely necessary to break the many-to-one mappings between diversification histories and extant trees (Louca and Pennell, 2020).

## 5. Acknowledgements

We thank the Center for Information Technology of the University of Groningen for their support and for providing access to the Hábrók high performance computing cluster.

## Appendix A. Loss Terms

In regression tasks, total loss comprises three key components: Huber loss, link prediction loss and entropy of regularization. Huber loss was used for optimizing regression accuracy while the remaining components focused on alleviating a possible issue where GNN can be hard to train, if incorporating the differentiable pooling method (Ying et al., 2018).

The Huber loss (Huber, 1992) for vectors *y* and *ŷ*, each with *n* elements, computed as the average loss across all elements, is given by:

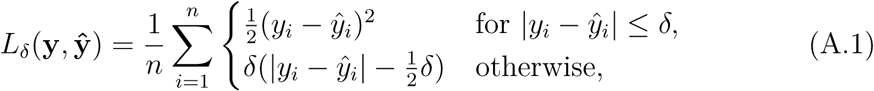

where **y** is the true value vector comprising the ground truth parameters used for simulating a phylogenetic tree, **ŷ** is the predicted value vector comprising the parameter predictions, *y_i_* and *ŷ_i_* are the *i*-th elements of **y** and **ŷ** respectively, *n* is the number of elements in the vectors **y** and **ŷ** and *δ* is the threshold parameter that defines the transition from squared to linear loss (here loss refers to the difference between ground truth and predicted values). In our research, we set *δ* = 0.8 for all the training sessions, making the neural networks more sensitive to smaller errors and more robust to outliers.

The total loss function *L*_1_ in regression is given by

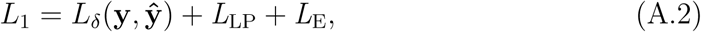

where *L*_LP_ is the link prediction loss and *L*_E_ is the entropy of regularization, see Ying et al. (2018) for their definitions.

In classification tasks, we replaced the Huber loss component with cross-entropy loss for the purpose of multi-class classification. It measures the difference between the true class labels and the predicted probabilities, penalizing confident but incorrect predictions more heavily. For a set of *n* instances, the cross-entropy loss between the true labels **y** and the predicted probabilities **ŷ** is defined as:

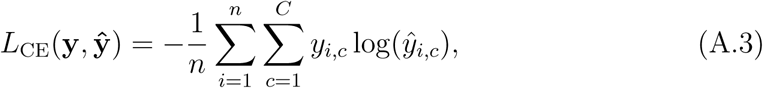

where *n* is the number of instances in the dataset, *C* is the total number of classes, *y_i,c_* is a binary indicator (0 or 1) if class label *c* is the correct classification for instance *i*, *ŷ_i,c_* is the predicted probability that instance *i* belongs to class *c* and log(*ŷ_i,c_*) is the natural logarithm of the predicted probability.

The total loss *L*_2_ in classification is given by

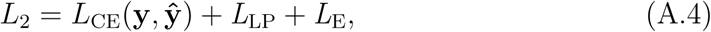

where *L*_LP_ is the link prediction loss and *L*_E_ is the entropy of regularization.

## Appendix B. Complete and Partial Data Training

**Figure B.12:**
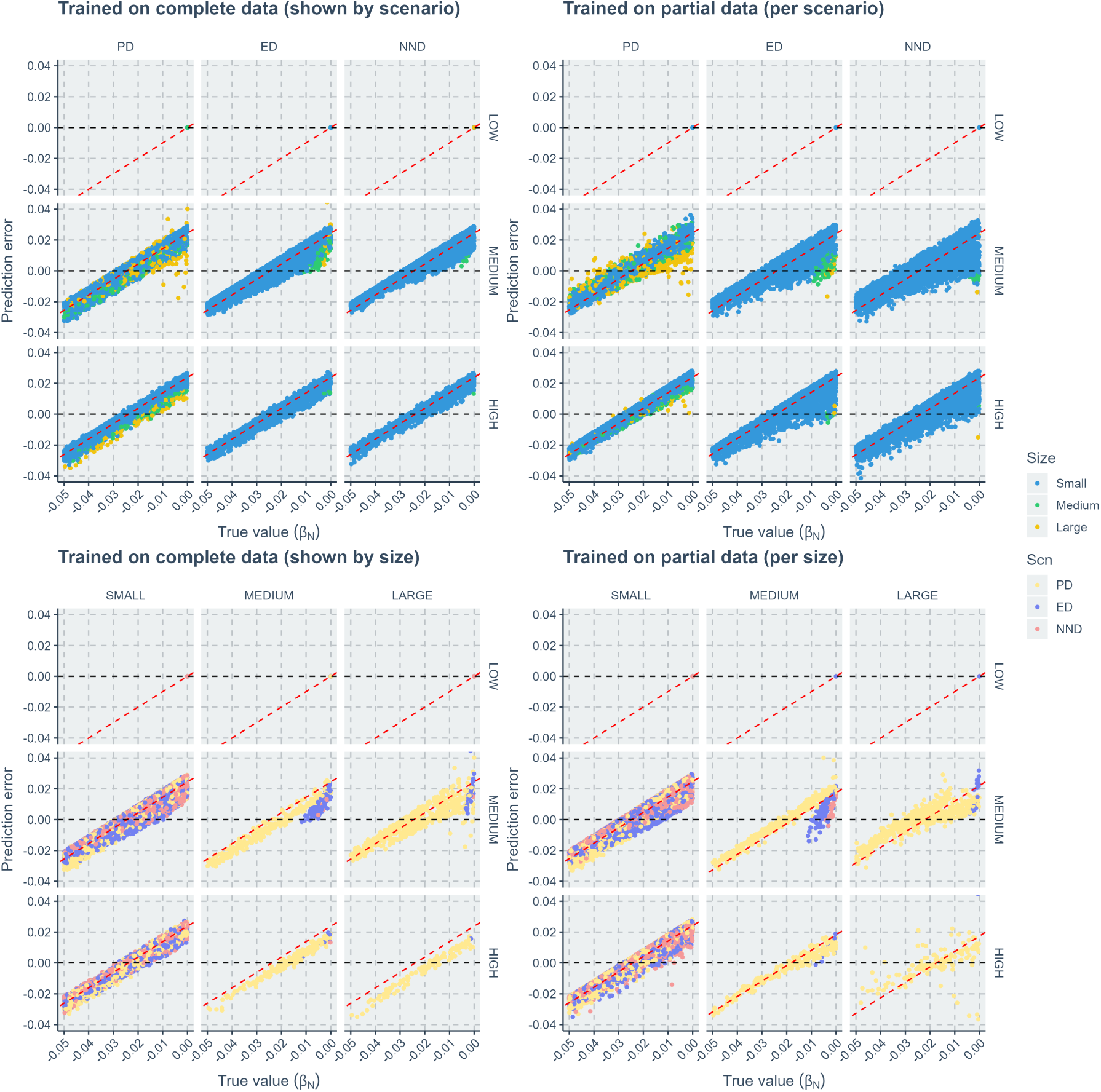
Comparison of neural network regression errors along true values of the species richness effect size on speciation (*β_N_*). The left two panels show the results when the neural networks were trained on complete datasets. The right two panels show the results when the neural networks were trained on partial datasets (training datasets were sliced either by evolutionary scenarios or by tree size groups while in visualization we slice by both). Within each panel, each facet column indicates results of either an evolutionary scenario or a tree size group. Each facet row indicates results of a diversification model complexity. When columns represent scenario groups, blue points correspond to large trees, green points to medium trees, and yellow points to large trees. When columns represent tree size groups, light yellow points correspond to performances of trees generated under the phylogenetic diversity scenario, dark blue points to performances of trees under the evolutionary distinctiveness scenario, and red points to performances of trees under the nearest neighbor distance scenario. X-axis: true value of the species richness effect size on speciation (*β_N_*). Y-axis: prediction error (absolute difference between true value and predicted value).

**Figure B.13:**
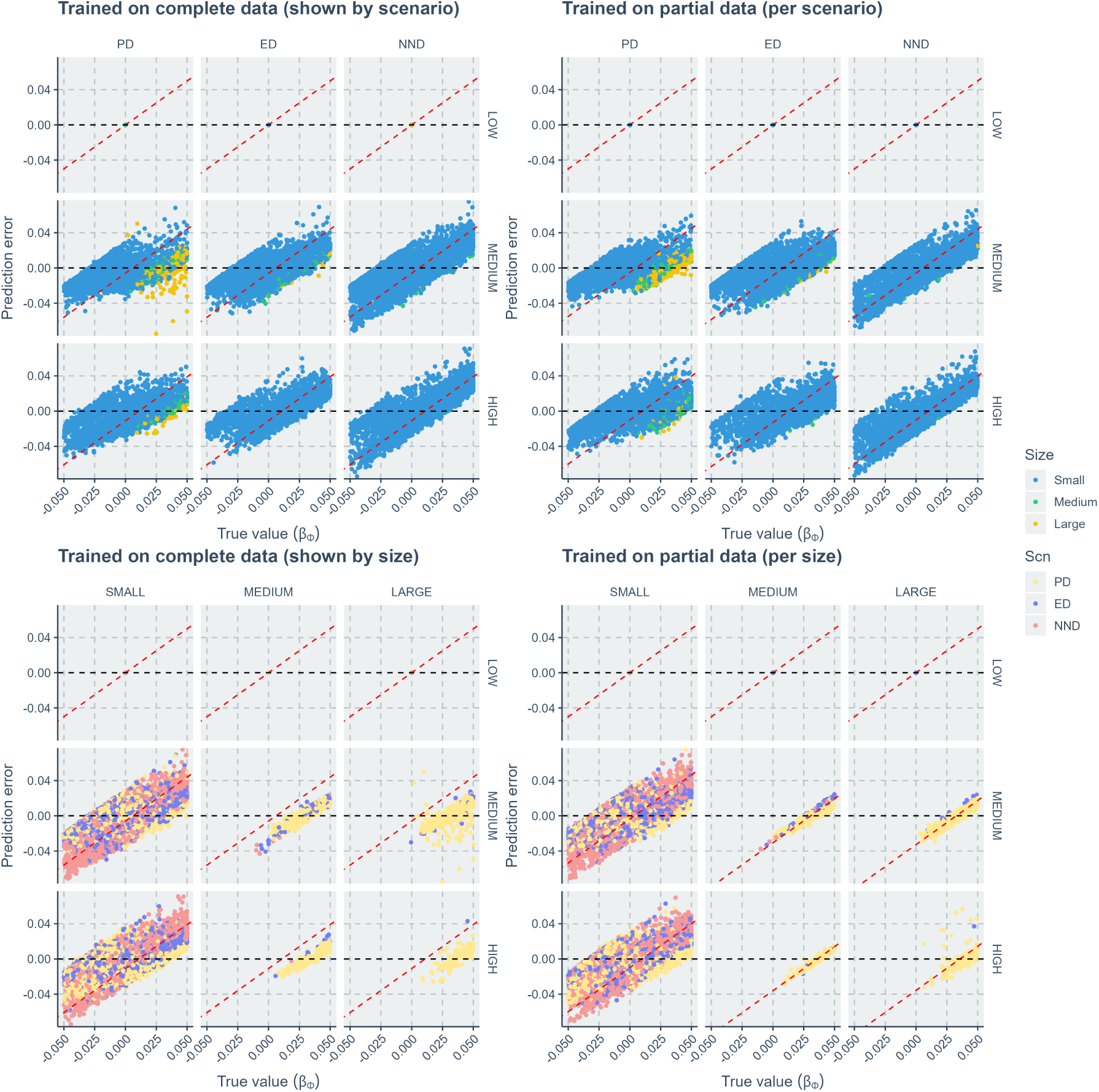
Comparison of neural network regression errors along true values of the evolutionary relatedness effect size on speciation (*β_Φ_*). The left two panels show the results when the neural networks were trained on complete datasets. The right two panels show the results when the neural networks were trained on partial datasets (training datasets were sliced either by evolutionary scenarios or by tree size groups while in visualization we slice by both). Within each panel, each facet column indicates results of either an evolutionary scenario or a tree size group. Each facet row indicates results of a diversification model complexity. When columns represent scenario groups, blue points correspond to large trees, green points to medium trees, and yellow points to large trees. When columns represent tree size groups, light yellow points correspond to performances of trees generated under the phylogenetic diversity scenario, dark blue points to performances of trees under the evolutionary distinctiveness scenario, and red points to performances of trees under the nearest neighbor distance scenario. X-axis: truevalue of the evolutionary relatedness effect size on speciation (*β_Φ_*). Y-axis: prediction error (absolute difference between true value and predicted value).

**Figure B.14:**
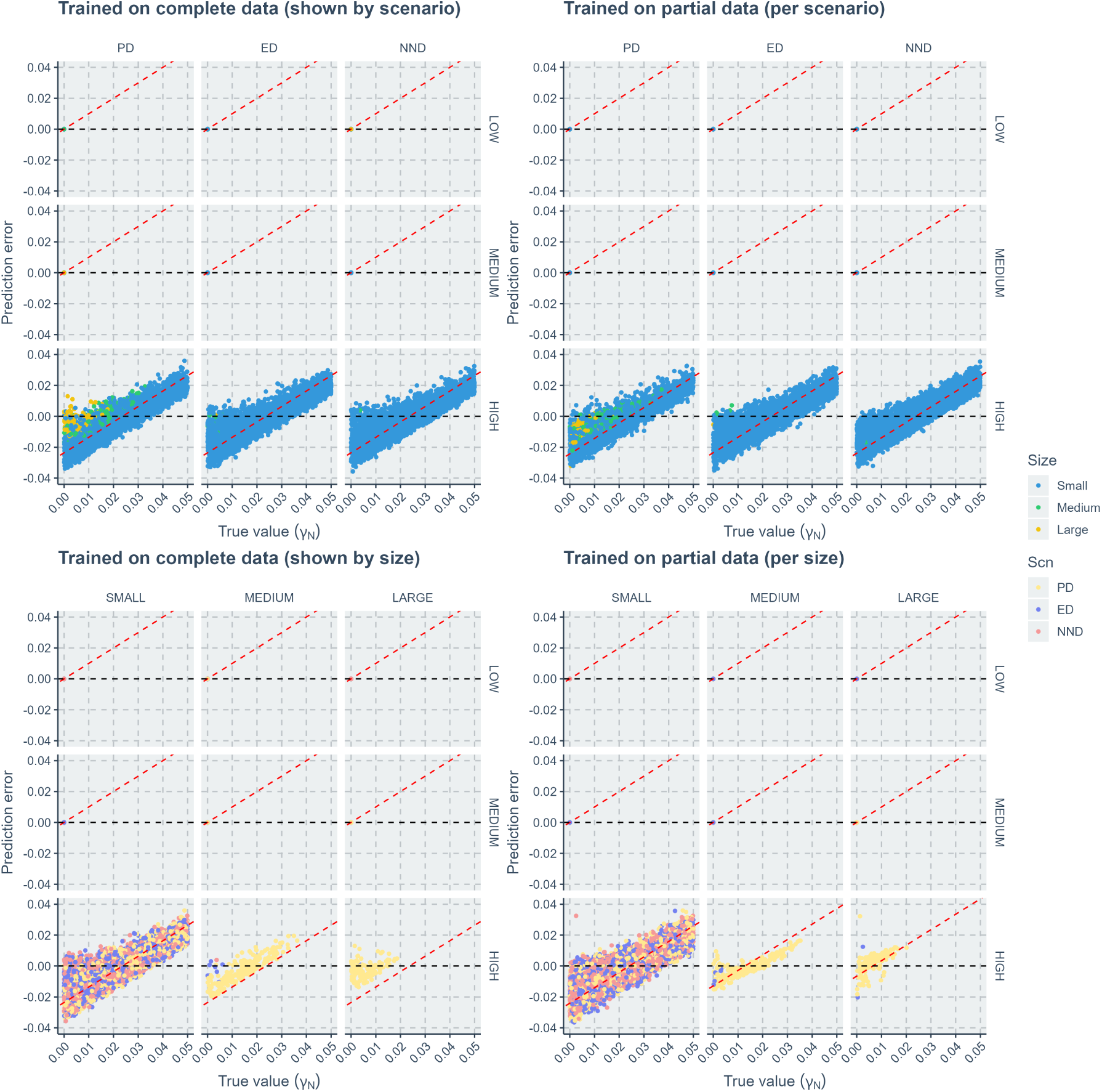
Comparison of neural network regression errors along true values of the species richness effect size on extinction (*γ_N_*). The left two panels show the results when the neural networks were trained on complete datasets. The right two panels show the results when the neural networks were trained on partial datasets (training datasets were sliced either by evolutionary scenarios or by tree size groups while in visualization we slice by both). Within each panel, each facet column indicates results of either an evolutionary scenario or a tree size group. Each facet row indicates results of a diversification model complexity. When columns represent scenario groups, blue points correspond to large trees, green points to medium trees, and yellow points to large trees. When columns represent tree size groups, light yellow points correspond to performances of trees generated under the phylogenetic diversity scenario, dark blue points to performances of trees under the evolutionary distinctiveness scenario, and red points to performances of trees under the nearest neighbor distance scenario. X-axis: true value of the species richness effect size on extinction (*γ_N_*). Y-axis: prediction error (absolute difference between true value and predicted value).

**Figure B.15:**
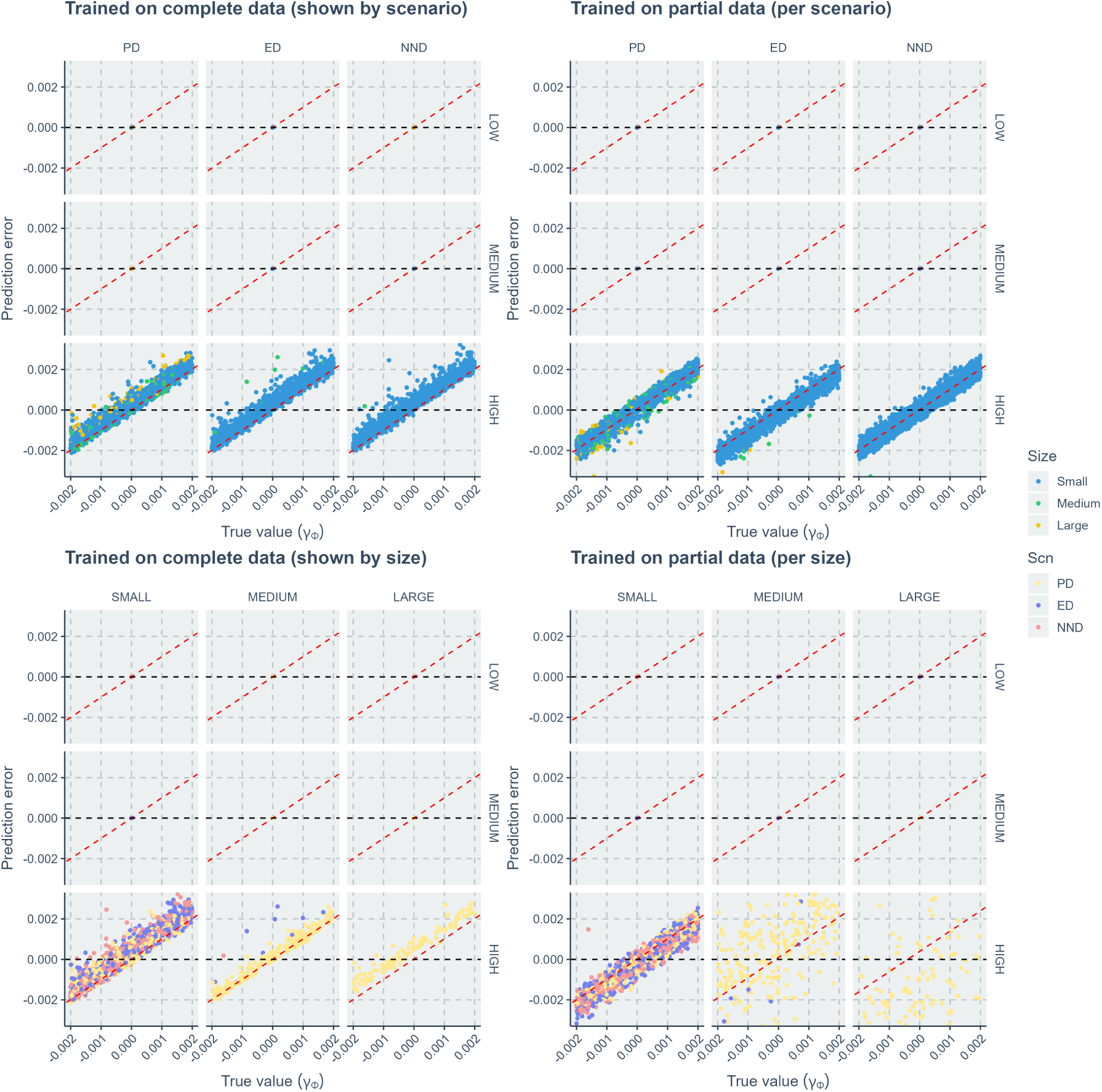
Comparison of neural network regression errors along true values of the evolutionary relatedness effect size on extinction (*γ_Φ_*). The left two panels show the results when the neural networks were trained on complete datasets. The right two panels show the results when the neural networks were trained on partial datasets (training datasets were sliced either by evolutionary scenarios or by tree size groups while in visualization we slice by both). Within each panel, each facet column indicates results of either an evolutionary scenario or a tree size group. Each facet row indicates results of a diversification model complexity. When columns represent scenario groups, blue points correspond to large trees, green points to medium trees, and yellow points to large trees. When columns represent tree size groups, light yellow points correspond to performances of trees generated under the phylogenetic diversity scenario, dark blue points to performances of trees under the evolutionary distinctiveness scenario, and red points to performances of trees under the nearest neighbor distance scenario. X-axis: true value of the evolutionary relatedness effect size on extinction (*γ_Φ_*). Y-axis: prediction error (absolute difference between true value and predicted value).

## Appendix C. Contour plots of the point estimates

**Figure C.16:**
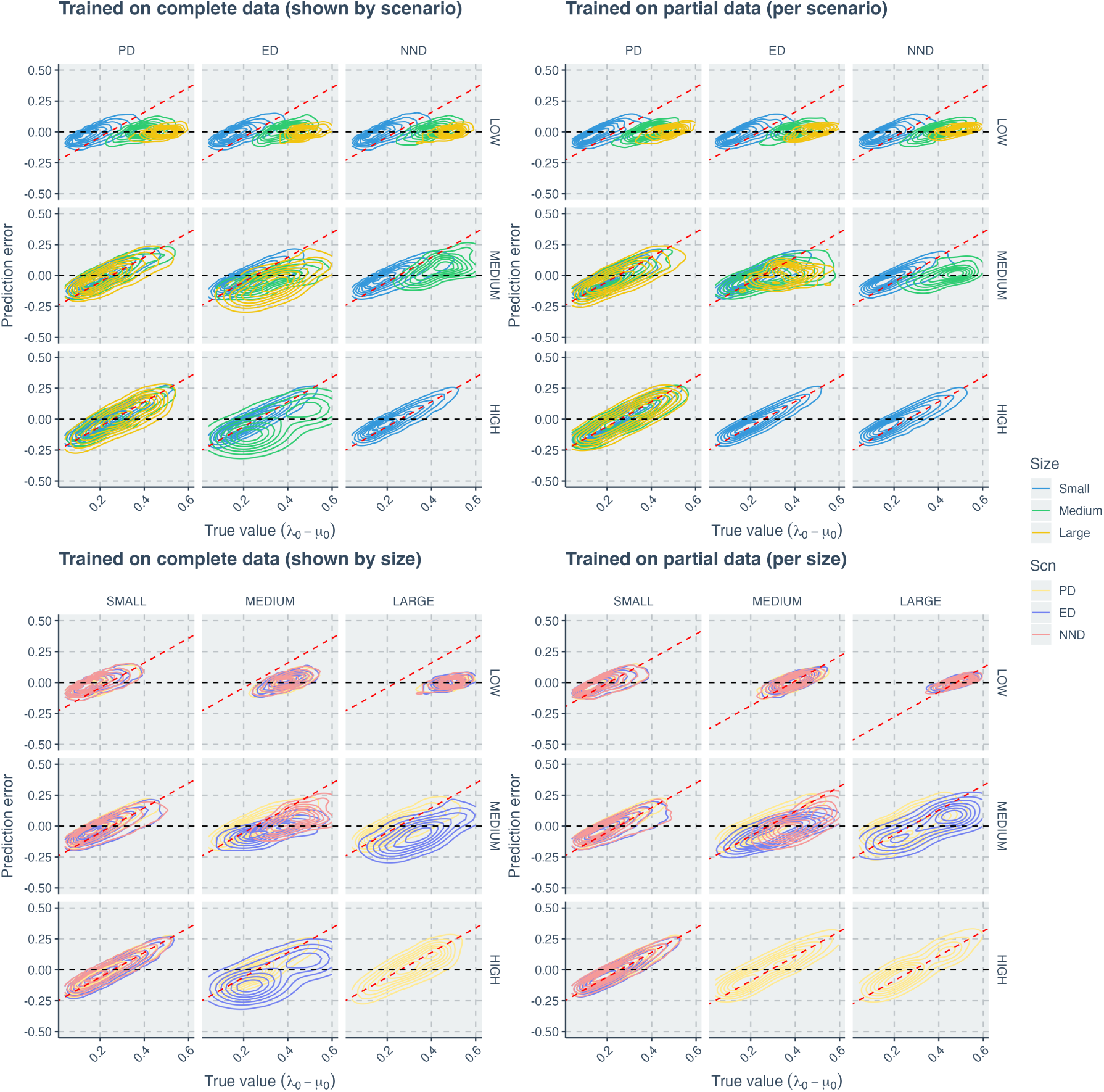
Comparison of neural network regression errors along true values of the net diversification rate (*λ*_0_ − *µ*_0_), using contour plot instead of point cloud. The left two panels show the results when the neural networks were trained on complete datasets. The right two panels show the results when the neural networks were trained on partial datasets (training datasets were sliced either by evolutionary scenarios or by tree size groups while in visualization we slice by both). Within each panel, each facet column indicates results of either an evolutionary scenario or a tree size group. Each facet row indicates results of a diversification model complexity. When columns represent scenario groups, blue contours correspond to large trees, green contours to medium trees, and yellow contours to large trees. When columns represent tree size groups, light yellow contours correspond to performances of trees generated under the phylogenetic diversity scenario, dark blue contours to performances of trees under the evolutionary distinctiveness scenario, and red contours to performances of trees under the nearest neighbor distance scenario. X-axis: true value of the net diversification rate (*λ*_0_ − *µ*_0_). Y-axis: prediction error (absolute difference between true value and predicted value).

**Figure C.17:**
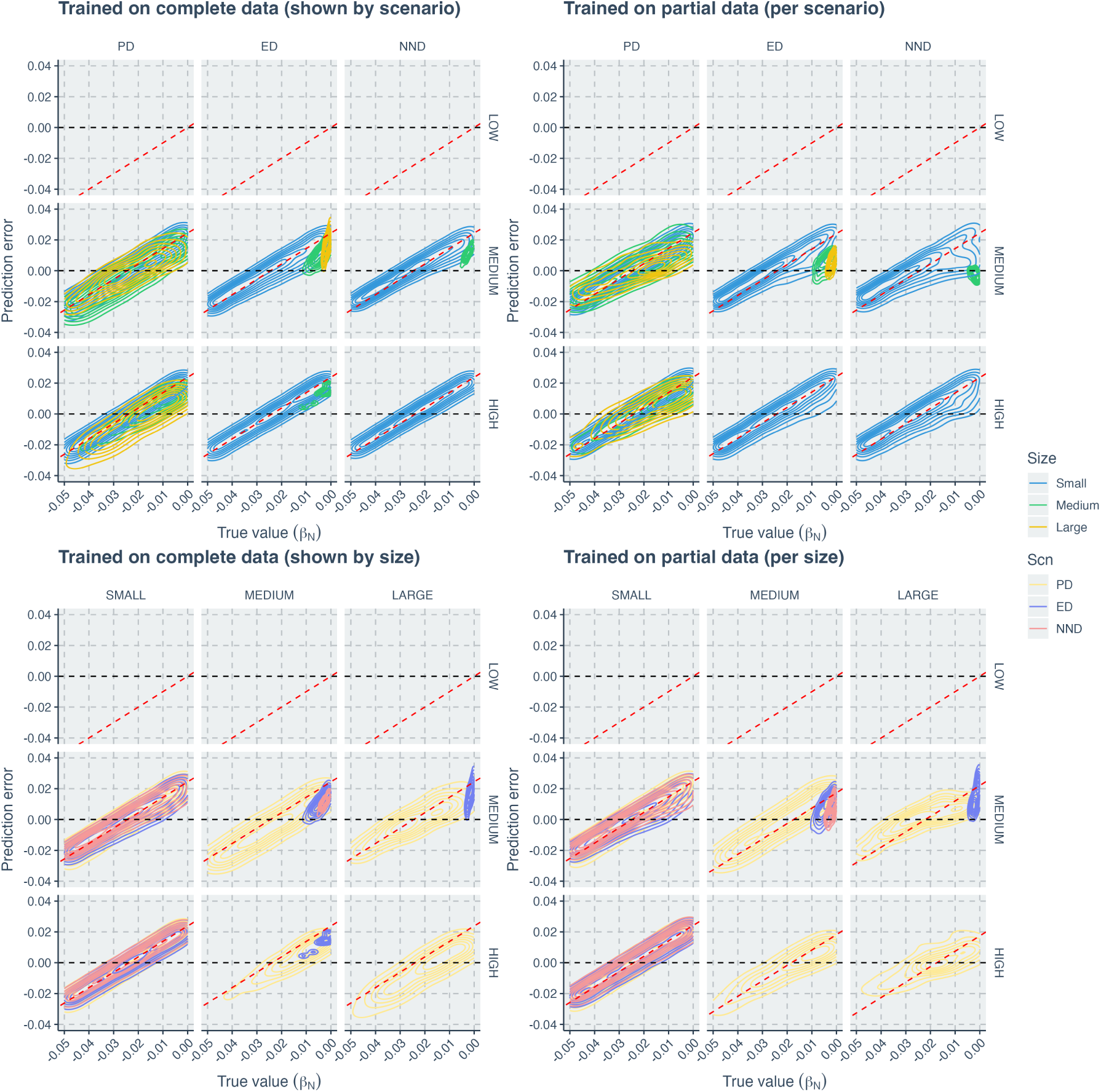
Comparison of neural network regression errors along true values of the species richness effect size on speciation (*β_N_*), using contour plot instead of point cloud. The left two panels show the results when the neural networks were trained on complete datasets. The right two panels show the results when the neural networks were trained on partial datasets (training datasets were sliced either by evolutionary scenarios or by tree size groups while in visualization we slice by both). Within each panel, each facet column indicates results of either an evolutionary scenario or a tree size group. Each facet row indicates results of a diversification model complexity. When columns represent scenario groups, blue contours correspond to large trees, green contours to medium trees, and yellow contours to large trees. When columns represent tree size groups, light yellow contours correspond to performances of trees generated under the phylogenetic diversity scenario, dark blue contours to performances of trees under the evolutionary distinctiveness scenario, and red contours to performances of trees under the nearest neighbor distance scenario. X-axis: true value of the species richness effect size on speciation (*β_N_*). Y-axis: prediction error (absolute difference between true value and predicted value).

**Figure C.18:**
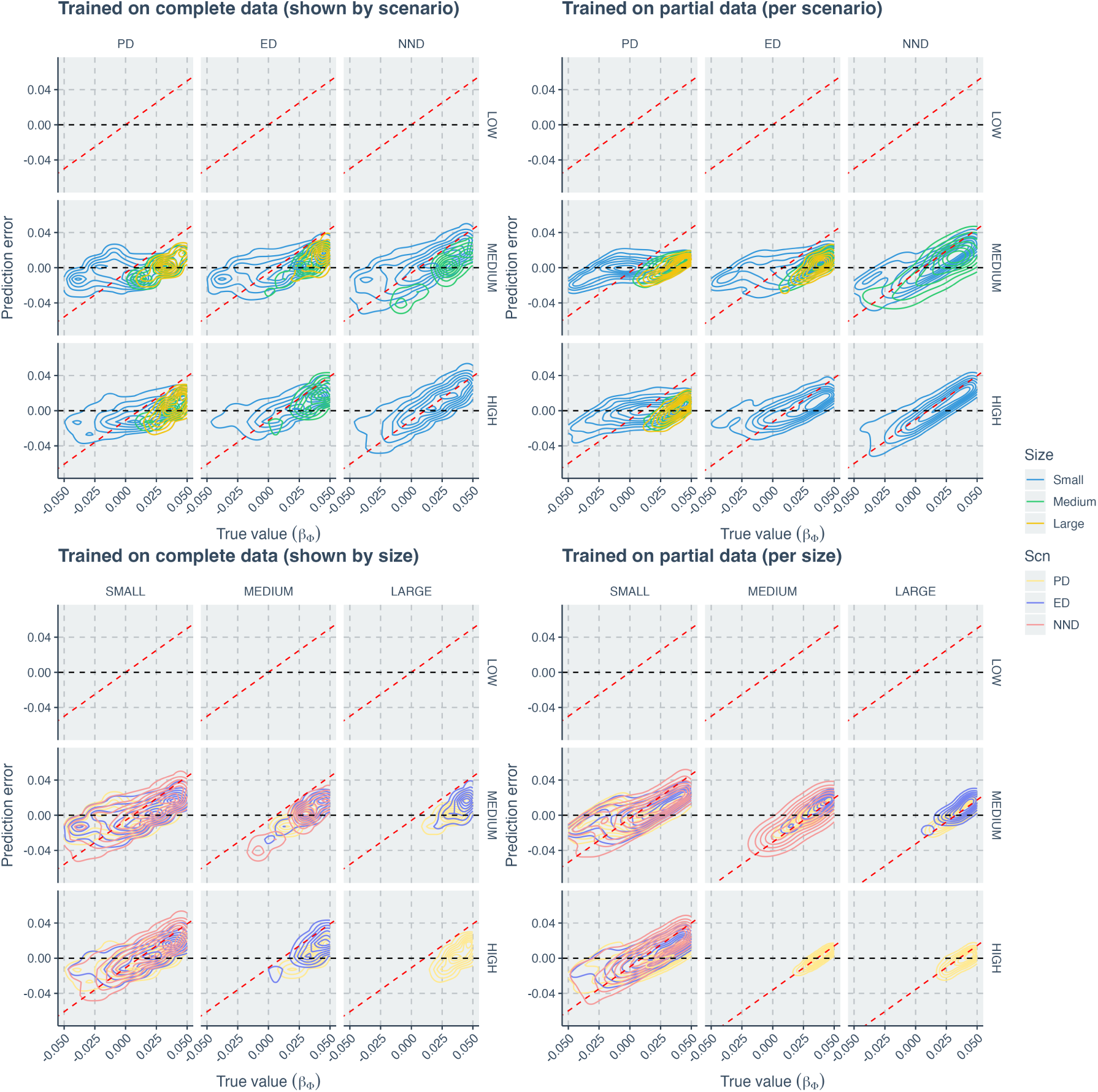
Comparison of neural network regression errors along true values of the evolutionary relatedness effect size on speciation (*β_Φ_*), using contour plot instead of point cloud. The left two panels show the results when the neural networks were trained on complete datasets. The right two panels show the results when the neural networks were trained on partial datasets (training datasets were sliced either by evolutionary scenarios or by tree size groups while in visualization we slice by both). Within each panel, each facet column indicates results of either an evolutionary scenario or a tree size group. Each facet row indicates results of a diversification model complexity. When columns represent scenario groups, blue contours correspond to large trees, green contours to medium trees, and yellow contours to large trees. When columns represent tree size groups, light yellow contours correspond to performances of trees generated under the phylogenetic diversity scenario, dark blue contours to performances of trees under the evolutionary distinctiveness scenario, and red contours to performances of trees under the nearest neighbor distance scenario. X-axis: true value of the evolutionary relatedness effect size on speciation (*β_Φ_*). Y-axis: prediction error (absolute difference between true value and predicted value).

**Figure C.19:**
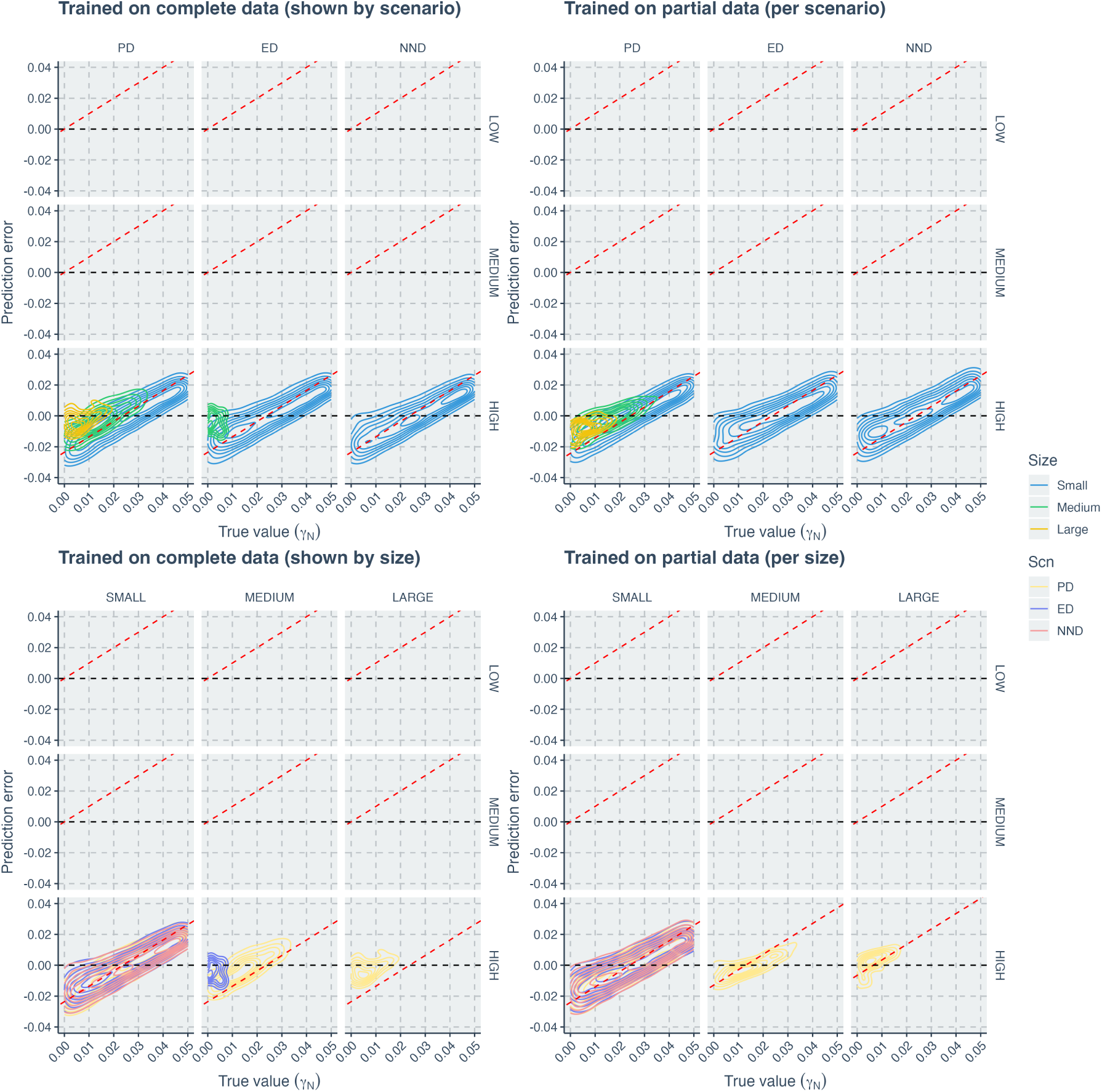
Comparison of neural network regression errors along true values of the species richness effect size on extinction (*γ_N_*), using contour plot instead of point cloud. The left two panels show the results when the neural networks were trained on complete datasets. The right two panels show the results when the neural networks were trained on partial datasets (training datasets were sliced either by evolutionary scenarios or by tree size groups while in visualization we slice by both). Within each panel, each facet column indicates results of either an evolutionary scenario or a tree size group. Each facet row indicates results of a diversification model complexity. When columns represent scenario groups, blue contours correspond to large trees, green contours to medium trees, and yellow contours to large trees. When columns represent tree size groups, light yellow contours correspond to performances of trees generated under the phylogenetic diversity scenario, dark blue contours to performances of trees under the evolutionary distinctiveness scenario, and red contours to performances of trees under the nearest neighbor distance scenario. X-axis: true value of the species richness effect size on extinction (*γ_N_*). Y-axis: prediction error (absolute difference between true value and predicted value).

**Figure C.20:**
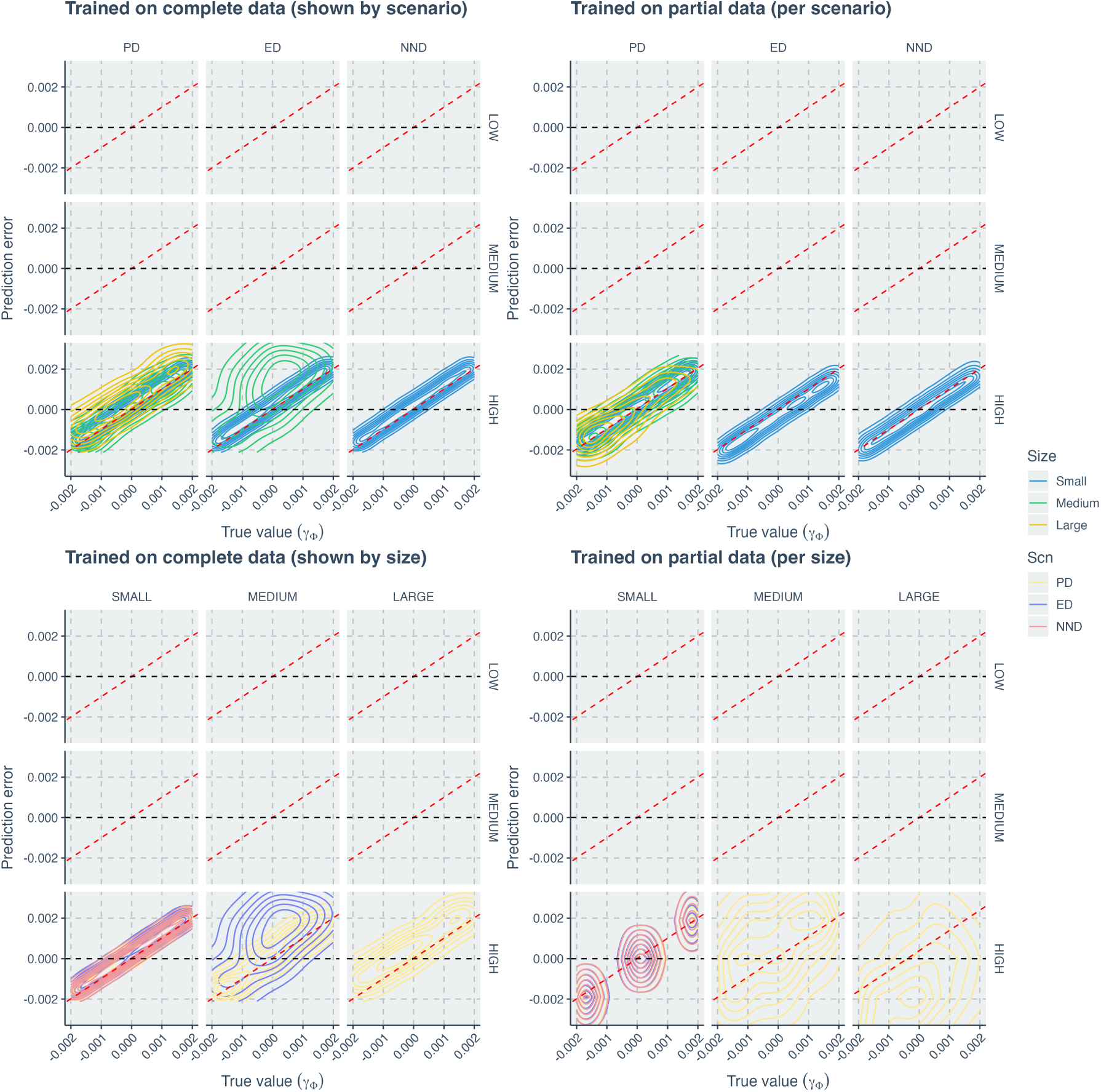
Comparison of neural network regression errors along true values of the evolutionary relatedness effect size on extinction (*γ_Φ_*), using contour plot instead of point cloud. The left two panels show the results when the neural networks were trained on complete datasets. The right two panels show the results when the neural networks were trained on partial datasets (training datasets were sliced either by evolutionary scenarios or by tree size groups while in visualization we slice by both). Within each panel, each facet column indicates results of either an evolutionary scenario or a tree size group. Each facet row indicates results of a diversification model complexity. When columns represent scenario groups, blue contours correspond to large trees, green contours to medium trees, and yellow contours to large trees. When columns represent tree size groups, light yellow contours correspond to performances of trees generated under the phylogenetic diversity scenario, dark blue contours to performances of trees under the evolutionary distinctiveness scenario, and red contours to performances of trees under the nearest neighbor distance scenario. X-axis: true value of the evolutionary relatedness effect size on extinction (*γ_Φ_*). Y-axis: prediction error (absolute difference between true value and predicted value).

## Appendix D. Alignment with Conditional Mean

**Figure D.21:**
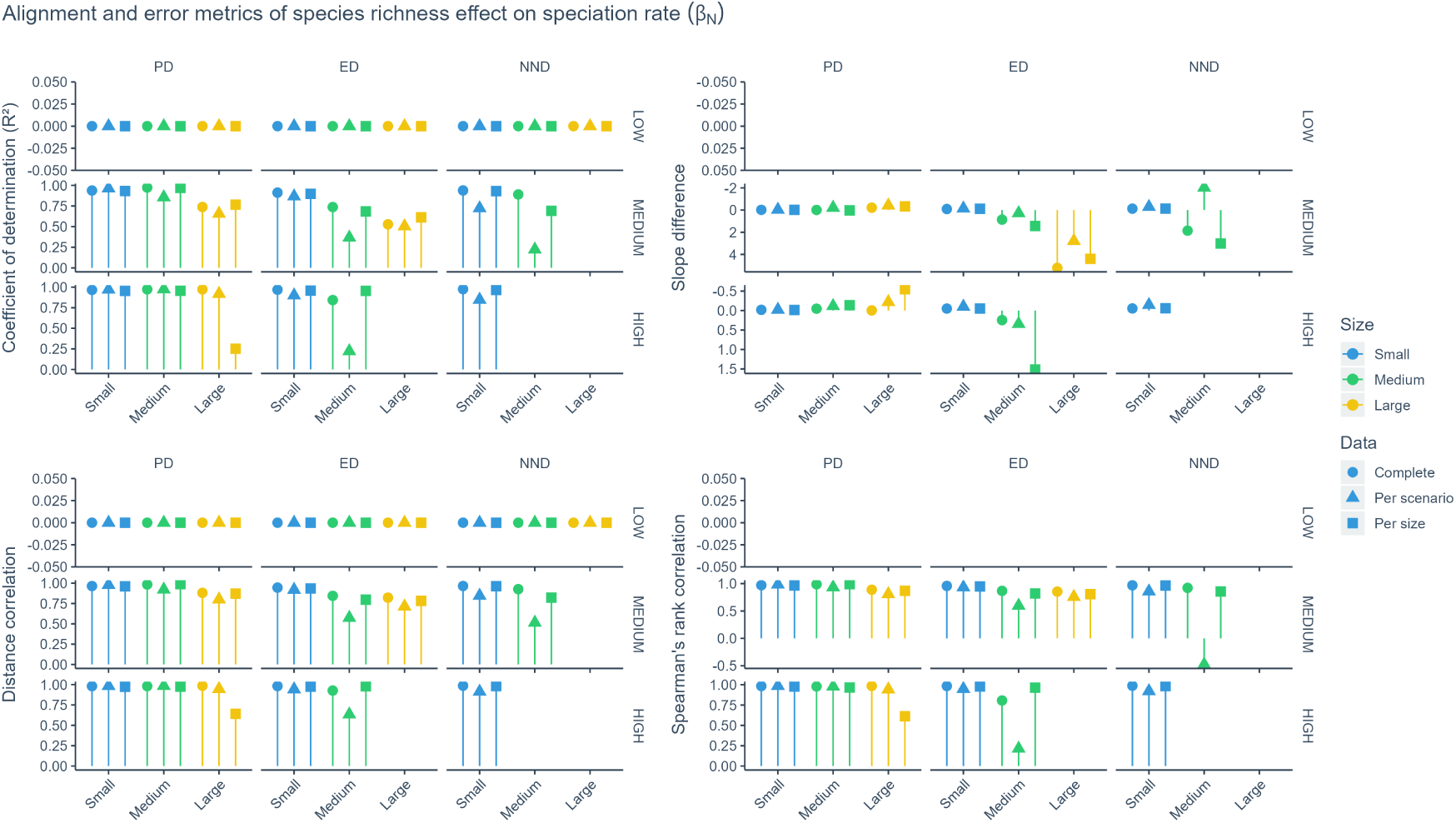
Comparison of the degree to which neural network regression estimates align with the conditional mean. There are four metrics, each describing a facet of the alignment, see Appendix I for details and interpretations. Within each panel, each facet column indicates results of a evolutionary scenario (PD for phylogenetic diversity, ED for evolutionary distinctiveness, NND for nearest neighbor distance); each facet row indicates results of a diversification model differing in complexity (as indicated by the number of parameters used to generate the trees). Yellow bars stand for metrics of small trees, green bars stand for medium trees and blue bars stand for large trees. At the top of the bars, shapes represent how the datasets were sliced to train the neural networks. Circles indicate complete datasets, triangles indicate slices by evolutionary scenario and squares indicate slices by tree size group. X-axis: size group of the trees. Y-axis: value of alignment metrics.

**Figure D.22:**
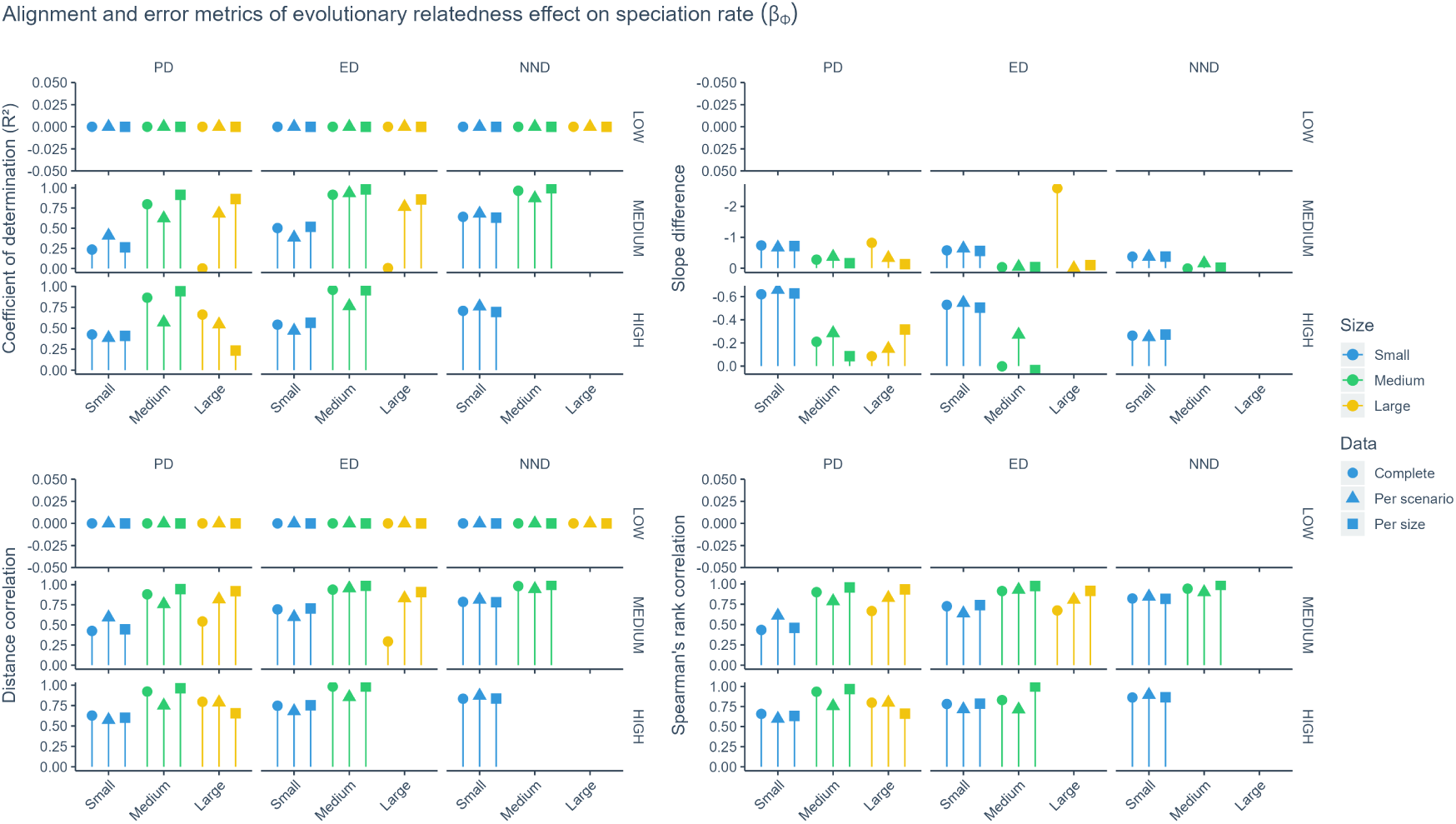
Comparison of the degree to which neural network regression estimates align with the conditional mean. There are four metrics, each describing a facet of the alignment, see Appendix I for details and interpretations. Within each panel, each facet column indicates results of a evolutionary scenario (PD for phylogenetic diversity, ED for evolutionary distinctiveness, NND for nearest neighbor distance); each facet row indicates results of a diversification model differing in complexity (as indicated by the number of parameters used to generate the trees). Yellow bars stand for metrics of small trees, green bars stand for medium trees and blue bars stand for large trees. At the top of the bars, shapes represent how the datasets were sliced to train the neural networks. Circles indicate complete datasets, triangles indicate slices by evolutionary scenario and squares indicate slices by tree size group. X-axis: size group of the trees. Y-axis: value of alignment metrics.

**Figure D.23:**
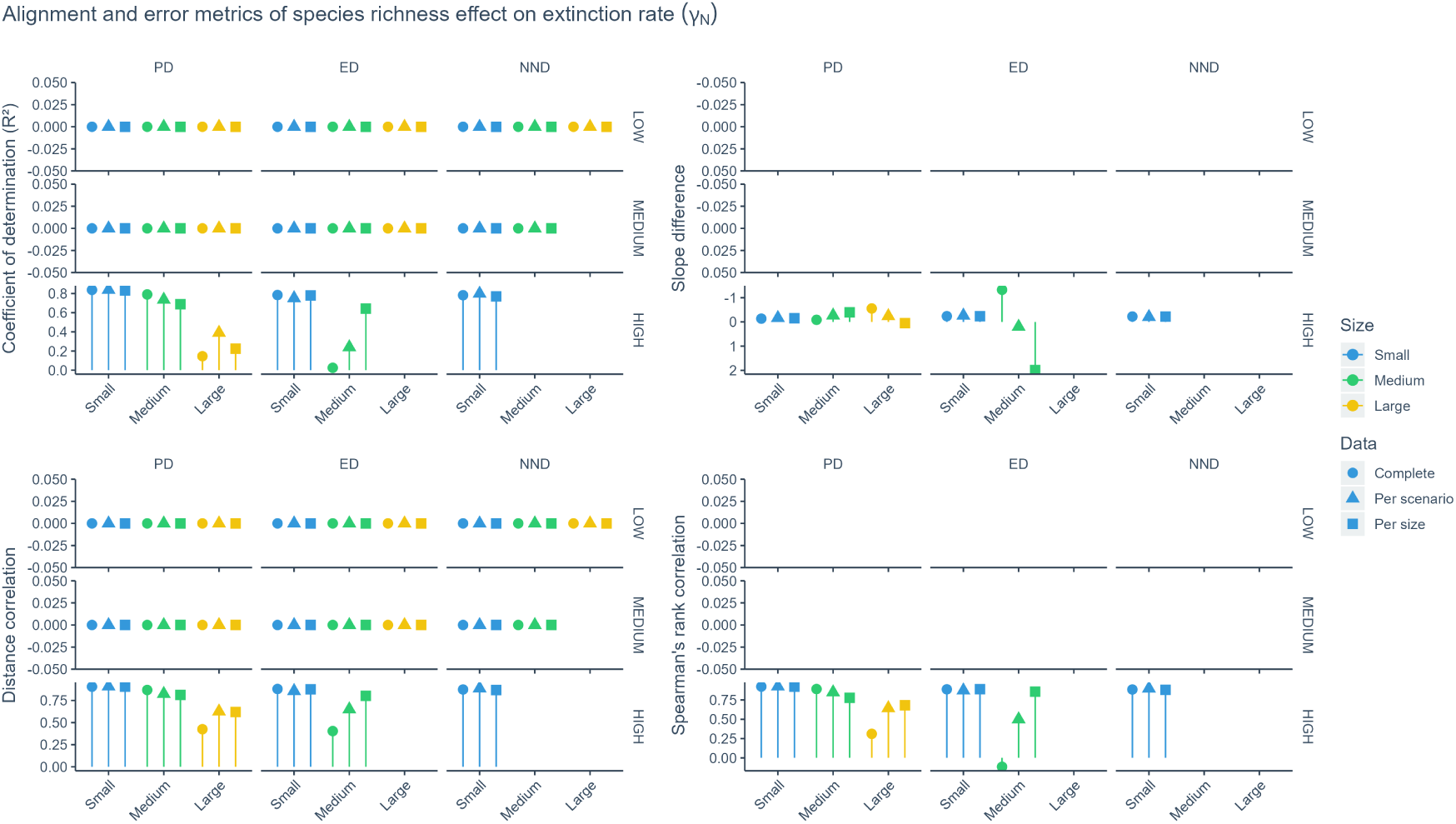
Comparison of the degree to which neural network regression estimates align with the conditional mean. There are four metrics, each describing a facet of the alignment, see Appendix I for details and interpretations. Within each panel, each facet column indicates results of a evolutionary scenario (PD for phylogenetic diversity, ED for evolutionary distinctiveness, NND for nearest neighbor distance); each facet row indicates results of a diversification model differing in complexity (as indicated by the number of parameters used to generate the trees). Yellow bars stand for metrics of small trees, green bars stand for medium trees and blue bars stand for large trees. At the top of the bars, shapes represent how the datasets were sliced to train the neural networks. Circles indicate complete datasets, triangles indicate slices by evolutionary scenario and squares indicate slices by tree size group. X-axis: size group of the trees. Y-axis: value of alignment metrics.

**Figure D.24:**
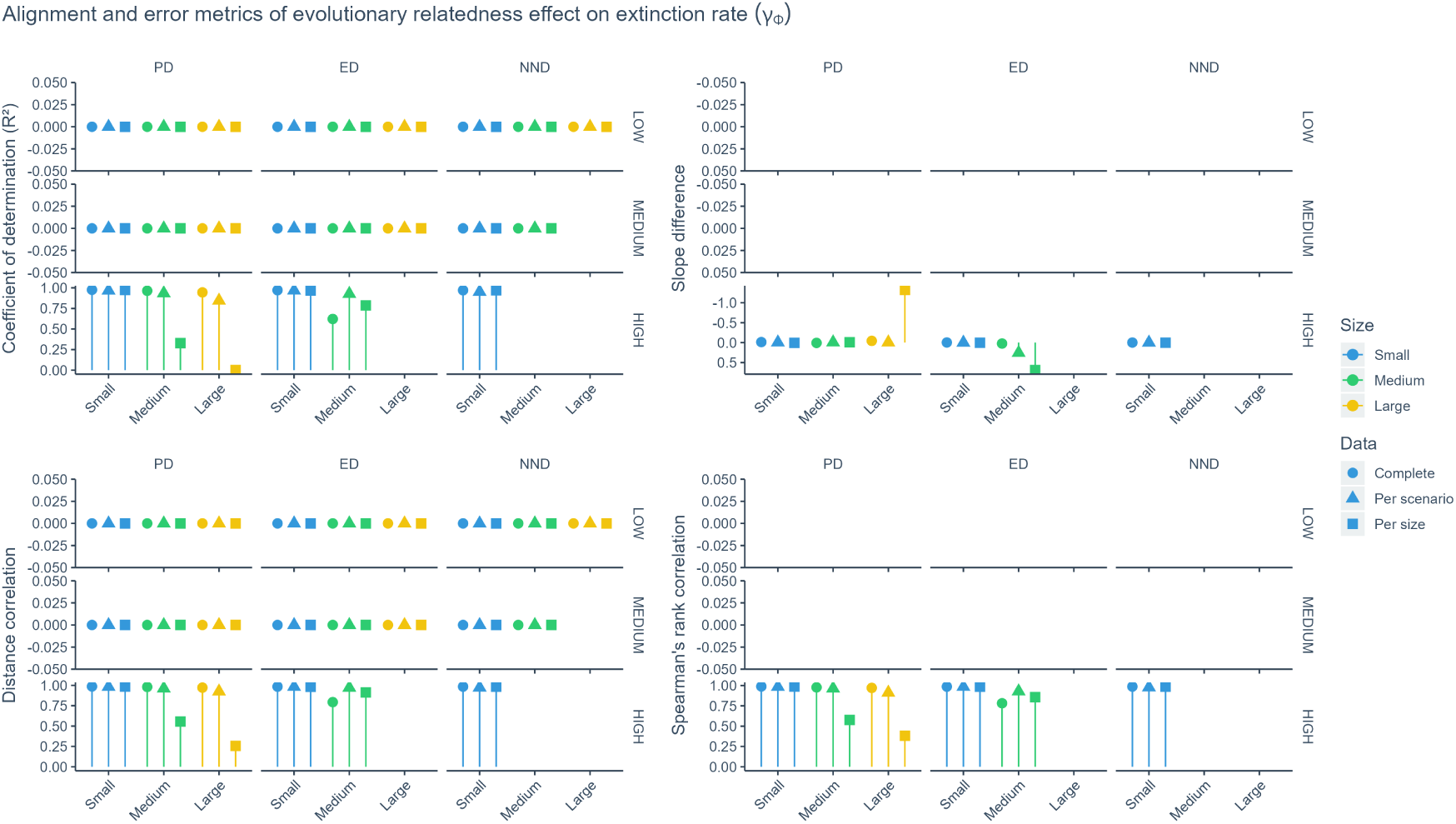
Comparison of the degree to which neural network regression estimates align with the conditional mean. There are four metrics, each describing a facet of the alignment, see Appendix I for details and interpretations. Within each panel, each facet column indicates results of a evolutionary scenario (PD for phylogenetic diversity, ED for evolutionary distinctiveness, NND for nearest neighbor distance); each facet row indicates results of a diversification model differing in complexity (as indicated by the number of parameters used to generate the trees). Yellow bars stand for metrics of small trees, green bars stand for medium trees and blue bars stand for large trees. At the top of the bars, shapes represent how the datasets were sliced to train the neural networks. Circles indicate complete datasets, triangles indicate slices by evolutionary scenario and squares indicate slices by tree size group. X-axis: size group of the trees. Y-axis: value of alignment metrics.

## Appendix E. Distribution of parameters

**Figure E.25:**
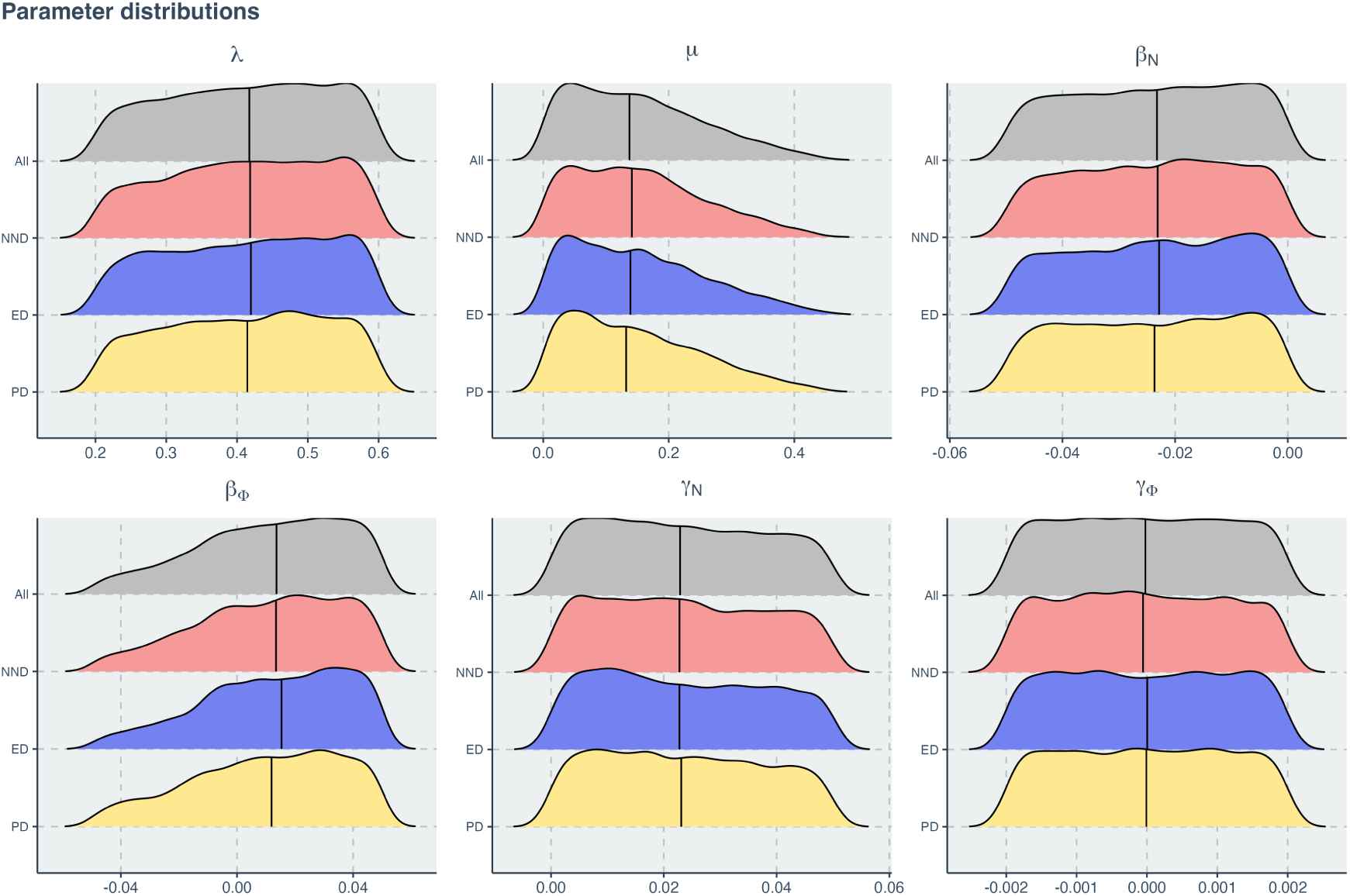
Generation parameter densities of phylogenetic trees in the training and validation dataset, before splitting. The densities of the trees in the test dataset are similar. Each panel displays the density of one tree generation parameter. Within panel, each ridge–line displays one angle view. The first ridge–line shows density of all trees. The second shows only NND (nearest-neighbor distance scenario) trees. The third shows only ED (evolutionary distinctiveness scenario) trees. The fourth shows only PD (phylogenetic diversity) trees.

## Appendix F. Effect Size and Tree Size

**Figure F.26:**
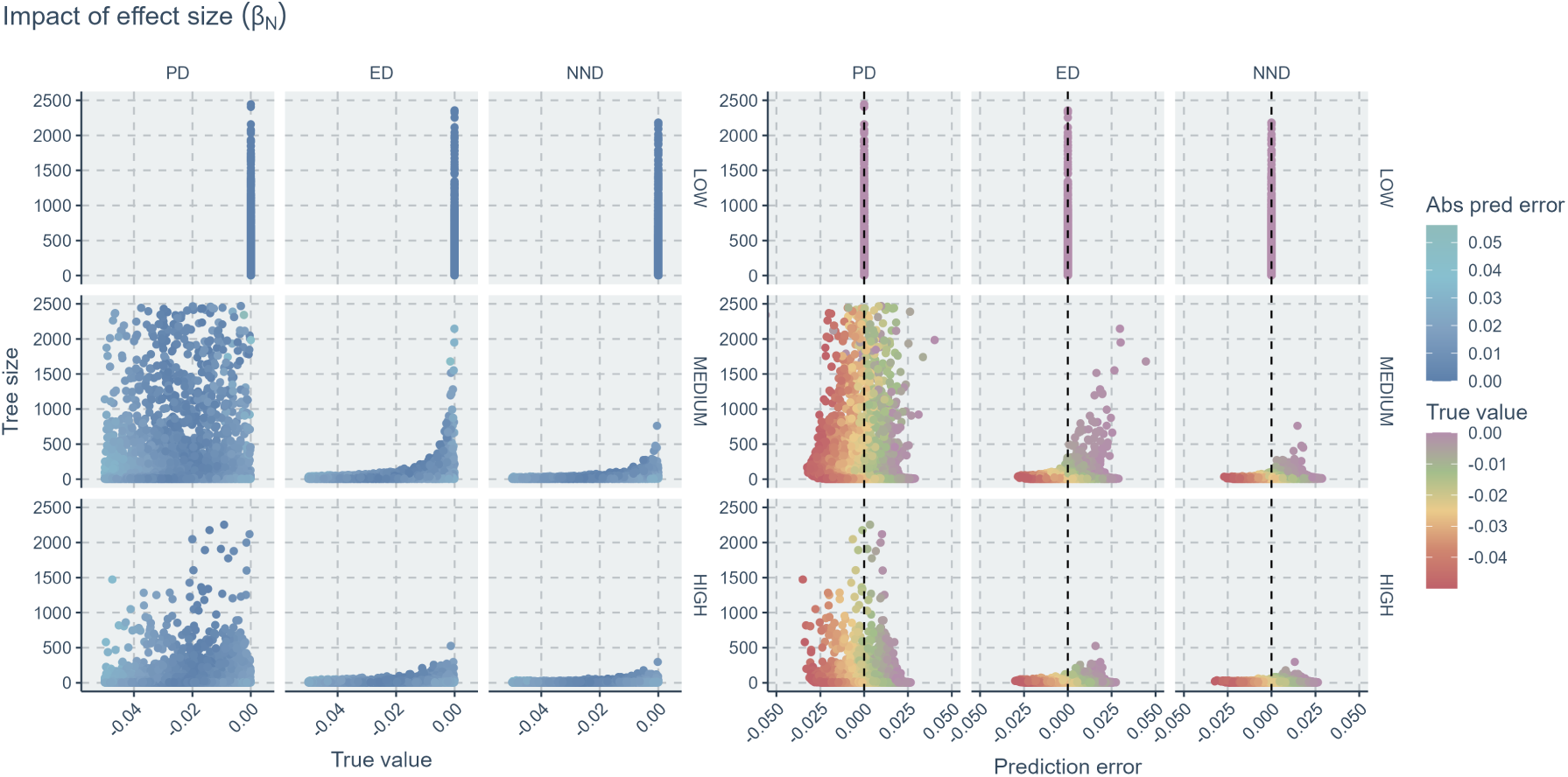
Relationships of tree size to true parameter values and prediction errors of the species richness effect size on speciation (*β_N_*). Within each panel, each column indicates results of an evolutionary scenario (PD for phylogenetic diversity, ED for evolutionary distinctiveness, NND for nearest neighbor distance); each row indicates results of a diversification model complexity (which indicates the number of parameters used to generate the trees). In the left panels, colors indicate the values of absolute prediction errors; as the blue points become darker, the errors become smaller thus the predictions more accurate. In the right panels, the colors indicate the values of true parameter. Points falling close to the vertical black dashed lines are accurate predictions (zero error). X-axis (left panel): true parameter values. X-axis (right panel): prediction error (absolute difference between true value and predicted value). Y-axis: tree size.

**Figure F.27:**
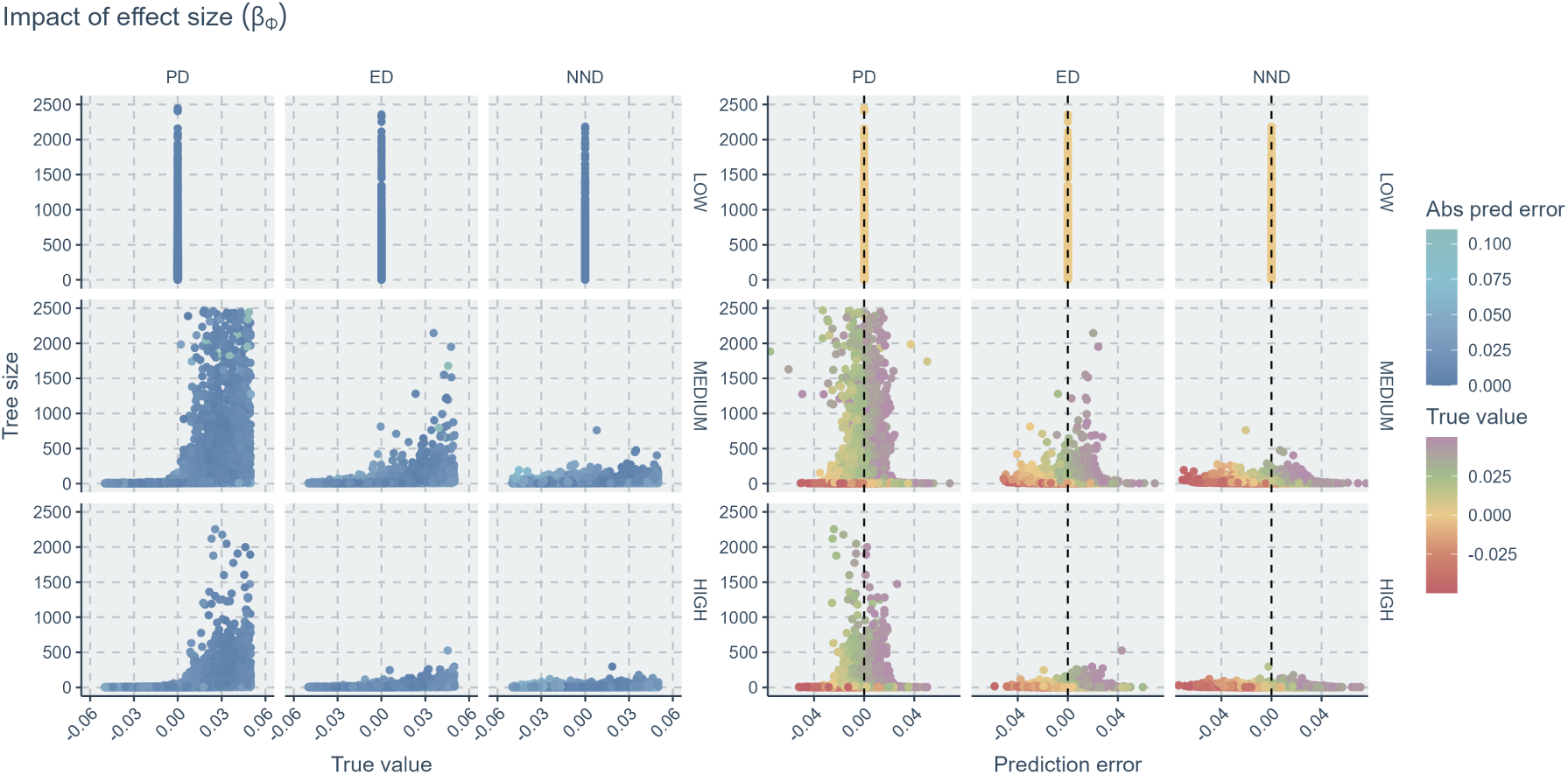
Relationships of tree size to true parameter values and prediction errors of the evolutionary relatedness effect size on speciation (*β_Φ_*). Within each panel, each column indicates results of an evolutionary scenario (PD for phylogenetic diversity, ED for evolutionary distinctiveness, NND for nearest neighbor distance); each row indicates results of a diversification model complexity (which indicates the number of parameters used to generate the trees). In the left panels, colors indicate the values of absolute prediction errors; as the blue points become darker, the errors become smaller thus the predictions more accurate. In the right panels, the colors indicate the values of true parameter. Points falling close to the vertical black dashed lines are accurate predictions (zero error). X-axis (left panel): true parameter values. X-axis (right panel): prediction error (absolute difference between true value and predicted value). Y-axis: tree size.

**Figure F.28:**
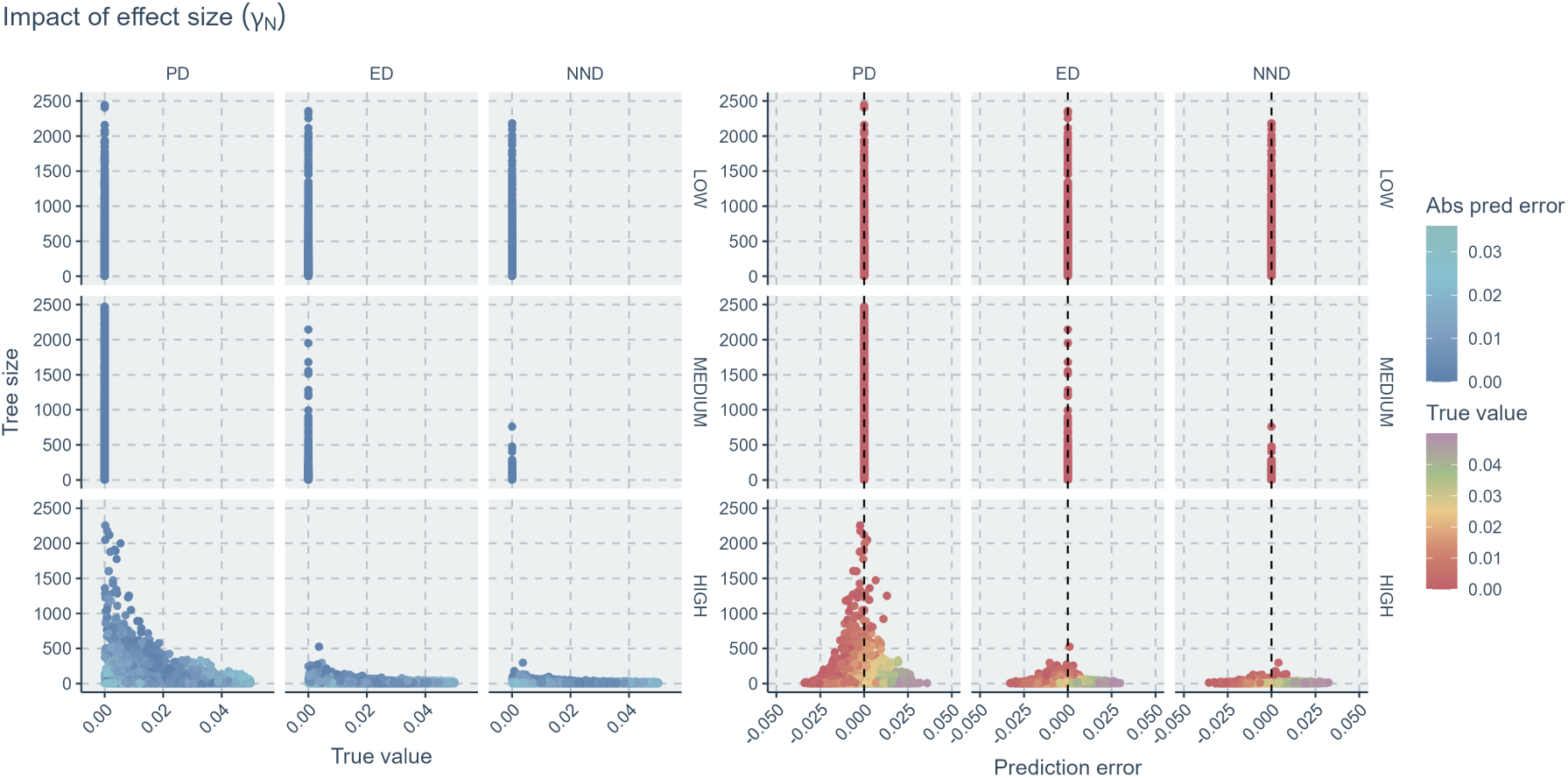
Relationships of tree size to true parameter values and prediction errors of the species richness effect size on extinction (*γ_N_*). Within each panel, each column indicates results of an evolutionary scenario (PD for phylogenetic diversity, ED for evolutionary distinctiveness, NND for nearest neighbor distance); each row indicates results of a diversification model complexity (which indicates the number of parameters used to generate the trees). In the left panels, colors indicate the values of absolute prediction errors; as the blue points become darker, the errors become smaller thus the predictions more accurate. In the right panels, the colors indicate the values of true parameter. Points falling close to the vertical black dashed lines are accurate predictions (zero error). X-axis (left panel): true parameter values. X-axis (right panel): prediction error (absolute difference between true value and predicted value). Y-axis: tree size.

**Figure F.29:**
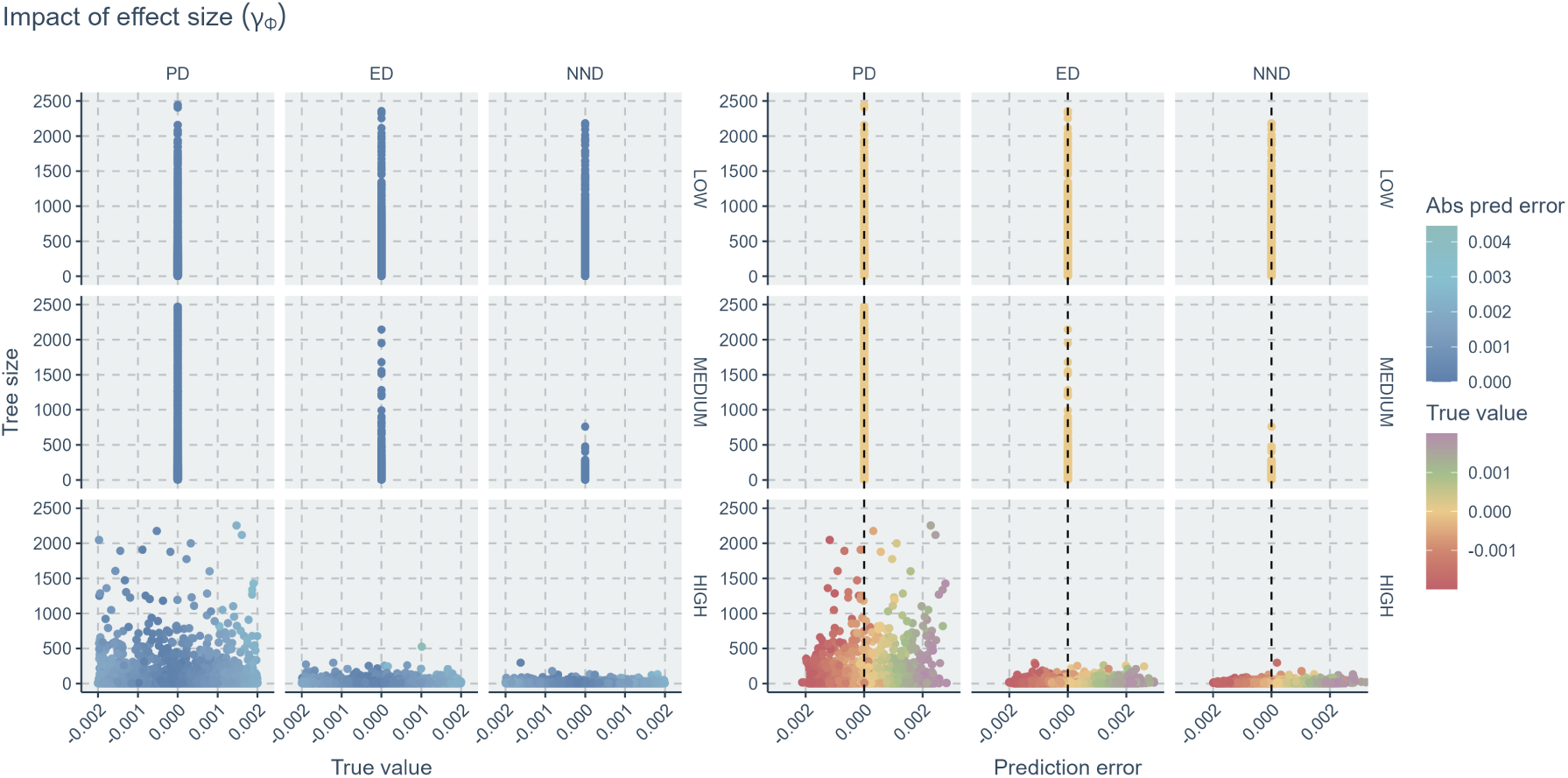
Relationships of tree size to true parameter values and prediction errors of the evolutionary relatedness effect size on extinction (*γ_Φ_*). Within each panel, each column indicates results of an evolutionary scenario (PD for phylogenetic diversity, ED for evolutionary distinctiveness, NND for nearest neighbor distance); each row indicates results of a diversification model complexity (which indicates the number of parameters used to generate the trees). In the left panels, colors indicate the values of absolute prediction errors; as the blue points become darker, the errors become smaller thus the predictions more accurate. In the right panels, the colors indicate the values of true parameter. Points falling close to the vertical black dashed lines are accurate predictions (zero error). X-axis (left panel): true parameter values. X-axis (right panel): prediction error (absolute difference between true value and predicted value). Y-axis: tree size.

## Appendix G. Regressor Misspecification

**Figure G.30:**
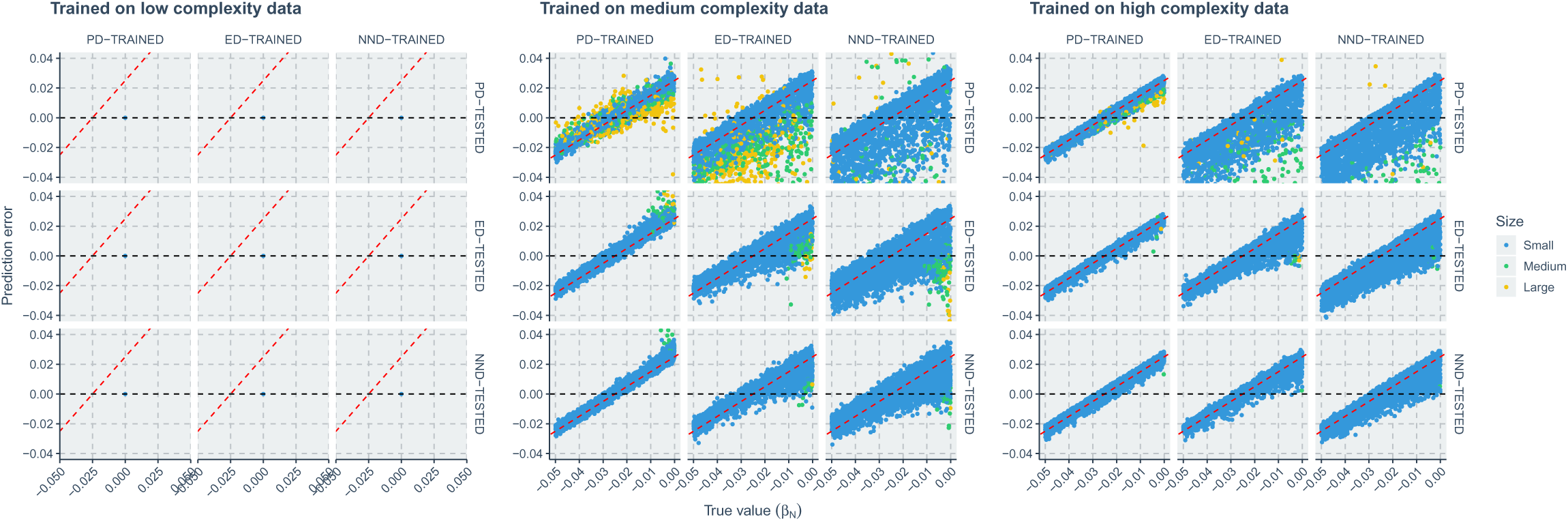
Comparison of neural network regression errors under misspecification. By misspecification, neural networks trained on trees under different evolutionary scenarios are used to estimate parameters on all trees, including those having different (misspecified) evolutionary scenarios. The three panels show the prediction error of three model complexity levels (which indicate the number of parameters used to generate the trees). Within each panel, each column indicates results of using neural networks trained on 3 evolutionary scenarios (see column facet strips, e.g. PD-TRAINED represents neural networks trained on trees generated under the phylogenetic diversity scenario) to estimate parameters on tree dataset generated under a particular scenario (see row facet strips, e.g. PD-TESTED indicate that neural network predictions were made on PD trees). Yellow points stand for results of small trees, green points stand for medium trees and blue points stand for large trees. X-axis: true value of the species richness effect size on speciation (*β_N_*). Y-axis: prediction error (absolute difference between true value and predicted value).

**Figure G.31:**
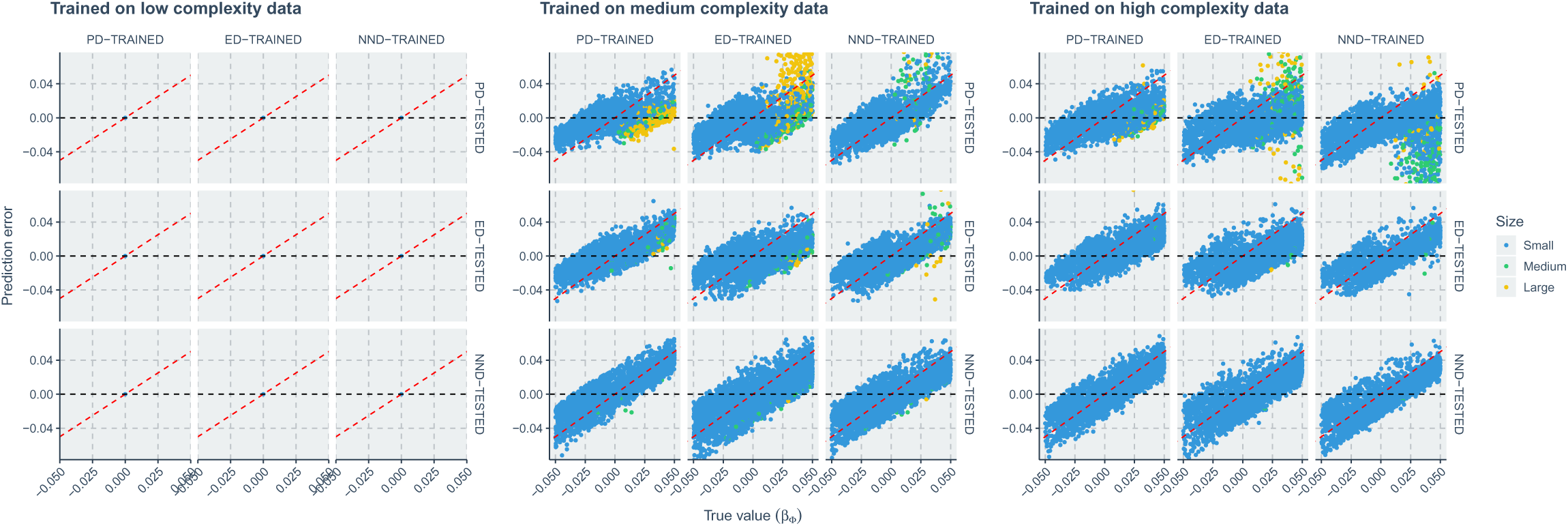
Comparison of neural network regression errors under misspecification. By misspecification, neural networks trained on trees under different evolutionary scenarios are used to estimate parameters on all trees, including those having different (misspecified) evolutionary scenarios. The three panels show the prediction error of three model complexity levels (which indicate the number of parameters used to generate the trees). Within each panel, each column indicates results of using neural networks trained on 3 evolutionary scenarios (see column facet strips, e.g. PD-TRAINED represents neural networks trained on trees generated under the phylogenetic diversity scenario) to estimate parameters on tree dataset generated under a particular scenario (see row facet strips, e.g. PD-TESTED indicate that neural network predictions were made on PD trees). Yellow points stand for results of small trees, green points stand for medium trees and blue points stand for large trees. X-axis: true value of the evolutionary relatedness effect size on speciation (*β_Φ_*). Y-axis: prediction error (absolute difference between true value and predicted value).

**Figure G.32:**
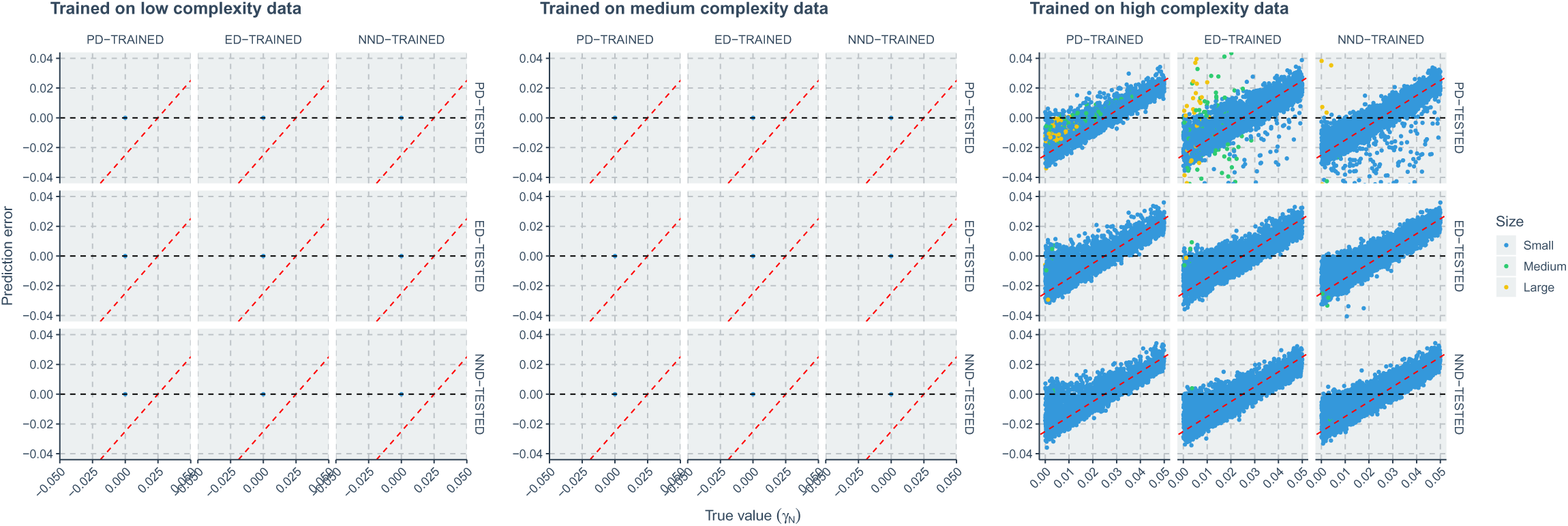
Comparison of neural network regression errors under misspecification. By misspecification, neural networks trained on trees under different evolutionary scenarios are used to estimate parameters on all trees, including those having different (misspecified) evolutionary scenarios. The three panels show the prediction error of three model complexity levels (which indicate the number of parameters used to generate the trees). Within each panel, each column indicates results of using neural networks trained on 3 evolutionary scenarios (see column facet strips, e.g. PD-TRAINED represents neural networks trained on trees generated under the phylogenetic diversity scenario) to estimate parameters on tree dataset generated under a particular scenario (see row facet strips, e.g. PD-TESTED indicate that neural network predictions were made on PD trees). Yellow points stand for results of small trees, green points stand for medium trees and blue points stand for large trees. X-axis: true value of the species richness effect size on extinction (*γ_N_*). Y-axis: prediction error (absolute difference between true value and predicted value).

**Figure G.33:**
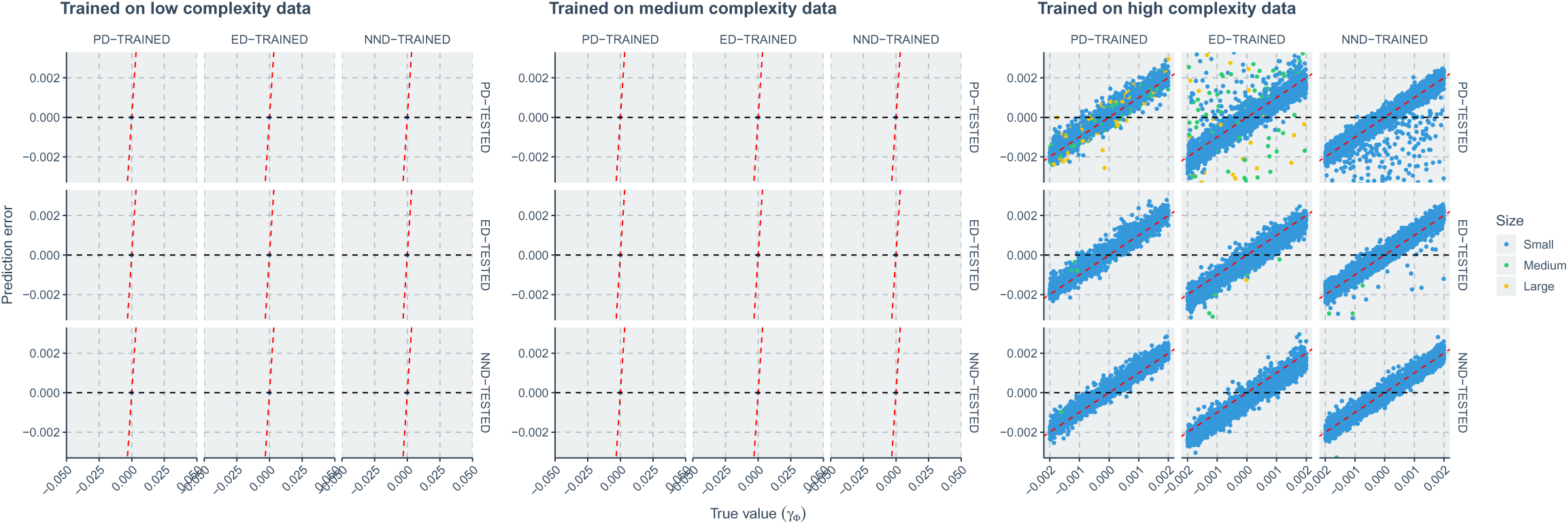
Comparison of neural network regression errors under misspecification. By misspecification, neural networks trained on trees under different evolutionary scenarios are used to estimate parameters on all trees, including those having different (misspecified) evolutionary scenarios. The three panels show the prediction error of three model complexity levels (which indicate the number of parameters used to generate the trees). Within each panel, each column indicates results of using neural networks trained on 3 evolutionary scenarios (see column facet strips, e.g. PD-TRAINED represents neural networks trained on trees generated under the phylogenetic diversity scenario) to estimate parameters on tree dataset generated under a particular scenario (see row facet strips, e.g. PD-TESTED indicate that neural network predictions were made on PD trees). Yellow points stand for results of small trees, green points stand for medium trees and blue points stand for large trees. X-axis: true value of the evolutionary relatedness effect size on extinction (*γ_Φ_*). Y-axis: prediction error (absolute difference between true value and predicted value).

## Appendix H. Details of Classification Performance Metrics

This appendix summarizes the notation and formulas used for the classification metrics reported in the main text. Let *y_i_* ∈ {PD, ED, NND} denote the true class of tree *i*, and 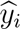 the predicted class (with the largest probability) returned by a classifier. For a given target class *k*, we define the usual entries of the 2 × 2 contingency table as

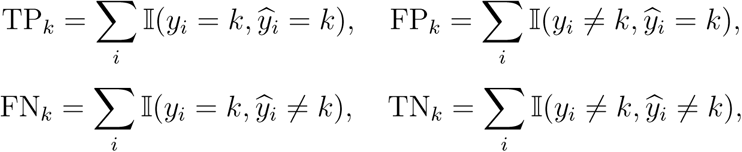

where 𝕀(·) denotes the indicator function. The class-wise precision, recall, and F1-score are then given by

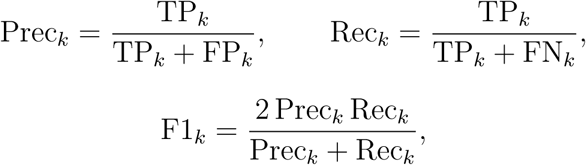

with the convention that ratios with zero denominators are treated as undefined and omitted from macro-averages. Overall (micro-averaged) accuracy is defined as

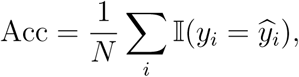

where *N* is the total number of test trees. Unless stated otherwise, we report macro-averaged precision, recall, and F1-score obtained by taking the unweighted arithmetic mean of the class-wise scores.

For confusion matrices, we index rows by the true class *r* and columns by the predicted class *c*. The raw counts are *C_rc_* = Σ_*i*_𝕀(*y_i_* = *r,* 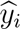 = *c*), and the row-normalized entries

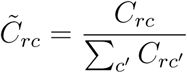

sum to one within each row. Hence, *C̃_rr_* equals the class-wise recall for class *r*, and *C̃_rc_* for *c* ≠ *r* equals the conditional probability that a tree from class *r* is misclassified as class *c*.

When comparing two methods *A* and *B*, we summarize differences in recall patterns using

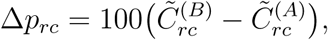

the change in row-normalized recall (in percentage points) for cell (*r, c*). Positive values indicate that method *B* predicts class *c* more often than method *A* for trees with true class *r*, and negative values indicate the reverse.

For the calibration analysis, we follow the binning scheme described in Section 2.3.2. The reliability diagram is obtained by plotting conf(*B_m_*) against acc(*B_m_*) for each bin *m*, together with the diagonal line acc = conf. The expected calibration error (ECE) is computed as

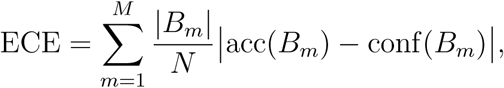

where *N* is the total number of test trees in the stratum under consideration. Lower ECE values correspond to better-calibrated predictive probabilities.

## Appendix I. Definitions and Interpretations of the Alignment Metrics

Several metrics are computed to assess the extent to which the neural network predictions align with, or deviate from, merely reporting the conditional mean.

Let *x* denote the true value of the quantity under investigation and let

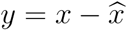

represent the prediction error, where *x̂* is the neural network prediction. In each subgroup, a linear model is fitted via

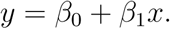

The coefficient of determination, denoted by *R*^2^, is used as a metric of prediction alignment to the conditional mean. It is defined as

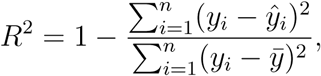

where *ŷ_i_* are the fitted values from the regression and *y̅* is the mean of the observed *y* values. A high *R*^2^ indicates that a large proportion of the variability in the prediction error is explained by the true values, suggesting that the neural network predictions are largely driven by the conditional mean relationship. A low *R*^2^ implies that the neural network is capturing additional patterns beyond this baseline.

The slope difference is defined as

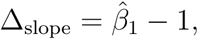

where 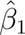 is the estimated slope from the regression. When predictions are simply the conditional mean, one would expect a near-unit slope; thus, a slope difference close to zero indicates strong alignment with the conditional mean. A large absolute value of Δ_slope_ signifies that the fitted relationship deviates substantially from a unit slope, implying that the neural network is capturing additional signal beyond the conditional mean.

Distance correlation is employed to quantify the overall dependence between *x* and *y* without assuming any specific functional form. It is defined as dCov(*x, y*)

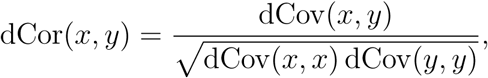

where dCov denotes the distance covariance. A high distance correlation indicates a strong association between the true values and the prediction error, implying a scenario where the predictions are largely driven by the conditional mean. A low distance correlation, on the other hand, suggests that the neural network is exploiting additional information.

Spearman’s rank correlation coefficient provides a robust, nonparametric measure of the monotonic association between *x* and *y*. When the network simply reports the conditional mean, the prediction error *y* should vary monotonically with *x* (for example, as *y* = *x* − *c* for some constant *c*), resulting in a high absolute Spearman correlation (close to 1). A Spearman correlation near zero, however, indicates that the prediction error does not follow a monotonic trend with respect to *x*; this lack of an association implies that the network’s predictions capture additional patterns or signals beyond the conditional mean.

## Appendix J. Parameter and Tree Size Slices

**Table J.2:** Parameter and tree size ranges used for slicing classification performance results.

| Parameter or Size | Ranges |
| --- | --- |
| $\lambda_0$ | $[0.2, 0.3), [0.3, 0.4), [0.4, 0.5), [0.5, 0.6)$ |
| $\mu_0$ | $[0.00, 0.12), [0.12, 0.24), [0.24, 0.36), [0.36, 0.48)$ |
| $\beta_N$ | $[-0.05, -0.04), [-0.04, -0.03), [-0.03, -0.02), [-0.02, -0.01), [-0.01, 0.00)$ |
| $\beta_\Phi$ | $[-0.05, -0.04), [-0.04, -0.03), [-0.03, -0.02), [-0.02, -0.01), [-0.01, 0.00), [0.00, 0.01),$<br>$[0.01, 0.02), [0.02, 0.03), [0.03, 0.04), [0.04, 0.05)$ |
| $\gamma_N$ | $[0.00, 0.01), [0.01, 0.02), [0.02, 0.03), [0.03, 0.04), [0.04, 0.05)$ |
| $\gamma_\Phi$ | $[-0.002, -0.001), [-0.001, 0.000), [0.000, 0.001), [0.001, 0.002)$ |
| Tree Size | $[0, 20), [20, 40), [40, 60), [60, 80), [80, 100), [100, 120), [120, 140), [140, 160), [160, +\infty)$ |

## Appendix K. Mean, variance, and an approximate confidence interval for a birth–death process

Consider a linear birth–death process {*N* (*t*)}*_t_*_≥0_ with *N* (0) = 1. Each lineage gives birth at rate *λ* and dies at rate *µ*, independently across lineages. Let *r* = *λ* − *µ*.

### Mean

Let *M* (*t*) = E[*N* (*t*)]. Conditional on *N* (*t*) = *n*, the total birth rate is *nλ* (increasing the count by 1) and the total death rate is *nµ* (decreasing the count by 1). Taking expectations yields

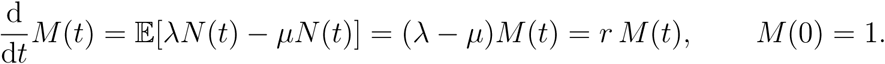

Hence,

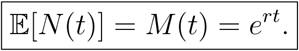

### Second moment and variance

Write *S*(*t*) = E[*N* (*t*)^2^]. Using the standard jump-moment calculation, conditional on *N* (*t*) = *n*,

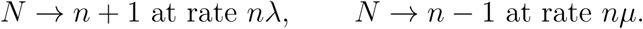

Therefore,

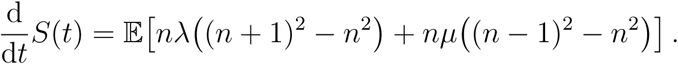

Since (*n* + 1)^2^ − *n*^2^ = 2*n* + 1 and (*n* − 1)^2^ − *n*^2^ = −2*n* + 1, we obtain

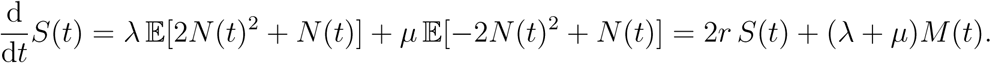

With *M* (*t*) = *e^rt^* and *S*(0) = E[*N* (0)^2^] = 1, we solve the linear ODE

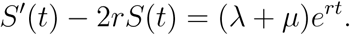

*Case λ* /= *µ (i.e. r* /= 0).. Using the integrating factor *e*^−2^*^rt^*,

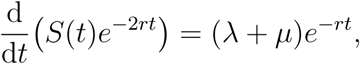

so

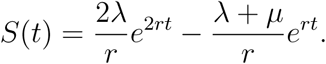

Hence,

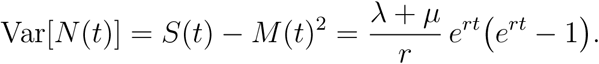

That is,

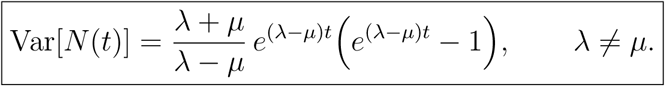

*Critical case λ* = *µ (i.e. r* = 0).. Then *M* (*t*) ≡ 1, and the moment equation reduces to *S*^t^(*t*) = 2*λM* (*t*) = 2*λ* with *S*(0) = 1, giving

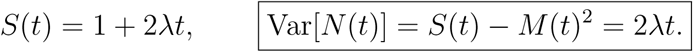

### Approximate (unconditional) normal confidence interval for N (t)

For sufficiently large expected counts (typically when *λ > µ* and *t* is not too small), one may use a crude normal approximation

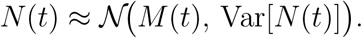

Let *z*_1−_*_α/_*_2_ denote the standard normal critical value (e.g. *z*_0.975_ ≈ 1.96). An approximate two-sided (1 − *α*) interval is

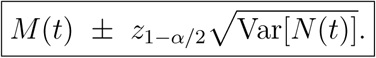

Because *N* (*t*) is discrete and can be highly skewed (especially when extinction probability is non-negligible), this interval should be interpreted cautiously; in practice one may also truncate the lower bound at 0.

## References

Angelopoulos, A.N., Bates, S., 2022.A gentle introduction to conformal predic-tion and distribution-free uncertainty quantification. doi:10.48550/arXiv.2107.07511. arXiv: 2107.07511 [cs] tex.copyright: arXiv.org perpetual, non-exclusive license.

Blomberg, S.P., Garland, T., Ives, A.R., 2003. Testing for phylogenetic signal in com-parative data: Behavioral traits are more labile. Evolution; international journal of organic evolution 57, 717–745. URL: https://academic.oup.com/evolut/article/57/4/717/6756141, doi:10.1111/j.0014-3820.2003.tb00285.x.

Blundell, C., Cornebise, J., Kavukcuoglu, K., Wierstra, D., 2015. Weight uncertainty in neural network, in: International conference on machine learning, PMLR. pp. 1613–1622. URL: https://proceedings.mlr.press/v37/blundell15.

Cole, D.J., 2019. Parameter redundancy and identifiability in hidden Markov models. METRON 77, 105–118. doi:10.1007/s40300-019-00156-3.

Etienne, R.S., Haegeman, B., Stadler, T., Aze, T., Pearson, P.N., Purvis, A., Phillimore, A.B., 2012. Diversity-dependence brings molecular phylogenies closer to agreement with the fossil record. Proceedings of the Royal Society B: Biological Sciences 279, 1300–1309. doi:10.1098/rspb.2011.1439.

Etienne, R.S., Morlon, H., Lambert, A., 2014. Estimating the duration of specia-tion from phylogenies. Evolution; international journal of organic evolution 68, 2430–2440. doi:10.1111/evo.12433.

Etienne, R.S., Rosindell, J., 2012. Prolonging the past counteracts the pull of the present: Protracted speciation can explain observed slowdowns in diversification. Systematic Biology 61, 204. doi:10.1093/sysbio/syr091.

Fey, M., Lenssen, J.E., 2019.Fast graph representation learning with PyTorch geo-metric. doi:10.48550/ARXIV.1903.02428. arXiv: 1903.02428v3 [cs.LG] tex.copy-right: arXiv.org perpetual, non-exclusive license.

Gal, Y., Ghahramani, Z., 2016. Dropout as a Bayesian approximation: Repre-senting model uncertainty in deep learning, in: Proceedings of The 33rd In-ternational Conference on Machine Learning, PMLR. pp. 1050–1059. URL: https://proceedings.mlr.press/v48/gal16.html.

Gernhard, T., 2008. The conditioned reconstructed process. Journal of the-oretical biology 253, 769–778. URL: https://www.sciencedirect.com/science/article/pii/S0022519308001811?casa_token=t5-lMXLukaYAAAAA:WGXnhgFTKSQuPQ6fx3FtWBqCBQawo8hZU9oaUjEe3dQFvNYbhatYMTIqAZLawOitkzAt8Zkg, doi:10.1016/j.jtbi.2008.04.005.

Guo, C., Pleiss, G., Sun, Y., Weinberger, K.Q., 2017. On calibration of modern neural networks, in: International conference on machine learning, PMLR. pp. 1321–1330. URL: http://proceedings.mlr.press/v70/guo17a.html.

Huber, P.J., 1992. Robust estimation of a location parameter, in: Breakthroughs in Statistics. Springer New York, New York, NY, pp. 492–518. doi:10.1007/978-1-4612-4380-9_35. iSSN: 0172-7397.

Imambi, S., Prakash, K.B., Kanagachidambaresan, G.R., 2021. PyTorch, in: Prakash, K.B., Kanagachidambaresan, G.R. (Eds.), Programming with Ten-sorFlow. Springer International Publishing, Cham, pp. 87–104. doi:10.1007/978-3-030-57077-4_10. iSSN: 2522-8609.

Kipf, T.N., Welling, M., 2016.Semi-supervised classification with graph convolu-tional networks. doi:10.48550/ARXIV.1609.02907. arXiv: 1609.02907v4 [cs.LG] tex.copyright: arXiv.org perpetual, non-exclusive license.

Kondratyeva, A., Grandcolas, P., Pavoine, S., 2019. Reconciling the concepts and measures of diversity, rarity and originality in ecology and evolution. Biological Reviews 94, 1317–1337. doi:10.1111/brv.12504.

Kubo, T., Iwasa, Y., 1995. Inferring the rates of branching and extinction from molecular phylogenies. Evolution; international journal of organic evolution 49, 694–704. doi:10.2307/2410323.

Lajaaiti, I., Lambert, S., Voznica, J., Morlon, H., Hartig, F., 2023. A compari-son of deep learning architectures for inferring parameters of diversification mod-els from extant phylogenies. doi:10.1101/2023.03.03.530992. tex.copyright: © 2023, Posted by Cold Spring Harbor Laboratory. This pre-print is available under a Creative Commons License (Attribution-NoDerivs 4.0 International), CC BY-ND 4.0, as described at http://creativecommons.org/licenses/by-nd/4.0/.

Lakshminarayanan, B., Pritzel, A., Blundell, C., 2017. Simple and scal-able predictive uncertainty estimation using deep ensembles, in: Ad-vances in Neural Information Processing Systems, Curran Associates, Inc. URL: https://proceedings.neurips.cc/paper_files/paper/2017/hash/9ef2ed4b7fd2c810847ffa5fa85bce38-Abstract.html.

Lambert, S., Voznica, J., Morlon, H., 2023. Deep learning from phyloge-nies for diversification analyses. Systematic Biology 72, 1262–1279. URL: DifferentialEvolution, doi:10.1093/sysbio/syad044.

Lei, J., G’Sell, M., Rinaldo, A., Tibshirani, R.J., Wasserman, L., 2018. Distribution-free predictive inference for regression. Journal of the American Statistical Asso-ciation 113, 1094–1111. doi:10.1080/01621459.2017.1307116.

Loshchilov, I., Hutter, F., 2017. Decoupled weight decay regularization. doi:10.48550/ARXIV.1711.05101. arXiv: 1711.05101v3 [cs.LG] tex.copyright: arXiv.org perpetual, non-exclusive license.

Louca, S., Pennell, M.W., 2020. Extant timetrees are consistent with a myriad of diversification histories. Nature 580, 502–505. URL: https://www.nature.com/articles/s41586-020-2176-1, doi:10.1038/s41586-020-2176-1.

Luebke, D., 2008. CUDA: Scalable parallel programming for high-performance scien-tific computing, in: 2008 5th IEEE international symposium on biomedical imag-ing: from nano to macro, IEEE. pp. 836–838. URL: https://ieeexplore.ieee.org/abstract/document/4541126/, doi:10.1109/isbi.2008.4541126.

Moen, D., Morlon, H., 2014. Why does diversification slow down? Trends in Ecology and Evolution 29, 190–197. doi:10.1016/j.tree.2014.01.010.

Nee, S., Holmes, E.C., May, R.M., Harvey, P.H., 1994a. Extinction rates can be estimated from molecular phylogenies. Philosophical Transactions of the Royal Society of London. Series B: Biological Sciences 344, 77–82. doi:10.1098/rstb.1994.0054.

Nee, S., May, R.M., Harvey, P.H., 1994b. The reconstructed evolutionary process. Philosophical Transactions of the Royal Society of London. Series B: Biological Sciences 344, 305–311. doi:10.1098/rstb.1994.0068.

Nee, S., Mooers, A.O., Harvey, P.H., 1992. Tempo and mode of evolution revealed from molecular phylogenies. Proceedings of the National Academy of Sciences, USA 89, 8322–8326. doi:10.1073/pnas.89.17.8322.

Niculescu-Mizil, A., Caruana, R., 2005. Predicting good probabilities with su-pervised learning, in: Proceedings of the 22nd international conference on Ma-chine learning - ICML ’05, ACM Press, Bonn, Germany. pp. 625–632. URL: http://portal.acm.org/citation.cfm?doid=1102351.1102430, doi:10.1145/1102351.1102430.

Pannetier, T., Martinez, C., Bunnefeld, L., Etienne, R.S., 2021. Branching patterns in phylogenies cannot distinguish diversity-dependent diversification from time-dependent diversification. Evolution; international journal of organic evolution 75, 25–38. doi:10.1111/evo.14124.

Phillimore, A.B., Price, T.D., 2008. Density-dependent cladogenesis in birds. PLoS Biology 6, e71. doi:10.1371/journal.pbio.0060071.

Qin, T., 2023. eveGNN: Codebase for phylogenetic tree parameter estimation with neural networks. URL: https://github.com/EvoLandEco/eveGNN.tex.copyright: MIT.

Qin, T., van Benthem, K.J., Valente, L., Etienne, R.S., 2025a. Parameter estimation from phylogenetic trees using neural networks and ensemble learning. Systematic Biology, syaf060doi:10.1093/sysbio/syaf060.

Qin, T., Valente, L., Etienne, R.S., 2025b. Impact of evolutionary related-ness on species diversification and tree shape. Journal of Theoretical Biology 598, 111992. URL: https://www.sciencedirect.com/science/article/pii/S0022519324002777, doi:10.1016/j.jtbi.2024.111992.

Quental, T.B., Marshall, C.R., 2010. Diversity dynamics: molecular phylogenies need the fossil record. Trends in Ecology and Evolution 25, 434–441. doi:10.1016/j.tree.2010.05.002.

R Core Team, 2025.R: A language and environment for statistical computing.

Rosindell, J., Cornell, S.J., Hubbell, S.P., Etienne, R.S., 2010. Protracted speciation revitalizes the neutral theory of biodiversity. Ecology Letters 13, 716–727. URL: https://onlinelibrary.wiley.com/doi/pdf/10.1111/j.1461-0248.2010.01463.x, doi:10.1111/j.1461-0248.2010.01463.x.

Sak, H., Senior, A., Beaufays, F., 2014.Long short-term memory based recur-rent neural network architectures for large vocabulary speech recognition. doi:10.48550/ARXIV.1402.1128. arXiv: 1402.1128v1 [cs.NE] tex.copyright: arXiv.org perpetual, non-exclusive license.

Shafer, G., Vovk, V., 2008. A tutorial on conformal prediction. Journal of Ma-chine Learning Research 9. URL: https://www.jmlr.org/papers/volume9/shafer08a/shafer08a.pdf?trk=public_post_comment-text.

Srivastava, D.S., Cadotte, M.W., Macdonald, A.A.M., Marushia, R.G., Mirotchnick, N., 2012. Phylogenetic diversity and the functioning of ecosystems. Ecology Letters 15, 637–648. doi:10.1111/j.1461-0248.2012.01795.x.

Steel, M., McKenzie, A., 2001. Properties of phylogenetic trees generated by Yule-type speciation models. Mathematical biosciences 170, 91–112. URL: https://www.sciencedirect.com/science/article/pii/S0025556400000614?casa_token=gq7LCYng5_wAAAAA:QH_IO7wCJdIeGALC3y8eT5VAbcR-nx28rgs_SOgjKPbXMqjyNkx1bnZsJMZY29pE7ulVrLnx, doi:10.1016/s0025-5564(00)00061-4.

Townsend, J.P., Su, Z., Tekle, Y.I., 2012. Phylogenetic signal and noise: Predicting the power of a data set to resolve phylogeny. Systematic Biology 61, 835. doi:10.1093/sysbio/sys036.

Voznica, J., Zhukova, A., Boskova, V., Saulnier, E., Lemoine, F., Moslonka-Lefebvre, M., Gascuel, O., 2022. Deep learning from phylogenies to uncover the epidemio-logical dynamics of outbreaks. Nature Communications 13, 3896. URL: https://www-nature-com.proxy-ub.rug.nl/articles/s41467-022-31511-0, doi:10.1038/s41467-022-31511-0.

Ying, Z., You, J., Morris, C., Ren, X., Hamilton, W., Leskovec, J., 2018. Hierarchical graph representation learning with differentiable pooling, in: Advances in Neural Information Processing Systems, Curran Associates, Inc. URL: https://proceedings.neurips.cc/paper_files/paper/2018/hash/e77dbaf6759253c7c6d0efc5690369c7-Abstract.html.

Zadrozny, B., Elkan, C., 2002. Transforming classifier scores into accurate multiclass probability estimates, in: Proceedings of the eighth ACM SIGKDD international conference on Knowledge discovery and data mining, ACM, Edmonton Alberta Canada. pp. 694–699. doi:10.1145/775047.775151.

